# RUNX1 controls the dynamics of cell cycle entry of naïve resting B cells by regulating expression of cell cycle and immunomodulatory genes in response to BCR stimulation

**DOI:** 10.1101/2020.12.01.406744

**Authors:** Inesa Thomsen, Natalia Kunowska, Roshni de Souza, Anne-Marie Moody, Greg Crawford, Yi-Fang Wang, Sanjay Khadayate, Jessica Strid, Mohammad M. Karimi, Alexis Barr, Niall Dillon, Pierangela Sabbattini

## Abstract

RUNX1 is a transcription factor that plays key roles in haematopoietic development and in adult haematopoiesis and lymphopoiesis. Here we report that RUNX1 is also involved in controlling the dynamics of cell cycle entry of naïve resting B cells in response to stimulation of the B cell receptor (BCR). Conditional knockout of *Runx1* in mouse resting B cells resulted in accelerated entry of the cells into S-phase following BCR engagement. Our results indicate that Runx1 regulates the cyclin D2 (*Ccnd2*) gene, the immediate early genes, *Fosl2*, *Atf3* and *Egr2*, and the Notch effector *Rbpj*, in B cells, reducing the rate at which transcription of these genes increases following BCR stimulation. RUNX1 interacts with the chromatin remodeller SRCAP, recruiting it to promoter and enhancer regions of the *Ccnd2* gene. BCR-mediated activation triggers switching between binding of RUNX1 and its paralog RUNX3 and between SRCAP and the SWI/SNF remodelling complex member BRG1. We also find that RUNX1 regulates expression of a number of immunomodulatory genes in resting B cells. These include the interferon receptor subunit gene *Ifnar1*, which is upregulated in B cells from lupus patients, the *Ptpn22* gene, which has been identified as a major lupus risk allele, and the *Lrrk2* gene, which is mutated in familial Parkinson’s disease. The hyperresponsiveness of the *Runx1* knockout B cells to antigen stimulation and its role in regulating a suite of genes that are known to be associated with autoimmune disease suggest that RUNX1 is a major regulator of B cell tolerance and autoimmunity.

## INTRODUCTION

RUNX1 is a pioneer factor, which is involved in the regulation of haematopoiesis from a very early stage in mammalian development and is required for fetal liver hematopoiesis and maintenance of the correct balance of adult hematopoietic stem cell compartment formation (1–4). In addition to its roles in haematopoiesis, RUNX1 also has critical roles in the early stages of T and B cell development (3, 5). RUNX1 belongs to the runt-related family of transcription factors, which has three members, RUNX1, RUNX2 and RUNX3. The runt-related factors act in conjunction with the non-DNA binding factor, CBFβ, to regulate gene expression in development, cell differentiation and cancer (reviewed in (2)).

The biological effects of RUNX1 are highly context dependent and it has been shown to function either as a transcriptional activator or as a repressor through interactions with different co-factors (6). This diversity of function is also reflected in the fact that RUNX1 can act as a tumour suppressor or as an oncogene in different cell types (7). The *RUNX1* gene is involved in multiple translocations that give rise to oncogenic fusion proteins (8). These include the *RUNX1*-ETO fusion, which is the most common cytogenetic abnormality in acute myeloid leukemia (9), and the ETV6-*RUNX1* fusion, which occurs in approximately 25% of cases of childhood precursor-B cell acute lymphoblastic leukemia (pre-B ALL) (10, 11).

During early B cell development, RUNX1 acts in conjunction with E2A and early B cell factor (EBF) to activate B cell-specific gene expression at the pre-pro-B cell stage (12, 13), and conditional knockout of *Runx1* in pro-B cells results in a block in the transition from the pro- to the pre-B cell stage (13). In human mature B cells, analysis of the functions of RUNX1 and RUNX3 in Epstein-Barr virus (EBV) transformed B cells has shown that the EBV transcription factor, EBNA2, enhances expression of RUNX3 resulting in RUNX3-mediated downregulation of *RUNX1* expression (14). Human RUNX1 was also shown to have a growth inhibitory effect on EBV-transformed cells, which was not observed for mouse RUNX1 due to the absence of a specific N-terminal segment from the mouse protein (15). Despite these results indicating a role for RUNX1 in regulating B cell proliferation, there is very little information about the mechanisms by which this might occur or how RUNX1 affects B cell activation in response to antigenic stimulation of the B cell receptor (BCR).

Here we show that RUNX1 acts in conjunction with the chromatin remodelling complex SRCAP to regulate the timing of entry of mouse resting B cells into S-phase. RUNX1 binds to promoters and distally-located elements at key cell cycle and immediate early genes that have a poised epigenetic configuration. Conditional knockout of the *Runx1* gene in resting B cells results in deregulation of Cyclin D2 and immediate early gene expression and accelerated entry into S-phase in response to BCR stimulation. We also show that RUNX1 is responsible for correct regulation of a number of genes that have been shown to be risk factors in autoimmune disease.

## Material and Methods

### Mice

The Runx1 c-k/o / Cd23-Cre conditional knockout mice were generated by crossing the Runx1^fl/fl^ mouse line (generously provided by Nancy Speck, University of Pennsylvania) (16) with the Cd23-Cre transgenic mouse line (generously provided by Meinrad Busslinger, IMP Vienna) (17). The CD23-cre transgene is first expressed expressed at the transitional stage, resulting in conditional kncockout of floxed target genes in mature resting B cells (17). All work involving mice was carried out under the regulations of the British Home Office and was approved by the Imperial College Animal Welfare and Ethical Review Body.

### Spleen and lymph node isolation and splenic resting B cell purification

Spleens and cervical lymph nodes were isolated from 6-10 weeks old C57B6 mice and homogenized through a sieve in B cell culture medium (RPMI-1640 (Lonza), 10% fetal calf serum (FCS) (Sigma), 0.1 U/ml penicillin (Lonza), 0.1 μg/ml streptomycin (Lonza), 2mM L-Glutamine (Lonza), 50μM beta-mercapthoethanol (Gibco)). The cell suspension was centrifuged on a Ficoll-Paque (GE Healthcare) cushion and the buffy coat layer was resuspended at a concentration of 1× 10^8^ cells/ml in PBS + 2% FCS/ 1mM EDTA. Resting B cells were isolated using the EasySep Negative Selection- Mouse B Cell Isolation Kit (STEMCELL Technologies), which depletes for non-B cell markers and activated CD43^+^ B cells. For activation, purified resting B cells were cultured at a density of 1.5-2× 10^6^/ml in B cell culture medium supplemented with 25mg anti-IgM antibody (Millipore) and 2ng/ml of IL-4 (PeproTech) for 20hrs. LPS (Sigma) was then added to a final concentration of 25μg/ml followed by incubation for 1 to 6hrs. For γ-secretase inhibition experiments, resting B cells were pre-cultured in B cell culture medium with 20 μM DAPT (Sigma) or with vehicle (DMSO) for 4hrs followed by addition of anti-IgM + IL4.

### FACS analysis

Cells (1× 10^6^) were collected by centrifugation, washed twice in PBS/2% FCS and the cell pellet was resuspended in 100μl of PBS/2% FCS. Fluorophore conjugated surface marker antibodies were added at a 1:100 dilution and incubated on ice for 20min. Cells were washed in PBS/2% FCS and resuspended in PBS/2% FCS for analysis on a BD LSR II Flow Cytometer. The antibodies used in the FACS analysis were as follows: CD21-FITC (553818) BD Biosciences Rat, CD23-PE (561773) BD Pharmingen Rat, CD23-Pacific blue (101616) BioLegend Rat, IgD-APC (405713) BioLegend Rat, IgM-PE-Cy7 (406513) BioLegend Rat, B220-Pacific blue (558108) BD Biosciences Rat, B220-PE (553089) BD Biosciences Rat. The LIVE/DEAD Fixable Aqua Dead Cell Stain Kit (Thermo Fisher Scientific) was used to distinguish between live and dead cells. Data was analysed with FlowJo Software. Gating strategy: Cells were initially gated for live lymphocytes followed by analysis of B220^+^ cells.

### FITC Annexin V/Dead Cell Apoptosis Kit

Cells (1×10^6^) were collected by centrifugation and washed in ice cold PBS. A FITC Annexin V/Dead Cell Apoptosis Kit (Invitrogen), containing FITC conjugated Annexin V and PI, was used to analyse the rate of apoptosis and death in cells. Cells were analysed on a BD LSR II Flow Cytometer.

### Cell cycle analysis by PI staining

Cells (1×10^6^) were collected by centrifugation and washed with PBS/2% FCS. The cell pellet was resuspended in 50μl of PBS/2% FCS. A volume of 500μl of ice cold 70% ethanol was added to the cell suspension, followed by 10 min incubation on ice. The sample was centrifuged and washed once in PBS/2% FCS. The pellet was resuspended in 50μl of PBS/2% FCS and incubated for 30min in the dark in 500μl of propidium iodide solution (PBS, 0.05mg/ml PI (Sigma), 0.05% NP40 (Sigma) and 1μg/ml RNase A (Sigma)). Samples were analysed on a BD LSR II Flow Cytometer (BD Biosciences).

### RNA analysis

B cells (3×10^6^) were pelleted by centrifugation and resuspended in 0.3 ml of Trizol (Thermo Fisher Scientific) and RNA spike-in (generated by in vitro transcription, sequence shown in Table S1) was added at a concentration of 0.1ng/1×10^6^ cells. Following RNA purification using the RNeasy mini kit (Qiagen) reverse transcription (RT) of 200 ng of RNA was carried out using the Super Script II reverse transcriptase kit (ThermoFisher). Real time quantitative PCR (RT-qPCR) analysis was carried out using the SensiMix SYBR No-Rox kit (Bioline), using primers and conditions shown in Supplemental Table 1. RNA levels were normalised using the following equation: (gene 2^−Ct^/spike 2^−Ct^) = gene expression relative to spike.

### RNA sequencing (RNA-seq)

B cells (30×10^6^) were lysed in 1ml of Trizol and 30μl of 1:100 diluted ERCC RNA Spike-In Mix (Ambion) was added to each lysate. RNA was isolated and eluted in a final volume of 20μl. 1μl of each sample was used for quality and concentration analysis on a 2100 Bioanalyzer using the RNA 6000 Nano kit (Agilent). 500ng of RNA was used to prepare each mRNA library. PolyA RNA selected RNA Libraries were generated by the LMS Core Genomics Facility using the TruSeq Stranded mRNA Library Prep Kit (Illumina). Paired-ended sequencing was performed on an Illumina HiSeq 2500 sequencer. The reads were aligned to mouse genome mm9 using Tophat2 with default parameters and gene annotation from Ensembl version66. Genome wide coverage of the RNA-seq datasets were generated as bedGraph files using BEDTools and converted to Bigwig files. Read counts on genes were computed using featureCounts function in Rsubread R package. Differentially expressed genes were identified using DESeq2 considering the covariates of interest and factors of unwanted variation computed using RUV-Seq.

### GSEA analysis

Mouse gene symbols were converted to human gene symbol using Ensembl Biomart (Hunt et al., 2018), Gene Set Enrichment Analysis (GSEA 2.2.0) was then performed with GseaPreranked tool using Hallmark and C2 Canonical Pathway gene sets.

### Quantitative Single Cell Imaging

Cells (60,000/well in 20 μl) were deposited in 384 well plates (CellCarrier 384-ultra – Perkin Elmer) coated with 0.01% polylysine and processed as follows: spin 800 g for 1 min; fixation in 2 % paraformaldheide/PBS for 15 min at RT; 3× 3 min wash with PBS; permeabilization with 0.3% tritonX/PBS for 15 min; 3× 3 min wash with PBS; blocked overnight with 0.2% fish gelatin (SIGMA)/5% horse serum/PBS (block solution); 1st antibody staining in block for 1 hr at RT. 2× 3 min wash with PBS/0.2% fish gelatin and 1× 3 min wash with PBS; 2nd antibody staining in block for 1 hr at RT. 1× 3 min wash with PBS/0.2% fish gelatin and 1× 3 min wash with PBS; Hoescht staining for 15 min at RT (in PBS, 5 ug/ml final); 2× 3 min wash with PBS. Cells were kept in PBS/0.05 Na azide, sealed with foil at 4C. Fixed and immunostained cells were imaged on the Operetta HCS CLS (PerkinElmer) with a 40x water immersion objective, NA 1.1. Quantitative, automated image analysis was performed using Harmony software (PerkinElmer). Nuclei were detected and segmented based on Hoechst intensity and nuclei touching the edge of the field were filtered out. Clumps of nuclei and dead nuclei were excluded based on nuclear area and Hoechst intensity such that only single nuclei were included in the analyses. The intensity of nuclear proteins was quantified using the nuclear segmentation mask. Due to B-cells having a very small cytoplasm, intensities of cytoplasmic proteins were quantified by segmenting a ring around the edge of the nuclear segmentation mask, and three pixels wide (referred to as the ‘ring region’). Single-cell and average well data were plotted using Prism 8 software (GraphPad).

### ChIP and ChIPseq

Chromatin immunoprecipitation (ChIP) analysis of resting B cell were performed on formaldehyde-fixed cells as described in (18), with the following antibodies: RUNX1 (Abcam- ab23980), SRCAP (Novus Biologicals- NBP145244), H3K27me3 (NEB- 9733S), RING1B (MBL - D139-3), RUNX3 (Abcam- ab135248), BRG1 (Abcam- ab110641), H2A.Z (Abcam- ab150402), Normal Rabbit IgG (Cell Signaling Technologies -2729S) and H3K4me3 (Merck Millipore- 07-473). qPCR-ChIP was carried out using the SensiMix SYBR No-Rox kit (Bioline), with the primers and conditions shown in Supplemental Table 1. ChIPseq library preparation was carried out on 3 independent ChIP replicates. ChIPseq replicates were pooled for further analysis. Peak calling was performed on the ChIPseq samples using corresponding input samples and MACS v2.1.2. ChIPseq read profiles were normalized to Reads Per Million (RPM) normalized values with bedtools and UCSC RPM tracks were generated for visualization.

### RNA pol II, H3K4me1 and H3K27ac ChIPseq datasets

RNA Pol II binding data for resting B cells was downloaded from the Short Read Archive (accession numbers SRR955859, SRR955860, SRR955861) (19). The IgG control data for resting B cells was downloaded from the Gene Expression Omnibus under accession number GSE24178 (20). Replicates were merged for downstream analysis. The Chipseq data with read length 50 bps were aligned to the mouse reference genome mm9 using Bowtie (version 1.1.1). Normalized custom tracks in bigwig format were then prepared using deepTools (version 3.2.1) and visualized in UCSC genome browser. H3K4me1 and H3K27ac ChIPseq datasets for mouse splenocytes were downloaded from the ENCODE Consortium dataset (www.encode.org).

### Proximity Ligation Assay (PLA)

PLAs were performed with the Duolink in situ PLA kit (Sigma, goat and rabbit probes) following the manufacturer’s instructions. Images were collected by confocal microscopy using an SP5 microscope (Leica Microsystems, Wetzlar, Germany) and LAS-AF software. The following antibodies were used: RUNX1 (ab23980), RUNX1 (ab35962), SRCAP (NBP145244) and SRCAP (ab9948).

### Immunofluorescence

Immunofluorescence was carried out as described in (20).

### Western blotting

Cells were sonicated (MSE Soniprep150, 5 times 30 sec on/30 sec off, 14 μm amplitude), subjected to SDS PAGE, transferred to nitrocellulose membranes and blocked with 5% dry milk (BSA) for 1hr at room temperature. Membranes were incubated o/n with the primary antibody diluted in 0.5% milk at 4C. They were then washed 3× 10 min, incubated with the appropriate secondary antibody for 1h at room temperature, washed 3× 10 min and developed using Millipore Crescendo ECL (Merck).

## RESULTS

### *Runx1*-knockout resting B cells are hyperresponsive to activating stimuli

The *Runx1* c-k/o mice generated by crossing a CD23-cre transgene (17) with a *Runx1* allele that has intronic LoxP-sites flanking exon 4 of *Runx1* (16) (see Methods), lacked RUNX1 protein at the mature resting B cell stage (Figure S1A). The mice had spleens and lymph nodes of apparently normal size, but FACS analysis measuring surface expression of the pan-B cell marker B220 showed reductions in the number of B cells in spleen (Figure 1A, top left panel) and in lymph nodes (Figure 1C, top left panel). A similar reduction was observed in the number of resting B cells that could be isolated from the Runx1 c-k/o spleens (Figure 1A, top middle panel).

**Figure 1.**
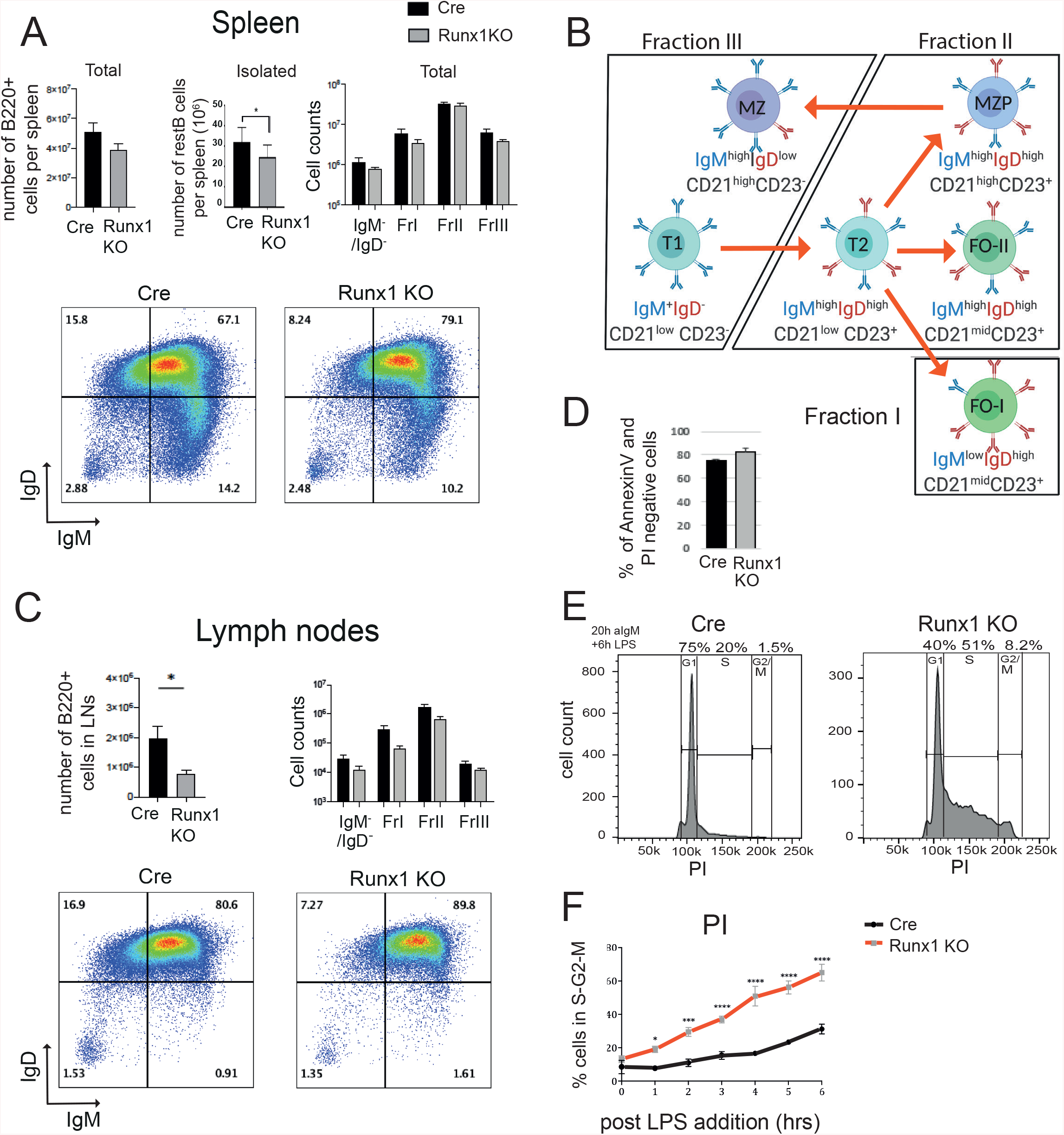
Runx1 knockout affects responsiveness of B cells to BCR stimulation. **A.** Top panel shows number of B220^+^ cells per spleen in CD-23-cre and Runx1 c-k/o (Runx1 KO) (left), number of resting B cells isolated from spleens (middle), and the numbers of fraction I-III B cells and IgD^−^IgM^−^ cells (right) in the spleens of Cd23-Cre and Runx1 c-k/o mice. For A, C and D, values are mean ± SEM *P<0.05, Student’s t-test, n=3. Bottom panel: FACS plot for splenic B220^+^ B cells showing staining for surface IgM and IgD expression. Numbers show fractions of cells based on gating for levels of IgM and IgD. Plots are representative of three Runx1 c-k/o and three control CD23-Cre spleen analyses. Percentages of total live cells for each gate are shown. **B**. Schematic representation of the cells in Fractions I, II and III (21, 22). T1 = transitional type 1, T2 = transitional type 2, MZP = marginal zone precursor, MZ = marginal zone, FO-I = type I follicular, FO-II = type II follicular. **C**. Top left panel: The percentage of B220+ B cells in cervical lymph nodes. Top right panel: Mean number of Fraction I-III B220^+^ B cells in lymph nodes. Bottom panel: FACS plot of lymph node B220^+^ B cells gated according to levels of IgM and IgD surface expression. **D**. Analysis of survival of ex vivo cultured Runx1 c-k/o resting B cells by staining unfixed cells with anti-Annexin V and propidium iodine (PI). The y-axis shows the percentage of cells that were negative for Annexin V and PI. **E**. Representative FACS analysis of PI-stained CD23-cre and Runx1 c-k/o B cells after 20 hrs treatment with anti-IgM followed by treatment with LPS for 6hrs. **F**. The proportion of cells in S or G2/M phase of the cell cycle after 18 hrs stimulation with anti-IgM followed with 0-6hrs of LPS treatment. Cells were fixed and DNA content was measured by PI staining and FACS analysis. Mean ± SEM. * p≤0.5, ** p≤0.01, *** p≤0.001, **** p≤0.0001, Student’s t-test. n= 3.

To assess the relative proportions of B cell sub-populations in the *Runx1* c-k/o mice, total splenocyte and cervical lymph node cell populations were stained for B220, IgM, IgD, CD23 and CD21 and staining profiles were analysed by FACS. B cells were identified by gating for B220 staining and the relative proportions of IgD^high^/IgM^low^ (Fraction I), IgD^high^/IgM^high^ (Fraction II) and IgD^low^/IgM^high^ (Fraction III) (21, 22) (Figure 1B) were measured (Figure 1A and 1C, top right and bottom left and right panels). Transitional (T: CD23 ^low^ /CD21^low^), follicular (FO: CD23^high^/CD21^mid-high^) and marginal zone (MZ: CD23^low^/CD21^high^) B cells were also identified among the B220 positive cells from spleen on the basis of CD23 and CD21 staining (Figure S1C, left panel). The analysis showed that there was relatively little change in the proportions of Fractions I, II and III in the spleens of *Runx1* c-k/o mice (Figure 1A and 1B). However, a small reduction in the proportion of FO and an increase in the number of marginal zone (MZ) cells was observed in the knockout spleens (Figure S1C, left panel). Increased surface IgD was also observed on MZ cells (Figure S1C, middle and right panels), which could be related to the known role of RUNX1 in promoting IgA class-switching (23). We conclude from these results that B cell numbers were reduced in the Runx1 ck/o mice, but the relative proportions of the B cell subsets were not substantially altered.

CD21 is a component of the BCR co-stimulator complex and a significant increase in CD21 surface levels was identified in the Runx1 c-k/o mice on B cells (B220^+^) in spleen and lymph nodes (Figure S1D and S1E). In the spleen, this increase was observed for Fractions I and II, but not for Fraction III (Figure S1D, bottom panels). It is likely that the increase in the number of CD21^high^ B cells in both Fractions I and II is caused by an overall increase in the expression of CD21 in follicular B (FO) cells, rather than being due to the relatively small increase that was observed in the proportion of MZ cells. This conclusion is supported by the observed increase in CD21 surface levels on the B cells in lymph nodes, which contain follicular cells (FO) but lack MZ and MZP cells (Figure S1E).

The lower overall number of B cells in the *Runx1* c-k/o spleens could be due to reduced survival of the naïve mature B cells. However, splenic resting B cells isolated from the *Runx1* c-k/o mice and cultured for 24hr in the presence of IL-4 showed an increase in the proportion of cells that lack apoptotic markers compared with CD23-Cre (Cre) control cells (Figure 1D). Another possible explanation for the reduced number of B cells could be that the mature splenic B cell pool is depleted by increased responsiveness to antigenic stimulation (24). We therefore set out to investigate whether the *Runx1* c-k/o B cells have an altered response to activating stimuli.

Incubation of resting B cells with anti-IgM has been shown to result in entry of the cells into G1 within 24 hours of the start of the incubation, with entry into S-phase occurring by 48 hours (25, 26). Treatment of resting B cells with lipopolysaccharide (LPS) alone has been reported to result in a limited activation response by the follicular mature (FM) B cells that make up 95% of the resting B cell population, due to FM resting B cells having a limited capacity to respond to toll-like receptor (TLR) stimulation (25). However, cells that have been pre-stimulated with anti-IgM respond rapidly to LPS stimulation (27). This allowed us to induce a controlled entry of purified wild-type and *Runx1* c-k/o resting B cells into the cell cycle by incubating the cells for 20 hours with anti-IgM followed by a time-course of lipopolysaccharide (LPS) treatment for 1 - 6 hours. Staining of the cells with propidium iodide (PI) (Figure 1E) showed a mean increase of approximately 3-fold, from 15% to 45%, in the proportion of *Runx1* c-k/o B cells that had progressed into S-phase compared with wild-type cells after 4 hours of LPS stimulation and a 2-fold increase from 30% to 60% after 6 hours of stimulation (Figure 1F). These results indicate that the *Runx1* c-k/o resting B cells were in a hyperresponsive state after stimulation of the BCR with anti-IgM that caused them to respond rapidly to TLR stimulation with LPS. In contrast, wild-type resting B cells showed very little response to LPS stimulation under the same conditions.

### RUNX1 antagonises BCR-mediated upregulation of genes that affect cell cycle progression

To investigate the reasons for the enhanced responsiveness of the *Runx1* c-k/o resting B cells to BCR stimulation, RNA-seq was used to analyse global gene expression patterns in wild-type and *Runx1* c-k/o B cells at the resting B cells stage and after 3 hours incubation with anti-IgM. The analysis showed that a total of 167 genes were upregulated and 187 genes were downregulated in the Runx1 c-k/o resting B cells and 409 genes were upregulated and 277 genes downregulated in the Runx1 c-k/o B cells after 3 hrs of stimulation with anti-IgM (adjusted p value < 0.05) (Figure 2A and S2A, Table S2A and S2B). Gene Set Enrichment Analysis (GSEA) revealed that there was upregulation of genes associated with G1-S transition, S-phase, BCR signalling and protein translation functions at 3 hrs post-anti-IgM addition in the *Runx1* c-k/o B cells compared with the level observed in wild-type cells (Figure 2B and S2B). The GSEA analysis also revealed upregulation of signalling and immune response genes at both 0 and 3 hrs after anti-IgM addition and downregulation of late cell cycle G2-M and mitotic spindle genes (Figure S2B).

**Figure 2.**
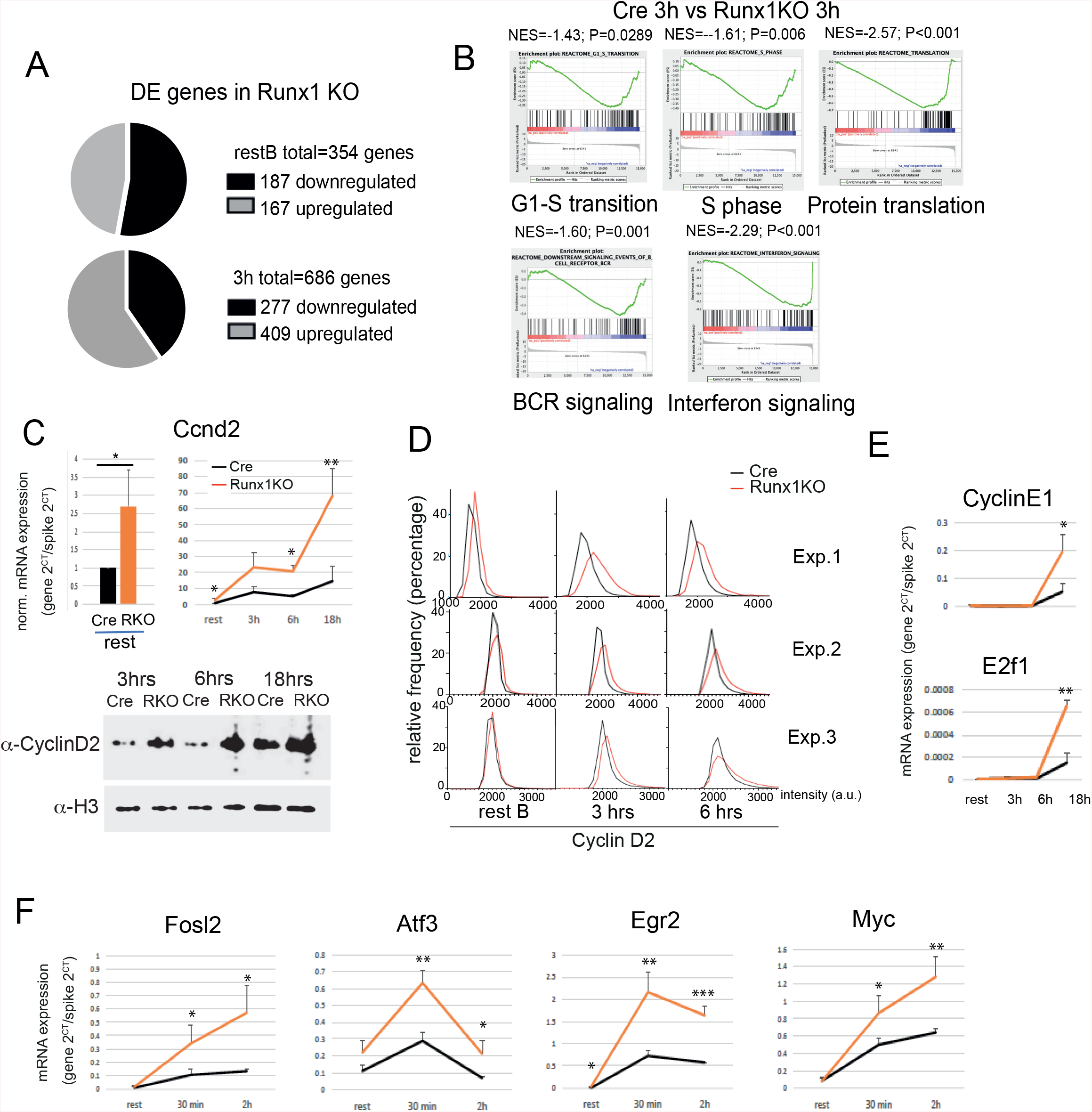
Expression of cell cycle genes during BCR-mediated activation of Runx1 knockout B cells. **A**. RNA-seq analysis of Runx1 c-k/o B cells showing upregulation and downregulation of genes in resting B cells (upper pie chart) and in 3hrs anti-IgM activated B cells (lower pie chart) with respect to CD23-Cre control B cells. Three biological replicates were analysed for each cell type. **B.** GSEA of the RNA-seq data from the 3hrs anti-IgM activation samples revealed that cell cycle and cell signalling gene categories are upregulated in the Runx1 c-k/o. Values below 0 in the graphs refer to genes whose expression is higher in the 3hrs activated Runx1 c-k/o B cells with respect to 3hr activated CD23-Cre B cells. NES = Normalised Enrichment Score. P= nominal p-value. **C**. RT-qPCR analysis of expression of the *Ccnd2* gene in resting B cells (top left panel) and following activation for the indicated times with anti-IgM (top right panel). Bottom panel: western blot analysis of cyclin D2 protein levels following anti-IgM activation for the indicated times. Loading control: histone H3. **D**. Single cell analysis of the level of cyclin D2 protein in resting B cells and after 3 and 6 hrs anti-IgM treatment using quantitative, automated single cell image analysis (see Methods). Three independent experiments are shown. **E-F**. RT-qPCR analysis of cell cycle gene transcription (**E**) and immediate early gene transcription (**F**) in the Runx1 c-k/o B cells upon activation with anti-IgM for 0-18 hrs. For C, E and F, values are mean ± SD. * p≤0.5, ** p≤0.01, *** p≤0.001, Student’s t-test, n=3

In order to gain additional insights into the hyperresponsive phenotype of the *Runx1* c-k/o B cells, we carried out a more detailed analysis of changes in the expression of genes that are involved in regulating entry into the cell cycle. Cyclin D2 is the main cyclin D involved in B cell activation as indicated by the failure of cyclin D2-deficient B cells to proliferate in response to BCR signalling (28). We analysed the kinetics of cyclin D2 (*Ccnd2*) expression in isolated *Runx1* c-k/o resting B cells upon activation with anti-IgM during a time course from 0 to 18 hrs. RT-qPCR analysis of *Ccnd2* mRNA showed a large increase in *Ccnd2* mRNA levels in the knockout after 3 hours treatment with anti-IgM and continuing through to 18 hours (Figure 2C, top panel).

The increase in cyclin D2 expression was confirmed by western blotting, which showed substantial increases in the level of cyclin D2 protein in *Runx1* c-k/o cells relative to wild-type cells at 3, 6 and 18 hrs after addition of anti-IgM (Figure 2C, bottom panel). The analysis of cyclin D2 protein was further extended by quantitative single cell protein imaging of cyclin D2 levels in resting B cells and after 3 and 6 hrs anti-IgM treatment (see Methods for details). The results of this analysis, obtained from three independent experiments, showed that the increase in cyclin D2 protein level over time is unimodal, with the effect distributed across the entire cell population rather than being confined to a subset of cells (Figure 2D). These data indicate that the presence of RUNX1 in resting B cells reduces *Ccnd2* transcription and is associated with a lower rate of increase in the level of *Ccnd2* mRNA and protein as the cell progresses through G1.

In contrast to the effect of the *Runx1* knockout on cyclin D2 expression, levels of transcription of the genes encoding cyclin E1 (*Ccne1*) and the transcription factor E2F1 (*E2f1*) were unchanged in Runx1 c-k/o resting B cells and after 3- and 6-hours incubation with anti-IgM compared with wild-type B cells (Figure 2E). Increased levels of *Ccne1* and *E2f1* mRNA were only observed after 18 hours of anti-IgM stimulation. This is consistent with previous reports that *Ccne1* transcription is upregulated in late G1 by the E2F factors (29). *E2f1* transcription has been shown to be subject to auto-activation by E2F factors that have been activated by Rb phosphorylation (30).

We also examined the activation profile of several immediate early genes in response to anti-IgM stimulation. Levels of transcription of the *Fosl2*, *Atf3, Egr2* and *Myc* genes were analysed at timepoints from 0 to 2 hr after addition of anti-IgM. Expression of *Atf3* and *Egr2* peaked at 30 minutes in wild-type cells and then declined. This profile was maintained in the *Runx1* c-k/o B cells, but the expression peak at 30 minutes was strongly enhanced and overall expression was increased at all three time-points (Figure 2F). Expression of *Fosl2* and *Myc* showed a progressive increase at 30 minutes and 2 hrs in wild-type and the slope of the expression curve was shifted upwards for both genes in the *Runx1* c-k/o cells (Figure 2F). These results implicate *Runx1* in the downregulation of cell cycle and immediate early genes during BCR-mediated activation of naïve resting B cells.

### RUNX1 binds to poised cell cycle and immediate early genes in resting B cells

ChIP-sequencing (ChIPseq) was carried out on chromatin from wild-type resting splenic B cells in order to examine profiles of factor binding and epigenetic modification at genes that showed altered expression in the *Runx1* c-k/o B cells (see Methods for details). RUNX1 binding peaks were found to be widely distributed at promoters and putative enhancers of the affected genes (Figure 3A – 3D and Figure S3). The *Ccnd2* and *Atf3* genes had strong peaks of RUNX1 binding at distally located elements that had the epigenetic characteristics of enhancer or silencer sequences (Figure 3A and 3C). Strong binding peaks for RUNX1 were located close to the TSS of the *Egr2* and *Fosl2* genes (Figure 3B and 3D) in regions with epigenetic profiles that are indicative of overlapping promoter/enhancer sequences (31). Overall, these results suggest that RUNX1 binds to promoters and enhancers at cell cycle and immediate early genes in resting B cells.

**Figure 3.**
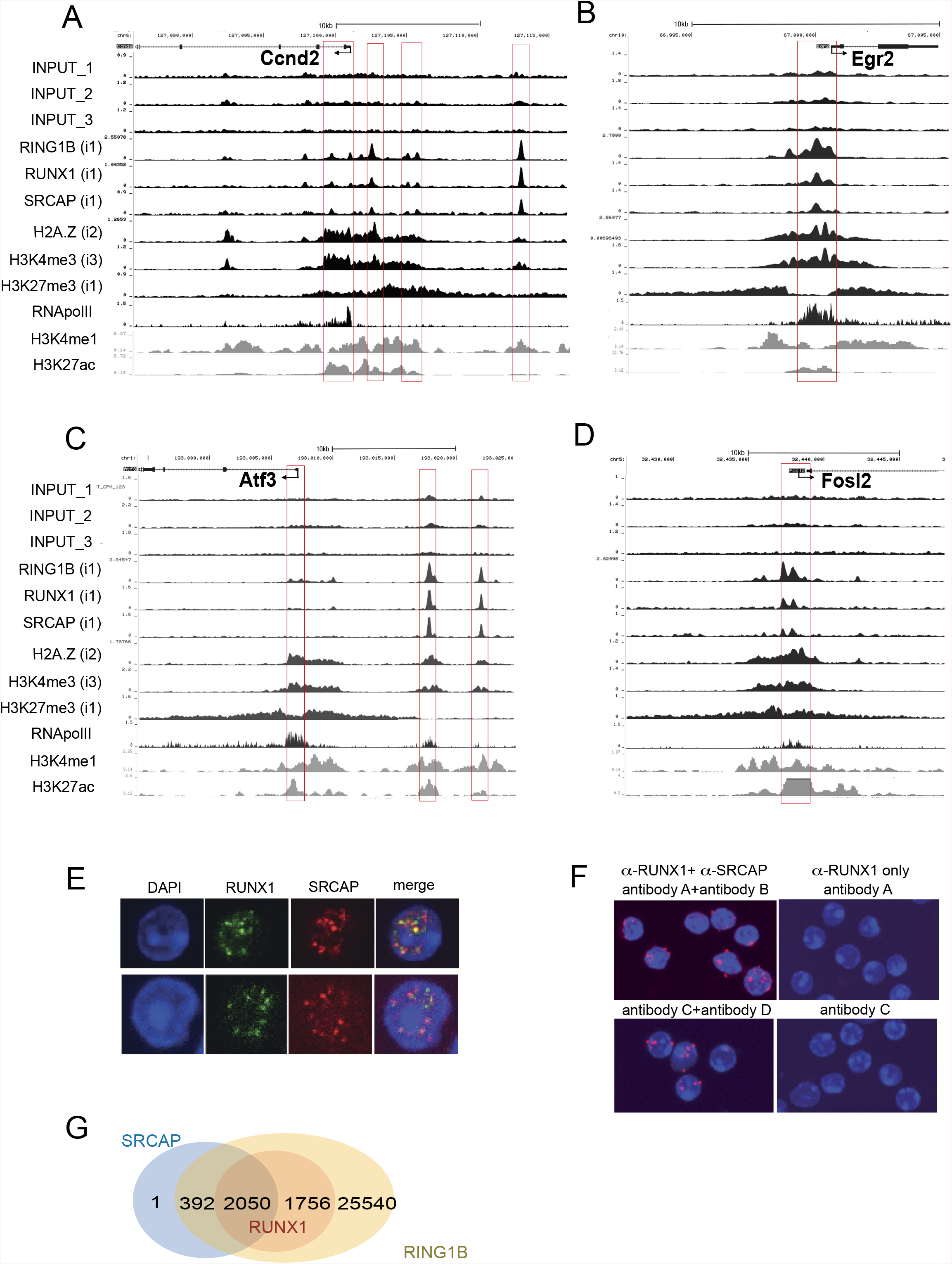
ChIP-seq analysis of cell cycle and immediate early genes. Tracks shown in black represent ChIP-seq profiles from purified resting B cells for *Ccnd2* (**A**) *Egr2* (**B**) *Atf3* (**C**) and *Fosl2* (**D**). Grey tracks represent data from splenocytes downloaded from ENCODE. The data for RNA pol II was downloaded from a published dataset obtained by analysis of purified resting B cells (19) (see Methods). The inputs for each track are indicated by numbering of the input tracks with the relevant input number shown in brackets for each analytical track. Red boxes highlight the positions of Transcription Start Sites (TSS) and the position of candidate enhancers co-occupied by RUNX1, SRCAP and RING1B. **E.** Localisation of RUNX1 and SRCAP was determined by immunofluorescence analysis of resting B cells. **F.** The proximity ligation assay (PLA) was carried out on resting B cells using 2 sets of antibodies for RUNX1 (antibody A and C) and for SRCAP (antibody B and D). Red = PLA amplification signals. Blue = Hoechst dye staining the nuclei. **G**. Venn diagram summarizing the degree of overlap of peaks by ChIP-seq peak calling for binding of RUNX1, SRCAP and RING1B in resting B cells.

RUNX1 has the characteristics of a pioneer factor which can initiate binding to closed nucleosomal chromatin (32) and has also been shown to interact with the Swi/Snf-like Brg1/Brm-associated factor (BAF) chromatin remodelling complex in thymocytes (33). Our initial ChIP analysis of chromatin remodelling subunits indicated that components of another remodelling complex, the SNF-2-related CREB-binding protein activator protein (SRCAP) complex, bind to active genes in resting B cells and to genes that are inducible by activation, with the level of binding to both categories of genes reduced when the B cells were activated (data not shown). ChIPseq analysis of the SRCAP protein revealed that binding peaks for SRCAP co-localised with RUNX1 at poised and active genes (Figure 3A – 3D). The SRCAP nucleosome remodelling complex is responsible for incorporating histone H2A.Z into nucleosomes (34), and our analysis showed that H2A.Z is enriched at active and the poised promoters in resting B cells (Figure 3A – 3D).

The binding of RUNX1 and SRCAP to the same regions at hyperresponsive genes in resting B cells, suggests that RUNX1 might be interacting with SRCAP and recruiting it to poised genes. The very large size of the SRCAP protein (360 kD) makes it challenging to carry out co-immunoprecipitation experiments. Therefore, we used immunostaining of SRCAP and RUNX1 to examine localisation of the two proteins. The results show that strong staining foci for both proteins with overlap observed between the majority of the foci (Figure 3E). Proximity ligation assays (PLA) performed with two independent sets of antibodies against RUNX1 and SRCAP confirmed that there is close contact between RUNX1 and SRCAP in the resting B cells (Figure 3F). This is consistent with the genome-wide co-localisation of SRCAP binding with binding peaks for RUNX1 (Figure 3G)

The ChIPseq analysis also revealed that a number of the hyperresponsive cell cycle and immediate early genes identified in the expression analysis had a poised epigenetic profile in wild-type resting B cells. The promoter-proximal regions of these genes had broad regions of enrichment for histone H3K27me3 (Figure 3A–3D), which is deposited by the polycomb PRC2 complex member EZH2 and is considered to be a repressive mark. Enrichment for H3K27me3 is also associated with poised promoters in ES cells where it forms part of the bivalent H3K4me3/H3K27me3 epigenetic mark, which has been associated with promoter poising in ES cells (35, 36). The regions of H3K27me3 enrichment that we observed at the hyperresponsive cell cycle and immediate early genes in resting B cells showed partial overlap with regions of enrichment for H3K4me3 (Figure 3A–3D).

RING1B, which is the catalytically active component of the polycomb PRC1 complex (37), has been shown to bind to poised and active genes in a variety of cell types (38–42), and RUNX1 and RING1B have been reported to co-occupy binding regions in megakaryocytes and T cells (42). Our ChIPseq analysis showed strong co-localisation of binding of RUNX1 and RING1B in resting B cells (Figure 3A – 3D and Figure 3G). The overlapping RUNX1 and RING1B peaks were associated with distally located sequences that had the epigenetic signature of enhancers at the *Ccnd2* and *Atf3* genes and were also found at or close to promoters of genes that were hyperresponsive in *Runx1* c-k/o B cells (Figure 3A – 3D). The co-localised peaks of RUNX1 and RING1B binding in resting B cells did not show any specific co-localisation with the regions of enrichment for H3K27me3, similar to our previous observation that RING1B binds at active genes in resting B cells, independently of H3K27me3 (39).

To examine RNA Pol II binding at poised promoters, we downloaded published ChIP-seq data for RNA-Pol II in resting mouse B cells (19). The ChIPseq tracks show high levels of RNA Pol II in the regions around the promoters of the *Ccnd2*, *Egr2*, *Atf3*, and *Fosl2*, genes compared with the levels of Pol II in the gene bodies (Figures 3A – 3D and Figure S3E). The presence of higher levels of Pol II at promoter regions compared with the levels observed in the body of the gene have been shown to be a characteristic feature of poised genes in *Drospophila* embryonic cells (43) and mouse ES cells (44).

### Knockout of *Runx1* reduces binding of SRCAP and increases binding of BRG1 and RING1B at the *Ccnd2* gene

To further analyse the role of RUNX1 in configuring the epigenetic profile of poised cell cycle genes in resting B cells, we focused on the promoter and upstream region of the *Ccnd2* gene. Cyclin D2 has critical roles in B cell activation and has also been shown to be positively regulated by the *RUNX1*-ETO fusion oncoprotein in acute myeloid leukemia (AML) (45). We first carried out an analysis of enrichment of H3K27me3 and H2A.Z and binding of SRCAP, BRG1, RING1B and the RUNX1 paralog RUNX3 at the *Ccnd2* promoter in wild-type resting B cells and after 20 hrs treatment with anti-IgM. The results of the analysis, which are shown in Figure 4A, revealed that BCR stimulation of wild-type cells resulted in a statistically significant reduction in binding of RUNX1 and SRCAP and significantly increased binding of BRG1 and RUNX3. Binding of the PRC1 component, RING1B, was significantly increased and enrichment for the PRC2 marker, H3K27me3, was strongly reduced in the activated cells whereas H2A.Z was largely unchanged. The fact that RUNX3 showed a strong increase in binding during BCR activation, which coincided with decreased binding of RUNX1 (Figure 4A), suggests that there is a switch between the two Runt-related factors. This is supported by the observation that the level of RUNX1 protein declines upon activation of the resting B cells (Figure S1A), while RUNX3 protein is upregulated (Figure S1B; see also (46)). The large increase in binding of BRG1 (Figure 4A), suggests that there is also a switch between binding of the SRCAP and SWI/SNF remodelling complexes to the *Ccnd2* promoter during B cell activation.

**Figure 4.**
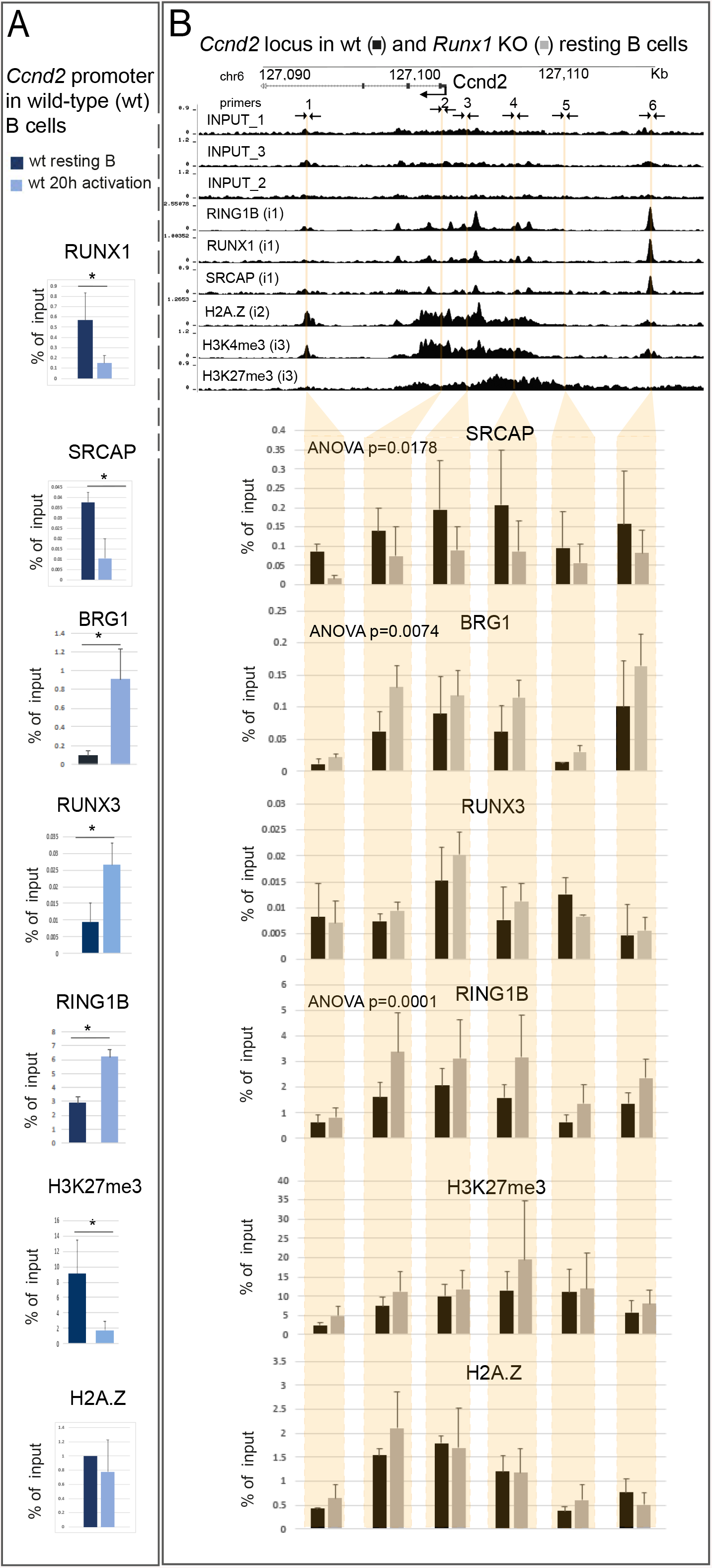
Analysis of the epigenetic architecture of the *Ccnd2* promoter in B cells. **A**. ChIP-qPCR of the indicated proteins and histone modifications at the *Ccnd2* promoter in wild-type (wt) resting B cells (dark blue) and after 20hrs anti-IgM activation (light blue). See Table S1 for primer pair sequence. Values are mean ± SD, n = 3. * p≤0.5, ** p≤0.01, Student’s t-test. **B**. ChIP-qPCR at 6 DNA sites corresponding to the *Ccnd2* promoter and surrounding regions in CD23-Cre (black) and Runx1 c-k/o resting B cells (grey). Vertical lines show positions of primer pairs used for the ChIP analysis (Table S1). (n=3). Mean ± SD. P values were calculated by ANOVA.

To analyse the effect of the *Runx1* knockout on factor binding and epigenetic marks across the regions upstream and downstream from the *Ccnd2* TSS, a series of qPCR-ChIP primers were used that covered the *Ccnd2* promoter region as well as candidate upstream and downstream regulatory elements and the domain of H3K27me3 enrichment (Figure 4B). The results of the qPCR-ChIP carried out on wild-type and *Runx1* c-k/o resting B cells showed that the *Runx1* knockout led to a clear and significant decrease in binding of SRCAP and a significant increase in binding of BRG1 at the promoter and upstream and downstream regions (Figure 4B). Binding of RING1B, was also significantly increased. Binding of RUNX3 and enrichment for H3K27me3 and H2A.Z were largely unaffected by the *Runx1* knockout (Figure 4B).

Taken together, the results of the ChIPseq and ChIP-qPCR analysis reveal binding of RUNX1, SRCAP and RING1B accompanied by a poised bivalent epigenetic configuration and enrichment for H2A.Z at several cell cycle and immediate early genes in resting B cells. Knockout of RUNX1 reduces SRCAP binding and increases binding of the activation-associated SWI/SNF component BRG1 and binding of RING1B at the *Ccnd2* gene, suggesting that RUNX1 recruits SRCAP to enhancers and promoters, but has a more complex relationship with RING1B binding. The ChIP analysis also shows that RUNX1 is largely replaced by RUNX3 at the *Ccnd2* promoter during BCR-stimulated activation of wild-type resting B cells. RUNX3 has been reported to interact with BRG1, recruiting it to promoters (47), suggesting that it could be involved in mediating the switch between SRCAP and SWI/SNF binding to the *Ccnd2* promoter.

### RUNX1 regulates expression levels of immunomodulatory and autoimmune disease-associated genes in resting and activated B cells

The GSEA analysis indicating altered expression of genes that affect BCR signalling (Figure 2B), led us to consider whether RUNX1 has a broader role in regulating expression of genes that affect BCR responsiveness, B cell tolerance and autoimmunity. To test whether this is the case, we scanned the genes that were identified by the RNA-seq analysis as being significantly up or downregulated in the *Runx1* c-k/o B cells, for published associations with modulation of BCR functioning and/or autoimmune disease. A total of 14 genes were identified that fitted into one of these categories and showed significant expression changes (Table 1, Figure S3B, S3D). Comparison of the expression profiles of the genes in wild-type resting B cells and after 3 hours activation with anti-IgM revealed that the majority showed increased expression in the RUNX1 c-k/o B cells (Table 1, Figure S3B, S3D). This suggests that the main effect of RUNX1 on the immunomodulatory genes is to control the level of expression in resting and activated B cells rather than affecting the direction of the expression change.

**Table 1.** Changes in expression of autoimmunity-related genes in *Runx1* c-k/o measured by RNA-seq analysis of resting B cells and after 3 hours activation with anti-IgM. Adjusted P values for all differences shown were <0.05 (see Tables S2A and S2B for the full set of base mean values). Red arrows indicate fold-downregulation in the *Runx1* c-k/o B cells. Green arrows indicate fold-upregulation.

| Gene | Base Mean | Fold Change_0hrs | Fold Change_3hrs | Functions | Autoimmune disease | References |
| --- | --- | --- | --- | --- | --- | --- |
| *Ptpn22* | 995 | 1.3 ↓ | 1.9 ↓ | Phosphatase – BCR regulation | Lupus | (59) |
| *Lrrk2* | 6056 | 1.7 ↓ | 3.7 ↓ | Autophagy; lysosome functioning | Lupus; Parkinson's disease; Colitis | (60, 61) |
| *Ccr6* | 974 | 3.0 ↓ | 4.0 ↓ | Chemokine receptor | Inflammatory bowel disease | (62) |
| *Cd22* | 11926 | 1.2 ↓ | 1.4 ↓ | Inhibition of BCR signaling | Lupus | (63, 64) |
| *Cd47* | 2638 | 1.2 ↑ | 2.3 ↑ | Regulation of B cell maturation and homeostasis | Lupus | (65) |
| *Irf7* | 1046 | 2.0 ↑ | 3.5 ↑ | Regulation of interferon response | Lupus | (66) |
| *Irf5* | 3126 | 1.1 ↑ | 1.4 ↑ | Regulation of interferon response | Lupus | (67) |
| *Ifnar1* | 4324 | 2.5 ↑ | 1.9 ↑ | Subunit of interferon-α receptor | Lupus | (68, 69) |
| *Zbp1* | 787 | 1.1 ↑ | 2.1 ↑ | DNA-dependent activator of interferon-regulatory factors (DAI) | Lupus | (70, 71) |
| *Trib1* | 1081 | 1.1 ↑ | 1.9 ↑ | Regulation of MAPK pathways | Lupus | (72) |
| *Lgals9* | 1512 | 1.2 ↑ | 1.7 ↑ | Regulation of cell survival and cytokine production | Lupus | (73) |
| *Stat1* | 3506 | 1.0 ↑ | 1.5 ↑ | Enhancement of interferon-induced gene expression | Lupus | (74) |
| *Tnfrsf13c* | 4888 | 1.2 ↑ | 1.4 ↑ | Encodes BAFF receptor | Lupus | (75, 76) |
| *Bank1* | 7290 | 1.0 ↑ | 1.3 ↑ | Enhancement of BCR signalling | Lupus | (77) |

Examination of ChIPseq profiles for the immunomodulatory genes showed strong co-localisation of binding peaks for RUNX1, SRCAP and RING1B and in the regions around their promoters and putative enhancer elements (Figure S3A, S3C). Interestingly, the immunomodulatory genes that were upregulated in the *Runx1* c-k/o B cells at 0 and 3hrs anti-IgM stimulation did not, in general, show enrichment for H3K27me3 close to the genes or high levels of poised RNA Pol II at their promoters (Figure S3A). The exceptions to this were the *Irf7* gene, which has a region of H3K27me3 upstream from the promoter and can be considered to be bivalent, and the *Ifnar1* gene, which lacks H3K27me3 in the region around the gene. Both genes have high levels of RNA Pol II at their promoters, relative to the levels across the gene bodies, which indicates promoter poising (Figure S3A). These results are indicative of the heterogeneity of gene regulation mechanisms that have evolved separately to generate similar outcomes.

The functions of this group of genes and their links with autoimmune disease are summarised in Table 1. A number of the genes that are repressed by RUNX1 are associated with the interferon response. The interferon alpha and beta receptor subunit 1 (*Ifnar1*) and the transcription factor interferon regulatory factor 7 (*Irf7*) genes showed particularly strong upregulation in the *Runx1* c-k/o resting B cells and after 3hrs of anti-IgM treatment. Of the genes that are positively regulated by RUNX1, *Ptpn22* and *Cd22* have been associated with lupus in humans and mice, mutations in *Lrrk2* have a causal role in familial Parkinson’s disease and *Ccr6* is associated with inflammatory bowel disease. These results suggest that RUNX1 is an important regulator of genes that are associated with B cell tolerance and autoimmunity.

### Notch signalling is enhanced in hyperresponsive *Runx1* c-k/o resting B cells

Among the genes that were upregulated in the hyperresponsive *Runx1* c-k/o resting B cells were components of the Notch signalling pathway. Signalling through the transmembrane NOTCH receptors occurs when the receptors are bound by ligands belonging to the Jagged and Delta-like (DLL) families. This leads to cleavage and release of the extracellular domain of the receptors, followed by a second cleavage mediated by presenilin (PSEN) family members, which act as the catalytic subunits of the γ-secretase complex. PSEN-mediated cleavage results in release of the Notch intracellular domain (NICD), which then translocates to the nucleus where it binds to RBPJ, forming a complex that recruits other co-activators and upregulates promoter of target genes (reviewed in (48, 49)). Notch signalling via NOTCH2 has generally been associated with specification of marginal zone B cells during B cell maturation, whereas development of follicular B cells was left unaffected by conditional knockout of *Notch2* (50, 51). However, stimulation of the Notch pathway has been shown to increase proliferation of purified splenic follicular B cells in response to anti-IgM stimulation (52), suggesting that Notch signalling does have a role in regulating the dynamics of B cell activation.

Our analysis of Notch pathway genes revealed that expression of *Rbpj*, *Notch2*, and *Psen2* was altered in the *Runx1* c-k/o B cells. All three genes showed elevated expression at the resting B cell stage. *Notch2* and *Rbpj* also showed strongly elevated expression after 18 hours of stimulation (Figure 5A, C) whereas expression levels of *Psen2* in the knockout cells was increased in resting B cells and after 30 min of anti-IgM activation (Figure 5E), before declining. Epigenetic and factor binding analysis of the *Rbpj* promoter (Figure 5B) revealed a similar profile to the *Ccnd2* gene (Figure 4B), suggesting an overlap in regulatory mechanisms for the two genes. We also used single cell analysis to measure the levels of NOTCH2 protein in the nuclei of wild-type and *Runx1* c-k/o resting B cells. The results of three independent experiments showed increased levels of nuclear NOTCH2 in *Runx1* c-k/o resting B cells (Figure 5D). The observation that distribution of the increased levels of Notch2 in the nuclei of isolated resting B cell populations is largely unimodal indicates that the *Runx1* knockout increases the level of nuclear NOTCH2 in follicular B cells.

**Figure 5.**
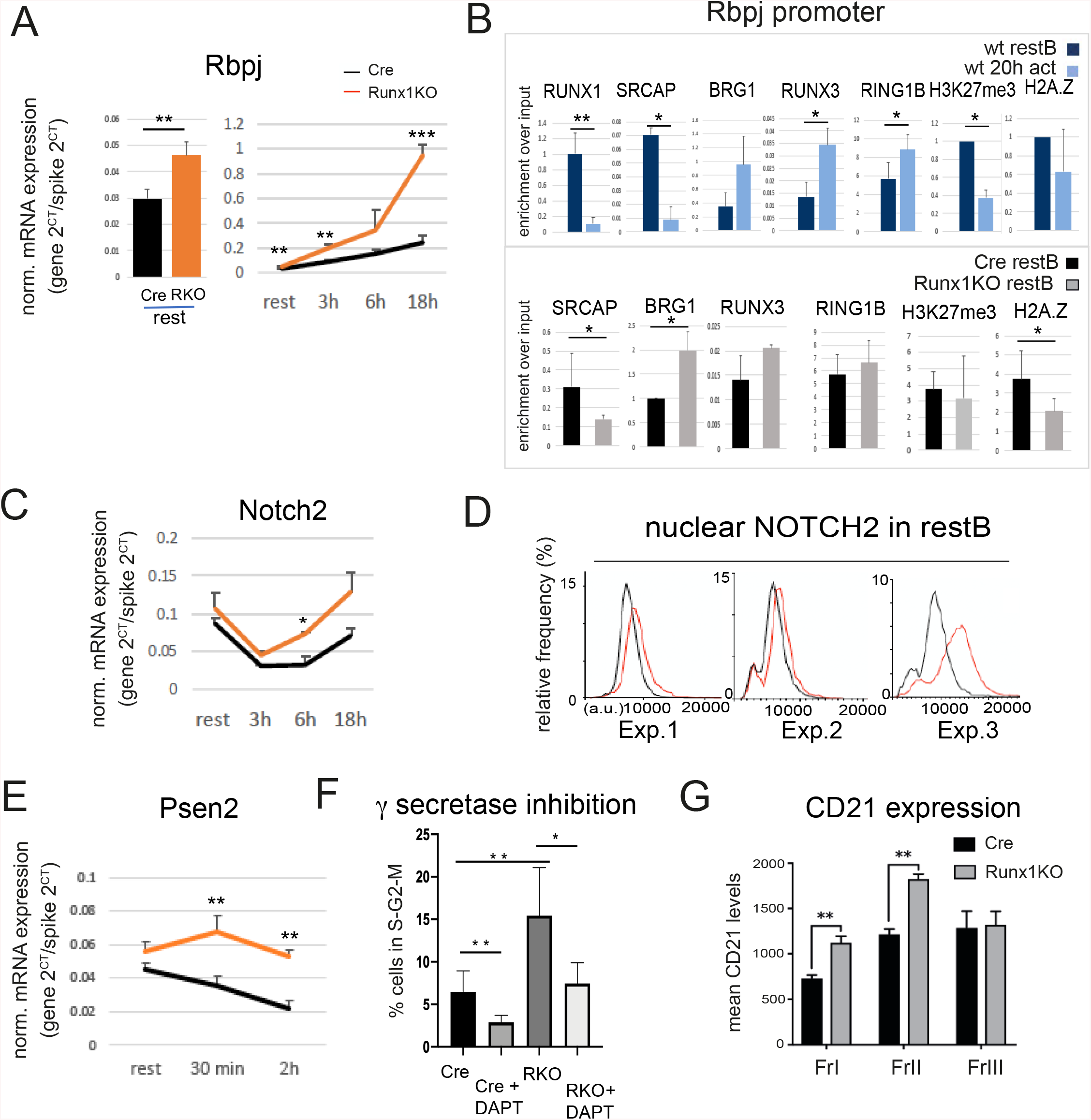
Runx1 represses Notch signaling in B cells. **A**. RT-qPCR analysis of *Rbpj* transcription in CD23-cre and Runx1 ck/o resting B cells (left panel) and upon activation with anti-IgM for 0-18 hrs (right panel). **B**. Top panel: ChIP-qPCR of wild-type (wt) resting B cells (dark blue) and after 20hrs anti-IgM activation (light blue) at the *Rbpj* promoter. Bottom panel: The same primers were used for ChIP-qPCR analysis of CD23-Cre (black) and Runx1 ck/o resting B cells (grey). For panels A, C and E, values are mean ± SD. * p≤0.5, ** p≤0.01, Student’s t-test, n>3. **C**. RT-qPCR of Notch2 transcript levels during a B cell activation time-course. **D**. Single cell analysis showing of the level of NOTCH2 protein in the nuclei of resting B cells (see Methods). The results of 3 independent experiments (Exp.1, 2 and 3) are shown. **E**. RT-qPCR of transcripts from *Psen2* after 30 min and 2hrs activation of Runx1 resting B cells with anti-IgM. **F**. Analysis of the effect of the γ-secretase inhibitor DAPT on entry of resting B cells into S-phase in response to 18 hrs incubation with anti-IgM followed by 6 hrs treatment with LPS. **G**. Mean CD21 surface levels on fraction I-III B cells from Cd23-Cre (black) and Runx1c-k/o/Cd23-Cre (grey) spleens. For panels F, G, values are mean ± SEM. ** p≤0.01, Students t-test, n=3.

The role of Notch signalling in the hyperresponsive phenotype was further tested by pre-incubating the wild-type and *Runx1* c-k/o cells with the γ-secretase inhibitor DAPT, followed by incubation with anti-IgM for 18 hours and LPS for 6 hours. The results showed that inhibiting the Notch pathway significantly reduced the proportion of the knockout cells that entered S-phase for both the cre and *Runx1* c-k/o B cells (Figure 5F). The γ-secretase inhibitor reduced S-phase entry of the knockout cells to levels that were similar to the untreated Cre cells, indicating that Notch signalling has an important role in the hyperresponsive phenotype of the *Runx1* c-k/o B cells. Consistent with Notch activation, we found that expression of CD21, which is encoded by the *Cr2* gene, is significantly upregulated in Runx1 K/O resting B cells (Figure 5G). It has been reported previously that ectopic expression of activated-NOTCH upregulates the BCR co-receptor CD21 in human B lymphoma cell lines (53).

## DISCUSSION

RUNX1 is known to be a critical regulator of gene expression during embryonic and adult haematopoiesis. The results described in this paper extend our knowledge of the functions of RUNX1 by showing that it plays a key role in regulating the level and rate of response of resting B cells to antigen through co-ordinated regulation of genes that affect cell cycle entry and BCR signalling. Conditional inactivation of *Runx1* in splenic resting B cells results in a phenotype of accelerated priming for cell cycle entry in response to BCR stimulation by anti-IgM. The RUNX1-regulated genes that we have identified can be grouped into three functional categories: (i) cell cycle and immediate early genes, (ii) immunomodulatory genes, many of which have been associated with autoimmune disease, and (iii) components of the Notch signalling pathway. All of the RUNX1 regulated genes are characterized by strong binding of RUNX1 to promoters and/or enhancers and the binding of the chromatin remodeller SRCAP shows a near absolute co-localization with binding peaks for RUNX1.

Analysis of the expression profiles of the cell cycle and immediate early genes that are affected by the knockout of *Runx1* revealed a rapid increase in transcript levels and enhancement of the rate at which transcription of the genes increases in response to BCR stimulation in the knockout cells. The conclusion that this indicates a role for RUNX1 in regulating poising of these genes is reinforced by the presence of high levels of RNA Pol II at their promoters in wild-type resting B cells and by the observation that they are bivalently marked with H3K4me3 and H3K27me3 histone modifications. Cyclin D2 has been shown to play an important role in B cell activation (28, 54) and ectopic overexpression of *Ccnd2* has been shown to induce post-mitotic cardiomyocytes to enter the cell cycle (55). Our data provide evidence that RUNX1-mediated downregulation of Cyclin D2 expression plays an important role in controlling the rate of entry of naïve B cells into S-phase.

Binding of RUNX1 to the poised cell cycle and immediate early genes occurs in conjunction with the chromatin remodeller SRCAP with particularly strong binding observed at distally located enhancer/silencer elements. BCR-mediated activation leads to a switch between binding of RUNX1 and its paralogue RUNX3 and between SRCAP and the SWI/SNF remodelling component BRG1. The hypothesis that RUNX1 recruits SRCAP to enhancers and promoters is supported by the strong co-localisation of SRCAP binding with RUNX1, the reduction in binding of SRCAP at the *Ccnd2* promoter in the *Runx1* c-k/o resting B cells and the fact that RUNX1 and SRCAP show a co-ordinated reduction in binding during B cell activation. It is noteworthy that RUNX3 has been reported to recruit BRG1 to gene promoters (47), suggesting that the switch from RUNX1 to RUNX3 binding plays a direct causal role in the SRCAP to BRG1 switch. In contrast, the knockout of RUNX1 does not significantly affect enrichment of the PRC2-dependent modification, H3K27me3, at the *Ccnd2* gene, although levels of H3K27me3 were strongly decreased at both genes following activation. Levels of the PRC1 protein RING1B were increased on the *Ccnd2* gene in anti-IgM treated wild-type cells and in the *Runx1* c-k/o resting B cells. These results indicate that that the PRC1 and PRC2 complexes are recruited independently of RUNX1 to the poised genes in B cells by as yet undetermined mechanisms. Binding of PRC1 independently of PRC2 has been shown to facilitate chromatin looping in mouse neural progenitor stem cells (41), so it is possible that increased binding of PRC1 at the *Ccnd2* gene during B cell activation acts to promote looping and stabilization of interactions between chromatin regulatory regions in the vicinity of the gene.

A second category of genes that are subject to RUNX1-mediated regulation are genes that are involved in the regulation of B cell signaling pathways or have known associations with autoimmune disease and are directly up or downregulated by RUNX1 in resting and early-stage activated B cells. These genes do not, in general, show evidence of poising and the major effect of RUNX1 binding is to increase or decrease the levels at which they are transcribed, before and immediately after activation. Of particular interest is the repressive effect of RUNX1 on the type I interferon response. Dysregulation of type 1 interferon signaling has been strongly associated with lupus and has also been observed in Sjogren’s syndrome and systemic sclerosis (reviewed in (56). Our results support the idea that RUNX1 has an important role in downregulating the interferon response and BCR signaling in naïve resting B cells, and that this forms part of a potential mechanism by which RUNX1 is involved in regulating B cell tolerance.

Components of the NOTCH signalling pathway form a third category of genes that are affected by RUNX1. Expression of the Notch effector, RBPJ, which binds to gene promoters in conjunction with activated Notch, was upregulated during activation of *Runx1* knockout B cells, similar to *Ccnd2*, and expression of the γ-secretase catalytic subunit, *Psen2* was also strongly upregulated by the *Runx1* knockout in resting B cells. This was reflected in the increase in activated nuclear Notch2 protein that was observed in *Runx1* c-k/o resting B cells. The involvement of the Notch pathway in the modulating effect of Runx1 on cell cycle entry was confirmed by the observation that premature entry of the *Runx1* knockout B cells into S-phase was reduced by treatment with a Notch inhibitor.

Taken together, our results provide evidence that RUNX1 acts as a major regulator of B cell activation and antigen responsiveness by directly affecting the expression of genes that affect the rate of cell cycle entry and by regulating expression of immunomodulators during the early stages of activation in response to antigen. These effects are shown schematically in Figure 6. Regulatory single nucleotide polymorphisms (rSNPs) that affect binding of RUNX1 have been identified in genes that are associated with rheumatoid arthritis (57) and lupus (58). The findings of this study suggest that changes to the expression and functioning of RUNX1 in B cells can have a causal role in human autoimmune disease.

**Figure 6.**
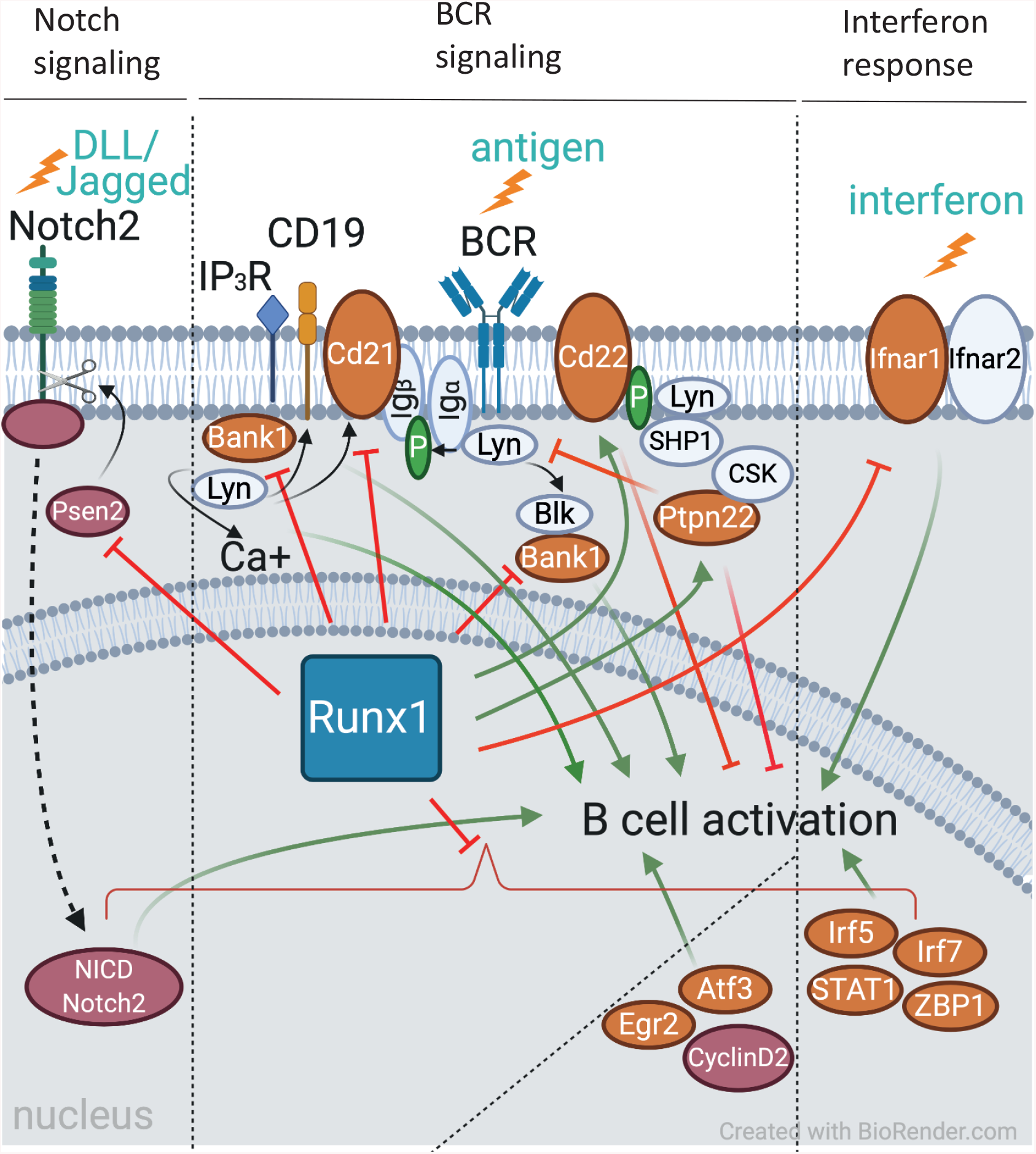
Genes that are implicated in B cell activation and autoimmunity and are regulated by RUNX1. A schematic representation is shown of the involvement in B cell pathways and in autoimmune disease, of proteins encoded by genes shown to be regulated by RUNX1 in this study. Red lines indicate negative regulation and green arrows represent positive regulation, either through direct effects of RUNX1 on gene expression, or as a consequence of downstream regulation of signaling and B cell activation by regulatory targets of RUNX1. Orange filled ovals indicate proteins that have been implicated in autoimmune disease.

## ACKNOWLEDGEMENTS

We thank Nancy Speck for providing the conditional *Runx1* knockout mouse line and Meinrad Busslinger for the CD23-cre transgenic mouse line. We also thank Tom Carroll for assistance with the initial stages of the bioinformatic analysis. The work was supported by the LMS/NIHR Imperial Biomedical Research Centre Flow Cytometry Facility. This research was funded by the Medical Research Council UK.

## REFERENCES

1. Imperato, M. R., P. Cauchy, N. Obier, and C. Bonifer. 2015. The RUNX1-PU.1 axis in the control of hematopoiesis. International journal of hematology 101: 319–329.

2. Mevel, R., J. E. Draper, A. L. M. Lie, V. Kouskoff, and G. Lacaud. 2019. RUNX transcription factors: orchestrators of development. Development 146.

3. Ichikawa, M., T. Asai, T. Saito, S. Seo, I. Yamazaki, T. Yamagata, K. Mitani, S. Chiba, S. Ogawa, M. Kurokawa, and H. Hirai. 2004. AML-1 is required for megakaryocytic maturation and lymphocytic differentiation, but not for maintenance of hematopoietic stem cells in adult hematopoiesis. Nat Med 10: 299–304.

4. Wang, Q., T. Stacy, M. Binder, M. Marin-Padilla, A. H. Sharpe, and N. A. Speck. 1996. Disruption of the Cbfa2 gene causes necrosis and hemorrhaging in the central nervous system and blocks definitive hematopoiesis. Proc Natl Acad Sci U S A 93: 3444–3449.

5. Seo, W., and I. Taniuchi. 2020. The Roles of RUNX Family Proteins in Development of Immune Cells. Mol Cells 43: 107–113.

6. Hong, D., A. J. Fritz, J. A. Gordon, C. E. Tye, J. R. Boyd, K. M. Tracy, S. E. Frietze, F. E. Carr, J. A. Nickerson, A. J. Van Wijnen, A. N. Imbalzano, S. K. Zaidi, J. B. Lian, J. L. Stein, and G. S. Stein. 2019. RUNX1-dependent mechanisms in biological control and dysregulation in cancer. J Cell Physiol 234: 8597–8609.

7. Ito, Y., S. C. Bae, and L. S. Chuang. 2015. The RUNX family: developmental regulators in cancer. Nature reviews. Cancer 15: 81–95.

8. Sood, R., Y. Kamikubo, and P. Liu. 2017. Role of RUNX1 in hematological malignancies. Blood 129: 2070–2082.

9. Yan, M., E. Kanbe, L. F. Peterson, A. Boyapati, Y. Miao, Y. Wang, I. M. Chen, Z. Chen, J. D. Rowley, C. L. Willman, and D. E. Zhang. 2006. A previously unidentified alternatively spliced isoform of t(8;21) transcript promotes leukemogenesis. Nat Med 12: 945–949.

10. Pui, C. H., M. V. Relling, and J. R. Downing. 2004. Acute lymphoblastic leukemia. The New England journal of medicine 350: 1535–1548.

11. Bhojwani, D., D. Pei, J. T. Sandlund, S. Jeha, R. C. Ribeiro, J. E. Rubnitz, S. C. Raimondi, S. Shurtleff, M. Onciu, C. Cheng, E. Coustan-Smith, W. P. Bowman, S. C. Howard, M. L. Metzger, H. Inaba, W. Leung, W. E. Evans, D. Campana, M. V. Relling, and C. H. Pui. 2012. ETV6-RUNX1-positive childhood acute lymphoblastic leukemia: improved outcome with contemporary therapy. Leukemia 26: 265–270.

12. Maier, H., R. Ostraat, H. Gao, S. Fields, S. A. Shinton, K. L. Medina, T. Ikawa, C. Murre, H. Singh, R. R. Hardy, and J. Hagman. 2004. Early B cell factor cooperates with Runx1 and mediates epigenetic changes associated with mb-1 transcription. Nat Immunol 5: 1069–1077.

13. Seo, W., T. Ikawa, H. Kawamoto, and I. Taniuchi. 2012. Runx1-Cbfbeta facilitates early B lymphocyte development by regulating expression of Ebf1. The Journal of experimental medicine 209: 1255–1262.

14. Gunnell, A., H. M. Webb, C. D. Wood, M. J. McClellan, B. Wichaidit, B. Kempkes, R. G. Jenner, C. Osborne, P. J. Farrell, and M. J. West. 2016. RUNX super-enhancer control through the Notch pathway by Epstein-Barr virus transcription factors regulates B cell growth. Nucleic Acids Res 44: 4636–4650.

15. Brady, G., C. Elgueta Karstegl, and P. J. Farrell. 2013. Novel function of the unique N-terminal region of RUNX1c in B cell growth regulation. Nucleic Acids Res 41: 1555–1568.

16. Growney, J. D., H. Shigematsu, Z. Li, B. H. Lee, J. Adelsperger, R. Rowan, D. P. Curley, J. L. Kutok, K. Akashi, I. R. Williams, N. A. Speck, and D. G. Gilliland. 2005. Loss of Runx1 perturbs adult hematopoiesis and is associated with a myeloproliferative phenotype. Blood 106: 494–504.

17. Kwon, K., C. Hutter, Q. Sun, I. Bilic, C. Cobaleda, S. Malin, and M. Busslinger. 2008. Instructive role of the transcription factor E2A in early B lymphopoiesis and germinal center B cell development. Immunity 28: 751–762.

18. Sabbattini, P., M. Sjoberg, S. Nikic, A. Frangini, P. H. Holmqvist, N. Kunowska, T. Carroll, E. Brookes, S. J. Arthur, A. Pombo, and N. Dillon. 2014. An H3K9/S10 methyl-phospho switch modulates Polycomb and Pol II binding at repressed genes during differentiation. Mol Biol Cell 25: 904–915.

19. Kouzine, F., D. Wojtowicz, A. Yamane, W. Resch, K. R. Kieffer-Kwon, R. Bandle, S. Nelson, H. Nakahashi, P. Awasthi, L. Feigenbaum, H. Menoni, J. Hoeijmakers, W. Vermeulen, H. Ge, T. M. Przytycka, D. Levens, and R. Casellas. 2013. Global regulation of promoter melting in naive lymphocytes. Cell 153: 988–999.

20. Sabbattini, P., C. Canzonetta, M. Sjoberg, S. Nikic, A. Georgiou, G. Kemball-Cook, H. W. Auner, and N. Dillon. 2007. A novel role for the Aurora B kinase in epigenetic marking of silent chromatin in differentiated postmitotic cells. Embo J 26: 4657–4669.

21. Hardy, R. R., K. Hayakawa, D. R. Parks, and L. A. Herzenberg. 1983. Demonstration of B-cell maturation in X-linked immunodeficient mice by simultaneous three-colour immunofluorescence. Nature 306: 270–272.

22. Cariappa, A., C. Boboila, S. T. Moran, H. Liu, H. N. Shi, and S. Pillai. 2007. The recirculating B cell pool contains two functionally distinct, long-lived, posttransitional, follicular B cell populations. J Immunol 179: 2270–2281.

23. Watanabe, K., M. Sugai, Y. Nambu, M. Osato, T. Hayashi, M. Kawaguchi, T. Komori, Y. Ito, and A. Shimizu. 2010. Requirement for Runx proteins in IgA class switching acting downstream of TGF-beta 1 and retinoic acid signaling. J Immunol 184: 2785–2792.

24. Moir, S., and A. S. Fauci. 2009. B cells in HIV infection and disease. Nature reviews. Immunology 9: 235–245.

25. Meyer-Bahlburg, A., A. D. Bandaranayake, S. F. Andrews, and D. J. Rawlings. 2009. Reduced c-myc expression levels limit follicular mature B cell cycling in response to TLR signals. J Immunol 182: 4065–4075.

26. Defranco, A. L., E. S. Raveche, R. Asofsky, and W. E. Paul. 1982. Frequency of B lymphocytes responsive to anti-immunoglobulin. The Journal of experimental medicine 155: 1523–1536.

27. Meyer-Bahlburg, A., S. Khim, and D. J. Rawlings. 2007. B cell intrinsic TLR signals amplify but are not required for humoral immunity. The Journal of experimental medicine 204: 3095–3101.

28. Solvason, N., W. W. Wu, D. Parry, D. Mahony, E. W. Lam, J. Glassford, G. G. Klaus, P. Sicinski, R. Weinberg, Y. J. Liu, M. Howard, and E. Lees. 2000. Cyclin D2 is essential for BCR-mediated proliferation and CD5 B cell development. Int Immunol 12: 631–638.

29. Perez-Roger, I., D. L. Solomon, A. Sewing, and H. Land. 1997. Myc activation of cyclin E/Cdk2 kinase involves induction of cyclin E gene transcription and inhibition of p27(Kip1) binding to newly formed complexes. Oncogene 14: 2373–2381.

30. Johnson, D. G., K. Ohtani, and J. R. Nevins. 1994. Autoregulatory control of E2F1 expression in response to positive and negative regulators of cell cycle progression. Genes Dev 8: 1514–1525.

31. Andersson, R., and A. Sandelin. 2020. Determinants of enhancer and promoter activities of regulatory elements. Nature reviews 21: 71–87.

32. Lichtinger, M., R. Ingram, R. Hannah, D. Muller, D. Clarke, S. A. Assi, A. L. M. Lie, L. Noailles, M. S. Vijayabaskar, M. Wu, D. G. Tenen, D. R. Westhead, V. Kouskoff, G. Lacaud, B. Gottgens, and C. Bonifer. 2012. RUNX1 reshapes the epigenetic landscape at the onset of haematopoiesis. EMBO J 31: 4318–4333.

33. Wan, M., J. Zhang, D. Lai, A. Jani, P. Prestone-Hurlburt, L. Zhao, A. Ramachandran, G. Schnitzler, and T. Chi. 2009. Molecular basis of CD4 repression by the Swi/Snf-like BAF chromatin remodeling complex. Eur J Immunol 39: 580–588.

34. Wong, M. M., L. K. Cox, and J. C. Chrivia. 2007. The chromatin remodeling protein, SRCAP, is critical for deposition of the histone variant H2A.Z at promoters. J Biol Chem 282: 26132–26139.

35. Azuara, V., P. Perry, S. Sauer, M. Spivakov, H. F. Jorgensen, R. M. John, M. Gouti, M. Casanova, G. Warnes, M. Merkenschlager, and A. G. Fisher. 2006. Chromatin signatures of pluripotent cell lines. Nat Cell Biol 8: 532–538.

36. Bernstein, B. E., T. S. Mikkelsen, X. Xie, M. Kamal, D. J. Huebert, J. Cuff, B. Fry, A. Meissner, M. Wernig, K. Plath, R. Jaenisch, A. Wagschal, R. Feil, S. L. Schreiber, and E. Lander. 2006. A bivalent chromatin structure marks key developmental genes in embryonic stem cells. Cell 125: 315–326.

37. Schwartz, Y. B., and V. Pirrotta. 2013. A new world of Polycombs: unexpected partnerships and emerging functions. Nature reviews 14: 853–864.

38. Brookes, E., I. De Santiago, D. Hebenstreit, Kelly J. Morris, T. Carroll, Sheila Q. Xie, Julie K. Stock, M. Heidemann, D. Eick, N. Nozaki, H. Kimura, J. Ragoussis, Sarah A. Teichmann, and A. Pombo. 2012. Polycomb Associates Genome-wide with a Specific RNA Polymerase II Variant, and Regulates Metabolic Genes in ESCs. Cell stem cell 10: 157–170.

39. Frangini, A., M. Sjöberg, M. Roman-Trufero, G. Dharmalingam, V. Haberle, T. Bartke, B. Lenhard, M. Malumbres, M. Vidal, and N. Dillon. 2013. The Aurora B Kinase and the Polycomb Protein Ring1B Combine to Regulate Active Promoters in Quiescent Lymphocytes. Molecular Cell 51: 647–661.

40. An, Z., B. Akily, M. Sabalic, G. Zong, Y. Chai, and P. T. Sharpe. 2018. Regulation of Mesenchymal Stem to Transit-Amplifying Cell Transition in the Continuously Growing Mouse Incisor. Cell reports 23: 3102–3111.

41. Loubiere, V., G. L. Papadopoulos, Q. Szabo, A. M. Martinez, and G. Cavalli. 2020. Widespread activation of developmental gene expression characterized by PRC1-dependent chromatin looping. Sci Adv 6: eaax4001.

42. Yu, M., T. Mazor, H. Huang, H.-T. Huang, Katie L. Kathrein, Andrew J. Woo, Candace R. Chouinard, A. Labadorf, Thomas E. Akie, Tyler B. Moran, H. Xie, S. Zacharek, I. Taniuchi, Robert G. Roeder, Carla F. Kim, Leonard I. Zon, E. Fraenkel, and Alan B. Cantor. 2012. Direct Recruitment of Polycomb Repressive Complex 1 to Chromatin by Core Binding Transcription Factors. Molecular Cell 45: 330–343.

43. Muse, G. W., D. A. Gilchrist, S. Nechaev, R. Shah, J. S. Parker, S. F. Grissom, J. Zeitlinger, and K. Adelman. 2007. RNA polymerase is poised for activation across the genome. Nat Genet 39: 1507–1511.

44. Stock, J. K., S. Giadrossi, M. Casanova, E. Brookes, M. Vidal, H. Koseki, N. Brockdorff, A. G. Fisher, and A. Pombo. 2007. Ring1-mediated ubiquitination of H2A restrains poised RNA polymerase II at bivalent genes in mouse ES cells. Nat Cell Biol 9: 1428–1435.

45. Martinez-Soria, N., L. McKenzie, J. Draper, A. Ptasinska, H. Issa, S. Potluri, H. J. Blair, A. Pickin, A. Isa, P. S. Chin, R. Tirtakusuma, D. Coleman, S. Nakjang, S. Assi, V. Forster, M. Reza, E. Law, P. Berry, D. Mueller, C. Osborne, A. Elder, S. N. Bomken, D. Pal, J. M. Allan, G. J. Veal, P. N. Cockerill, C. Wichmann, J. Vormoor, G. Lacaud, C. Bonifer, and O. Heidenreich. 2018. The Oncogenic Transcription Factor RUNX1/ETO Corrupts Cell Cycle Regulation to Drive Leukemic Transformation. Cancer Cell 34: 626–642 e628.

46. Spender, L. C., G. H. Cornish, A. Sullivan, and P. J. Farrell. 2002. Expression of transcription factor AML-2 (RUNX3, CBF(alpha)-3) is induced by Epstein-Barr virus EBNA-2 and correlates with the B-cell activation phenotype. J Virol 76: 4919–4927.

47. Lee, J. W., D. M. Kim, J. W. Jang, T. G. Park, S. H. Song, Y. S. Lee, X. Z. Chi, I. Y. Park, J. W. Hyun, Y. Ito, and S. C. Bae. 2019. RUNX3 regulates cell cycle-dependent chromatin dynamics by functioning as a pioneer factor of the restriction-point. Nat Commun 10: 1897.

48. Kopan, R., and M. X. Ilagan. 2009. The canonical Notch signaling pathway: unfolding the activation mechanism. Cell 137: 216–233.

49. Radtke, F., H. R. MacDonald, and F. Tacchini-Cottier. 2013. Regulation of innate and adaptive immunity by Notch. Nature reviews. Immunology 13: 427–437.

50. Saito, T., S. Chiba, M. Ichikawa, A. Kunisato, T. Asai, K. Shimizu, T. Yamaguchi, G. Yamamoto, S. Seo, K. Kumano, E. Nakagami-Yamaguchi, Y. Hamada, S. Aizawa, and H. Hirai. 2003. Notch2 is preferentially expressed in mature B cells and indispensable for marginal zone B lineage development. Immunity 18: 675–685.

51. Tanigaki, K., H. Han, N. Yamamoto, K. Tashiro, M. Ikegawa, K. Kuroda, A. Suzuki, T. Nakano, and T. Honjo. 2002. Notch-RBP-J signaling is involved in cell fate determination of marginal zone B cells. Nat Immunol 3: 443–450.

52. Thomas, M., M. Calamito, B. Srivastava, I. Maillard, W. S. Pear, and D. Allman. 2007. Notch activity synergizes with B-cell-receptor and CD40 signaling to enhance B-cell activation. Blood 109: 3342–3350.

53. Strobl, L. J., H. Hofelmayr, G. Marschall, M. Brielmeier, G. W. Bornkamm, and U. Zimber-Strobl. 2000. Activated Notch1 modulates gene expression in B cells similarly to Epstein-Barr viral nuclear antigen 2. J Virol 74: 1727–1735.

54. Piatelli, M. J., C. Doughty, and T. C. Chiles. 2002. Requirement for a hsp90 chaperone-dependent MEK1/2-ERK pathway for B cell antigen receptor-induced cyclin D2 expression in mature B lymphocytes. J Biol Chem 277: 12144–12150.

55. Busk, P. K., R. Hinrichsen, J. Bartkova, A. H. Hansen, T. E. Christoffersen, J. Bartek, and S. Haunso. 2005. Cyclin D2 induces proliferation of cardiac myocytes and represses hypertrophy. Exp Cell Res 304: 149–161.

56. Crow, M. K., M. Olferiev, and K. A. Kirou. 2019. Type I Interferons in Autoimmune Disease. Annual review of pathology 14: 369–393.

57. Tokuhiro, S., R. Yamada, X. Chang, A. Suzuki, Y. Kochi, T. Sawada, M. Suzuki, M. Nagasaki, M. Ohtsuki, M. Ono, H. Furukawa, M. Nagashima, S. Yoshino, A. Mabuchi, A. Sekine, S. Saito, A. Takahashi, T. Tsunoda, Y. Nakamura, and K. Yamamoto. 2003. An intronic SNP in a RUNX1 binding site of SLC22A4, encoding an organic cation transporter, is associated with rheumatoid arthritis. Nat Genet 35: 341–348.

58. Prokunina, L., C. Castillejo-Lopez, F. Oberg, I. Gunnarsson, L. Berg, V. Magnusson, A. J. Brookes, D. Tentler, H. Kristjansdottir, G. Grondal, A. I. Bolstad, E. Svenungsson, I. Lundberg, G. Sturfelt, A. Jonssen, L. Truedsson, G. Lima, J. Alcocer-Varela, R. Jonsson, U. B. Gyllensten, J. B. Harley, D. Alarcon-Segovia, K. Steinsson, and M. E. Alarcon-Riquelme. 2002. A regulatory polymorphism in PDCD1 is associated with susceptibility to systemic lupus erythematosus in humans. Nat Genet 32: 666–669.

59. Menard, L., D. Saadoun, I. Isnardi, Y. S. Ng, G. Meyers, C. Massad, C. Price, C. Abraham, R. Motaghedi, J. H. Buckner, P. K. Gregersen, and E. Meffre. 2011. The PTPN22 allele encoding an R620W variant interferes with the removal of developing autoreactive B cells in humans. J Clin Invest 121: 3635–3644.

60. Cook, D. A., G. T. Kannarkat, A. F. Cintron, L. M. Butkovich, K. B. Fraser, J. Chang, N. Grigoryan, S. A. Factor, A. B. West, J. M. Boss, and M. G. Tansey. 2017. LRRK2 levels in immune cells are increased in Parkinson’s disease. NPJ Parkinsons Dis 3: 11.

61. Zhang, M., C. Yao, J. Cai, S. Liu, X. N. Liu, Y. Chen, S. Wang, P. Ji, M. Pan, Z. Kang, and Y. Wang. 2019. LRRK2 is involved in the pathogenesis of system lupus erythematosus through promoting pathogenic antibody production. J Transl Med 17: 37.

62. Varona, R., V. Cadenas, J. Flores, A. C. Martinez, and G. Marquez. 2003. CCR6 has a non-redundant role in the development of inflammatory bowel disease. Eur J Immunol 33: 2937–2946.

63. O’Keefe, T. L., G. T. Williams, S. L. Davies, and M. S. Neuberger. 1996. Hyperresponsive B cells in CD22-deficient mice. Science 274: 798–801.

64. Muller, J., and L. Nitschke. 2014. The role of CD22 and Siglec-G in B-cell tolerance and autoimmune disease. Nat Rev Rheumatol 10: 422–428.

65. Shi, L., Z. Bian, C. X. Chen, Y. N. Guo, Z. Lv, C. Zeng, Z. Liu, K. Zen, and Y. Liu. 2015. CD47 deficiency ameliorates autoimmune nephritis in Fas(lpr) mice by suppressing IgG autoantibody production. J Pathol 237: 285–295.

66. Ning, S., J. S. Pagano, and G. N. Barber. 2011. IRF7: activation, regulation, modification and function. Genes Immun 12: 399–414.

67. Ban, T., G. R. Sato, and T. Tamura. 2018. Regulation and role of the transcription factor IRF5 in innate immune responses and systemic lupus erythematosus. Int Immunol 30: 529–536.

68. Domeier, P. P., S. B. Chodisetti, S. L. Schell, Y. I. Kawasawa, M. J. Fasnacht, C. Soni, and Z. S. M. Rahman. 2018. B-Cell-Intrinsic Type 1 Interferon Signaling Is Crucial for Loss of Tolerance and the Development of Autoreactive B Cells. Cell reports 24: 406–418.

69. Hagberg, N., and L. Ronnblom. 2015. Systemic Lupus Erythematosus--A Disease with A Dysregulated Type I Interferon System. Scand J Immunol 82: 199–207.

70. Wahadat, M. J., I. L. A. Bodewes, N. I. Maria, C. G. van Helden-Meeuwsen, A. van Dijk-Hummelman, E. C. Steenwijk, S. Kamphuis, and M. A. Versnel. 2018. Type I IFN signature in childhood-onset systemic lupus erythematosus: a conspiracy of DNA- and RNA-sensing receptors? Arthritis research & therapy 20: 4.

71. Zhang, W., Q. Zhou, W. Xu, Y. Cai, Z. Yin, X. Gao, and S. Xiong. 2013. DNA-dependent activator of interferon-regulatory factors (DAI) promotes lupus nephritis by activating the calcium pathway. J Biol Chem 288: 13534–13550.

72. Simoni, L., V. Delgado, J. Ruer-Laventie, D. Bouis, A. Soley, V. Heyer, I. Robert, V. Gies, T. Martin, A. S. Korganow, B. Reina-San-Martin, and P. Soulas-Sprauel. 2018. Trib1 Is Overexpressed in Systemic Lupus Erythematosus, While It Regulates Immunoglobulin Production in Murine B Cells. Front Immunol 9: 373.

73. Zeggar, S., K. S. Watanabe, S. Teshigawara, S. Hiramatsu, T. Katsuyama, E. Katsuyama, H. Watanabe, Y. Matsumoto, T. Kawabata, K. E. Sada, T. Niki, M. Hirashima, and J. Wada. 2018. Role of Lgals9 Deficiency in Attenuating Nephritis and Arthritis in BALB/c Mice in a Pristane-Induced Lupus Model. Arthritis Rheumatol 70: 1089–1101.

74. Aue, A., F. Szelinski, S. Y. Weissenberg, A. Wiedemann, T. Rose, A. C. Lino, and T. Dorner. 2020. Elevated STAT1 expression but not phosphorylation in lupus B cells correlates with disease activity and increased plasmablast susceptibility. Rheumatology (Oxford).

75. Neusser, M. A., M. T. Lindenmeyer, I. Edenhofer, S. Gaiser, M. Kretzler, H. Regele, S. Segerer, and C. D. Cohen. 2011. Intrarenal production of B-cell survival factors in human lupus nephritis. Mod Pathol 24: 98–107.

76. Du, S. W., H. M. Jacobs, T. Arkatkar, D. J. Rawlings, and S. W. Jackson. 2018. Integrated B Cell, Toll-like, and BAFF Receptor Signals Promote Autoantibody Production by Transitional B Cells. J Immunol 201: 3258–3268.

77. Kozyrev, S. V., A. K. Abelson, J. Wojcik, A. Zaghlool, M. V. Linga Reddy, E. Sanchez, I. Gunnarsson, E. Svenungsson, G. Sturfelt, A. Jonsen, L. Truedsson, B. A. Pons-Estel, T. Witte, S. D’Alfonso, N. Barizzone, M. G. Danieli, C. Gutierrez, A. Suarez, P. Junker, H. Laustrup, M. F. Gonzalez-Escribano, J. Martin, H. Abderrahim, and M. E. Alarcon-Riquelme. 2008. Functional variants in the B-cell gene BANK1 are associated with systemic lupus erythematosus. Nat Genet 40: 211–216.

