## Supplementary information for "RUNX1 controls the dynamics of cell cycle entry of naïve resting B cells by regulating expression of cell cycle and immunomodulatory genes in response to BCR stimulation"

**Supplementary Figure 1**

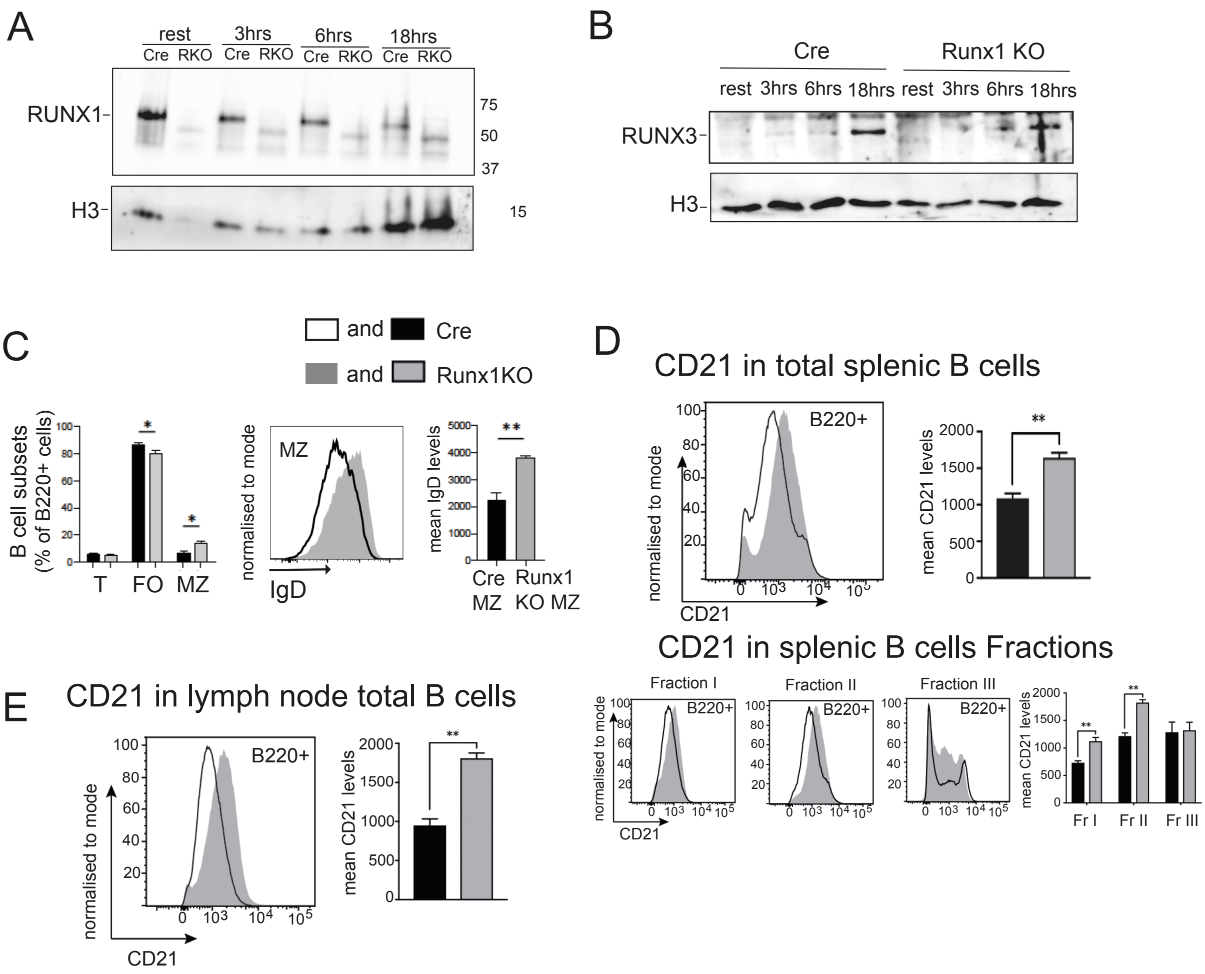


**Supplementary Figure 1**. **A, B.** Switch in RUNX paralog expression following B cell activation. Western Blot analysis of the levels of RUNX1 (**A**) and RUNX3 (**B**) in *Runx1* c-k/o (RKO) and CD23-cre (Cre) control resting B cells after activation with anti-IgM for 0-18hrs. **C**. Left panel; percentages of Transitional (T – CD23low/CD21low), Follicular (FO – CD23high/CD21mid-high) and Marginal Zone (MZ – CD23low/CD21high) B cells in splenic B220$^{+}$ cell populations. Central and right panels: Histogram and mean of IgD levels on the surface of MZ cells. Statistical calculations for all panels in this figure: error bars represent SEM. * p≤0.05, ** p≤0.01, Student's t test. n=3. **D**: Runx1 c-k/o splenic B cells express higher levels of surface CD21. Top left panel: representative histogram of CD21 surface levels on B220+ splenic B cells. Y-axis represents frequency of the CD21 level on the X-axis shown as a percentage of maximum count. Top right panel: mean

CD21 surface levels on splenic B220+ cells. Bottom right panels: representative histograms of CD21 surface levels in Fraction I-III cells from B220+ splenic B cells. Bottom left panel: mean CD21 surface levels on Fraction I-III spenic B cells. **E**. Left panel: Representative histograms of CD21 level as a percentage of maximum count. Right panel: mean CD21 surface levels on B220+ lymph node B cells.

#### Supplementary Figure 2

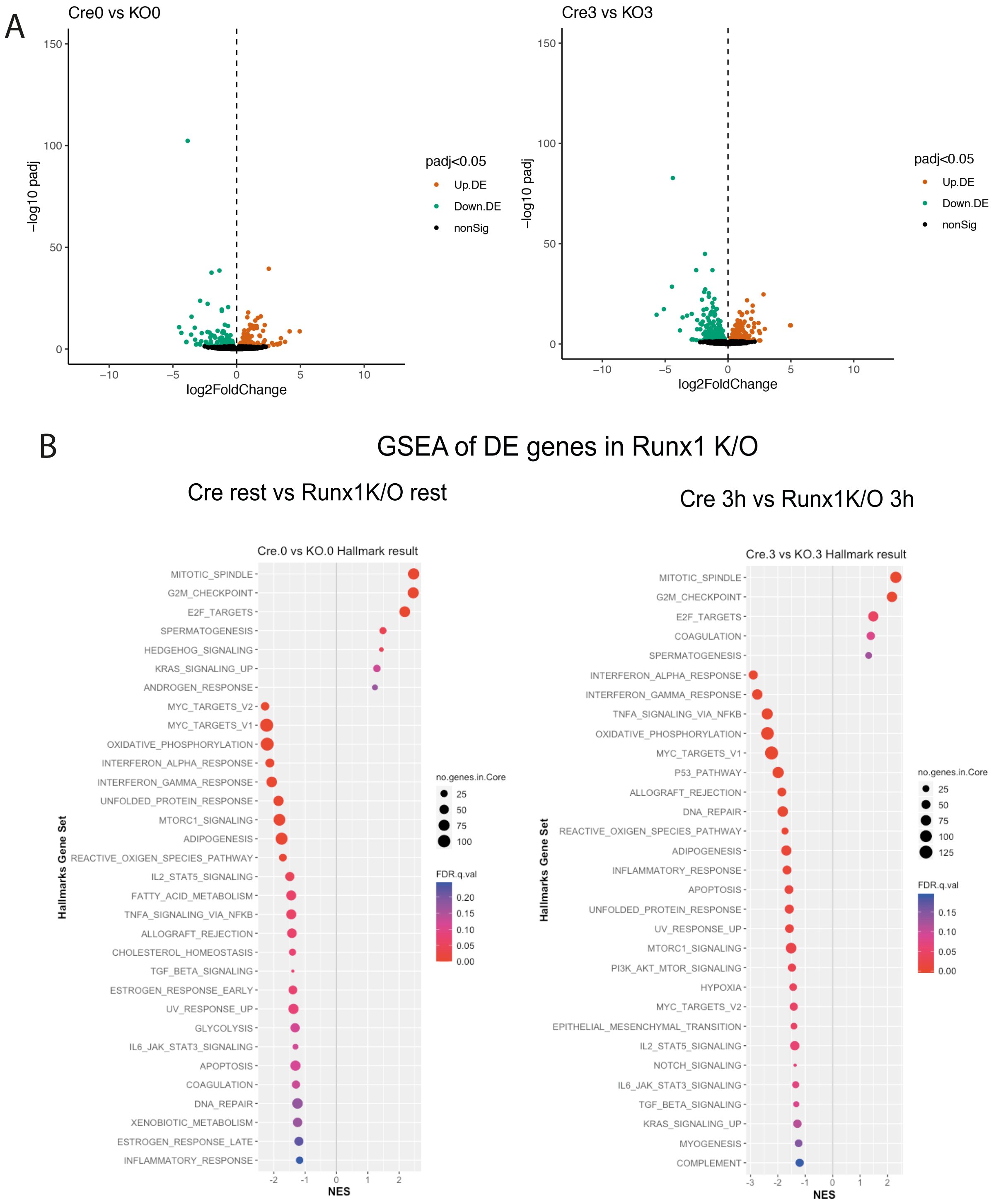


**Supplementary Figure 2. A.** 'Volcano plot' of statistical significance (adjusted p value) against fold- change of differentially expressed (DE) genes in Cre vs Runx1 ck/o B cell at the resting state (left panels) or after 3 hrs of activation with anti-IgM. **B.** Scatter plots illustrate enriched Runx1 c-k/o database pathways in resting B cells (left panel) and in

3hrs with anti-IgM activated cells (right panel). The vertical axis represents the enriched pathway categories and the horizontal axis represents the NES = Normalised Enrichment Score of the enriched pathways. The size and colour of dots represent the gene number and the range of p-values, respectively. The Normalised Enrichment Score (NES) is the ratio of differentially expressed gene number enriched in the pathway to the total gene number in a certain pathway. The enriched pathways with negative values represent pathways that are upregulated in the Runx1 c-k/o B cells, as they are expressed at a lower level in the CD23-cre control than in the Runx1 c-k/o cells.

### Supplementary Figure 3

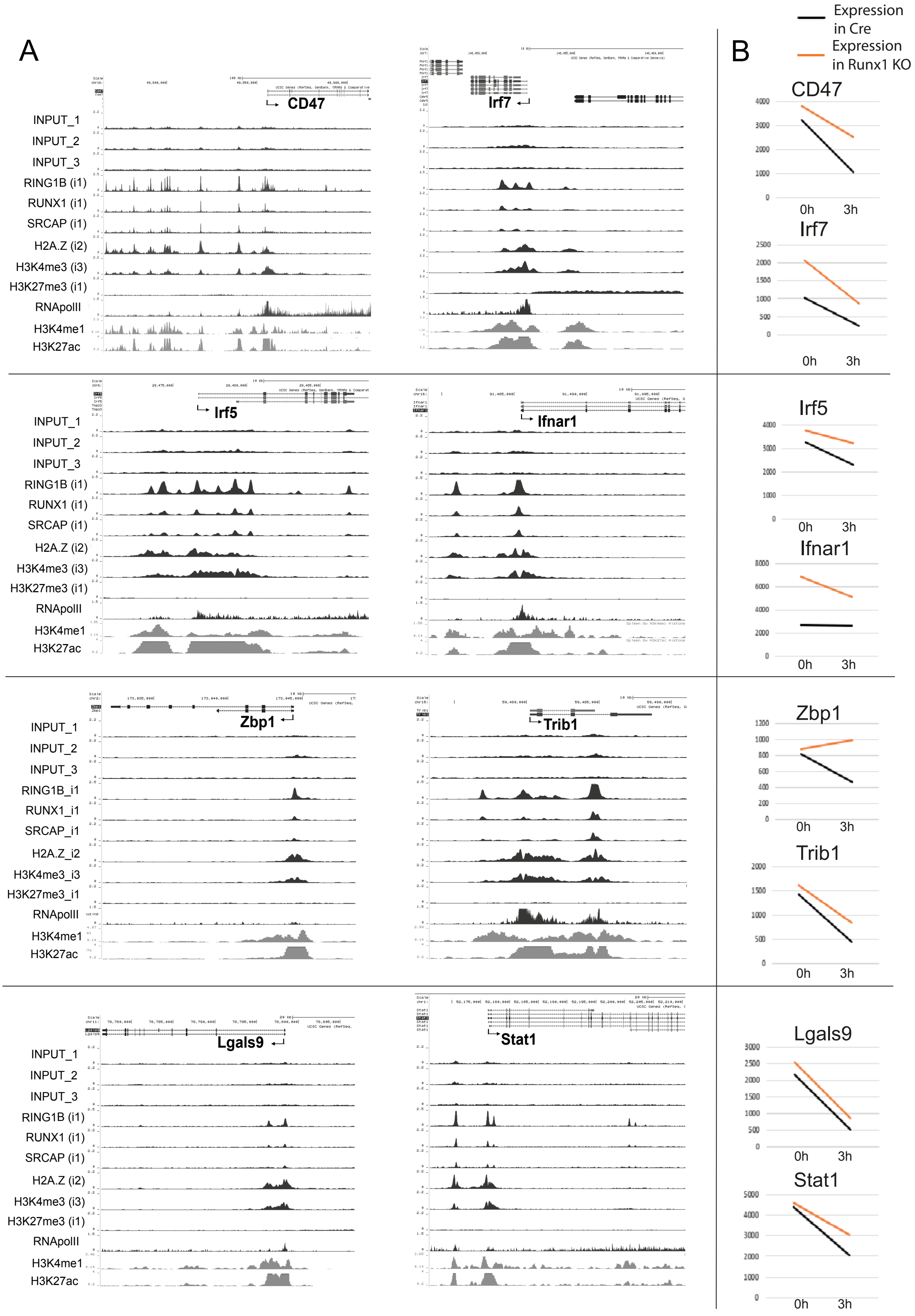

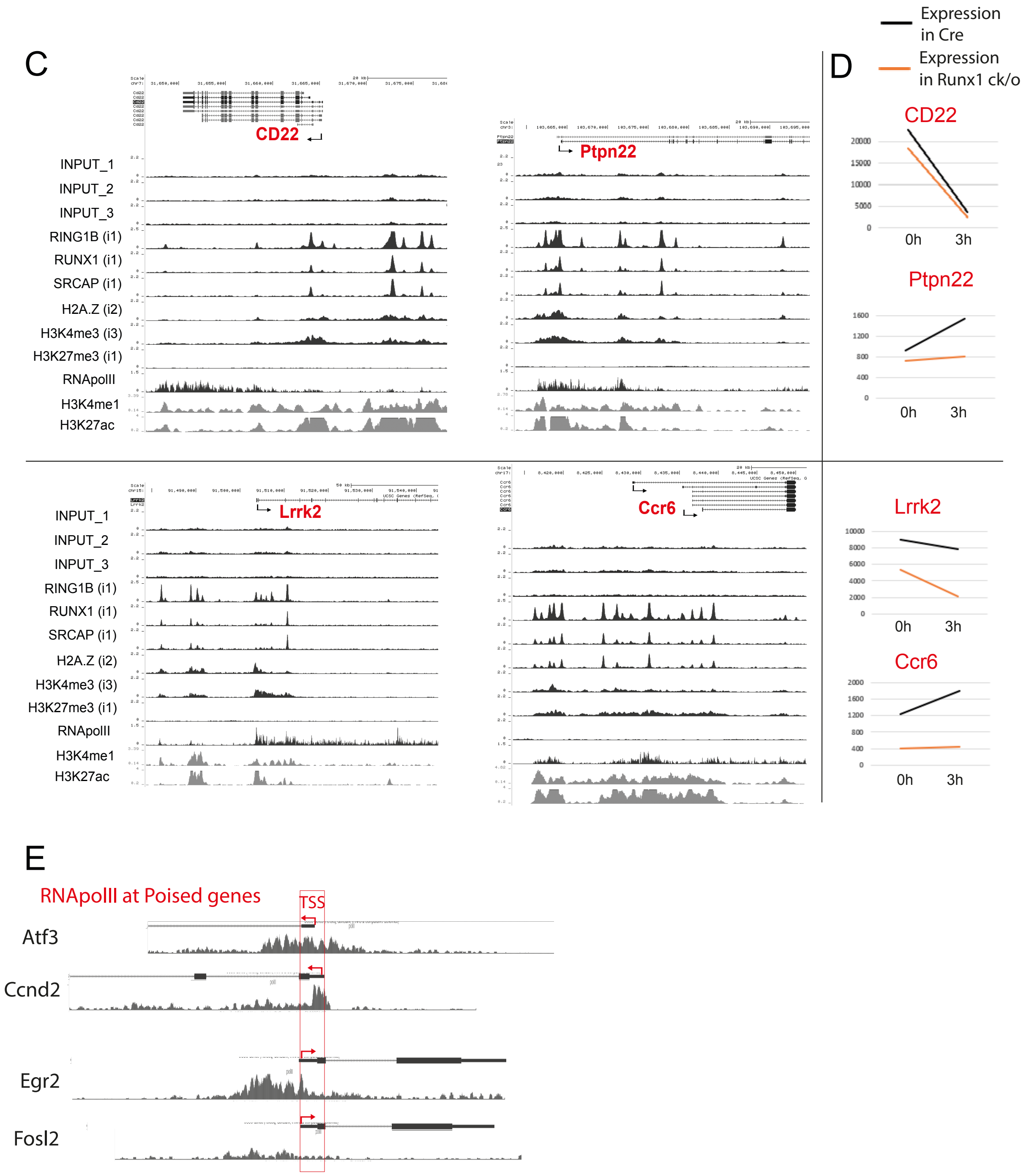


**Supplementary Figure 3**: **A**. ChIPseq tracks in resting B cells from wild-type mice in resting B (0h) and after 3hrs of activation with anti-IgM (3h) for active genes involved in autoimmunity that are upregulated in the Runx1 c-k/o resting B cells (Table 1). **B.** Basemean values for gene expression for the genes shown in (A) obtained from the RNAseq analysis (see Table S2A and S2B) at 0 and 3hrs anti-IgM treatment. All values are the mean of 3 biological replicates. Adjusted p values for the differences gene expression levels were <0.05 for at least one of the points. **C, D**. ChIP-seq tracks (C) and RNAseq FPKM values (D) for active genes that are involved in autoimmunity that are downregulated in the Runx1 c-k/o resting B cells. **E.** Close up of RNA Pol II occupancy levels at poised genes that are regulated by RUNX1.

**Supplementary Table 1.**

Primers for gene expression analysis

| Target | Primer |
| --- | --- |
| *Ccnd2* | **Fw:** GAGTGGGAACTGGTAGTGTTG |
| | **Rv:** CGCACAGAGCGATGAAGGT |
| *Ccne1* | **Fw:** GTGGCTCCGACCTTTCAGTC |
| | **Rv:** CACAGTCTTGTCAATCTTGGCA |
| *E2f1* | **Fw:** CTCGACTCCTCGCAGATCG |
| | **Rv:** GATCCAGCCTCCGTTTCACC |
| *Fosl2* | **Fw:** CCAGCAGAAGTTCCGGGTAG |
| | **Rv:** GTAGGGATGTGAGCGTGGATA |
| *Atf3* | **Fw:** GAGGATTTTGCTAACCTGACACC |
| | **Rv:** TTGACGGTAACTGACTCCAGC |
| *Egr2* | **Fw:** CCTTTGACCAGATGAACGGAGTG |
| | **Rv:** CTGGTTTCTAGGTGCAGAGATGG |
| *c-Myc* | **Fw:** TCGCTGCTGTCCTCCGAGTCC |
| | **Rv:** GGTTTGCCTCTTCTCCACAGAC |
| *Rbpj* | **Fw:** CTCCACCCAAACGACTCACTA |
| | **Rv:** TCCAACCACTGCCCATAAGATA |
| *Notch2* | **Fw:** ATGTGGACGAGTGTCTGTTGC |
| | **Rv:** GGAAGCATAGGCACAGTCATC |
| *Psen2* | **Fw:** GAAGACTCCTACGACAGTTTTGG |
| | **Rv:** CACCAGGACGCTGTAGAAGAT |
| *Runx1* | **Fw**: AACGACCTCAGGTTTGTCGG |
| | **Rv:** TGGCAACTTGTGGCGGATTT |
| *Hprt* | **Fw:** GGGGGCTATAAGTTCTTTGC |
| | **Rv:** TCCAACACTTCGAGAGGTCC |
| Spike-in | **Fw:** GAGGGCCATACATGGTGGT |
| | **Rv:** CTCTAACCACTGGCGCCTTA |

Primers for ChIP

| Target | Primer |
| --- | --- |
| *Ccnd2 1* | **Fw:** AAAGGAAGACTGCTGGTCGG |
| | **Rv:** TTGAGGGTTCACAAAGGCGT |
| *Ccnd2 2* | **Fw:** CGTGTTGGAGGTAGGGATGG |
| | **Rv:** TACCTCCCGCAGTGTTCCTA |
| *Ccnd2 3* | **Fw:** TCGGATAGGGGAACCCACAA |
| | **Rv:** ATCACATTGCAAGCCTCCGA |
| *Ccnd2 4* | **Fw:** TCAACTTCAGGACACGCCTC |
| | **Rv:** GTAGGGATGTGAGCGTGGATA |
| *Ccnd2 5* | **Fw:** ATGCCCACGTATGTTTCCGT |
| | **Rv:** CTGGCAAATCAGCCCAAACC |

| | |
|---|---|
| *Ccnd2 6* | **Fw:** TCTGTGTGCATCTGCCTCTG |
| | **Rv:** TTCACAGCAGTCCTGACACC |
| *Ccnd2 pr* | **Fw:** TTGCAAGCCTCCGAAGTTAGA |
| | **Rv:** CCACAAACCCCATGGATTCCTA |
| *Rbpj pr* | **Fw:** GGGTTCTGACAGCTCTTGGG |
| | **Rv:** TGTAGCAAGCATCCCGTTCT |

Primers for genotyping

| Target | Primer | Product size (bp) | Tm |
|---|---|---|---|
| *Runx1* | P1 (Fw): CCCACTGTGTGCATTCCAGATTGG | WT=203<br>FL=275<br>Δ=310 | 58°C |
| | P2 (Rv): GACGGTGATGGTCAGAGTGAA GC | | |
| | P3 (Rv): CACCATAGCTTCTGGGTGCAG | | |
| *Cd23-Cre* | Fw: ATGCTCCTGTCTGTGTGCAG | 300 | 60°C |
| | Rv: GGTCAAAGTCAGTGCGTTCA | | |

Sequence of Spike-in RNA used in RT-qPCR expression analysis

| | |
|---|---|
| *Spike* | 5'CACCATGGTACTCTGGGGTGATATCTTTCCGGCATTTTGGGCAA<br>TGACCGGAACAGATAGGCATTGTGGGACCGGACGCGAGGGCCAT<br>ACATGGTGGTGCCGACATACCACCCGAGGCGCACCACGAGAATCA<br>TTTATTCTGTGTGTGGCATGTACGCTGACCATAAGGCGCCAGTGG<br>TTAGAGCTTTCCTTTCGGCATT3' |

**Table S2A. List of genes differentially expressed in resting B cells of Runx1 cK/O versus Cre control mice. P-values were generated by Wald test and p-adj are BH adjusted (DESeq2).**

| | Associated. Gene.Name | baseMean | log2Fold Change | lfcSE | stat | pvalue | padj | Cre_0 baseMean | Cre_3 baseMean | KO_0 baseMean | KO_3 baseMean |
|---|---|---|---|---|---|---|---|---|---|---|---|
| 1 | Dmwd | 740.7569 | 3.842332 | 0.17472 | -21.992 | 3.47E-107 | 5.26E-103 | 150.7769 | 26.31985 | 2207.437 | 578.494 |
| 2 | Plcxd2 | 483.2408 | -2.51836 | 0.181 | 13.9139 | 5.21E-44 | 3.94E-40 | 500.2545 | 891.4289 | 85.55695 | 455.723 |
| 3 | Ifnar1 | 4324.171 | 1.344699 | 0.09792 | -13.733 | 6.46E-43 | 3.26E-39 | 2694.542 | 2673.909 | 6863.977 | 5064.26 |
| 4 | Dab2ip | 274.2097 | 1.97465 | 0.14572 | -13.551 | 7.84E-42 | 2.97E-38 | 168.6841 | 37.46776 | 669.1077 | 221.579 |
| 5 | Lrig1 | 62.78883 | 2.872359 | 0.26216 | -10.957 | 6.18E-28 | 1.87E-24 | 21.97156 | 19.18423 | 167.0451 | 42.9544 |
| 6 | Rhobtb1 | 73.7577 | 2.284395 | 0.21523 | -10.614 | 2.57E-26 | 6.49E-23 | 38.45599 | 11.58919 | 191.984 | 53.0016 |
| 7 | Ski | 1225.733 | 0.663052 | 0.06479 | -10.235 | 1.39E-24 | 3.00E-21 | 1075.962 | 914.6709 | 1704.562 | 1207.74 |
| 8 | Cd200 | 782.1183 | 1.172515 | 0.11767 | -9.9648 | 2.17E-23 | 4.11E-20 | 596.1455 | 258.2163 | 1347.269 | 926.842 |
| 9 | Ass1 | 377.3964 | 1.16281 | 0.11865 | -9.8 | 1.13E-22 | 1.89E-19 | 199.4116 | 405.1855 | 448.8154 | 456.173 |
| 10 | Slc2a3 | 633.9378 | -0.90661 | 0.09446 | 9.59757 | 8.18E-22 | 1.24E-18 | 1098.55 | 474.3096 | 584.8259 | 378.066 |
| 11 | Eya1 | 91.24181 | -1.895 | 0.2077 | 9.12384 | 7.25E-20 | 9.98E-17 | 155.8559 | 134.2627 | 40.99364 | 33.8551 |
| 12 | Gm9855 | 84.55802 | 3.538251 | 0.3888 | -9.1004 | 9.00E-20 | 1.14E-16 | 7.69247 | 8.596798 | 102.4293 | 219.513 |
| 13 | Pisd-ps1 | 1992.179 | -0.84073 | 0.09357 | 8.98518 | 2.58E-19 | 3.01E-16 | 3654.134 | 1434.486 | 2036.677 | 843.418 |
| 14 | Slc2a6 | 506.8291 | -1.66463 | 0.18674 | 8.9142 | 4.91E-19 | 5.31E-16 | 182.4591 | 1326.727 | 56.21433 | 461.916 |
| 15 | St8sia6 | 268.5616 | -1.46265 | 0.17062 | 8.57242 | 1.01E-17 | 1.02E-14 | 423.4261 | 356.9067 | 152.131 | 141.782 |
| 16 | Gsn | 690.0347 | -1.03235 | 0.1289 | 8.00877 | 1.16E-15 | 1.10E-12 | 1604.784 | 249.1434 | 781.8312 | 124.38 |
| 17 | Saraf | 1143.446 | 0.929164 | 0.11645 | -7.9793 | 1.47E-15 | 1.31E-12 | 1119.065 | 401.3009 | 2137.335 | 916.084 |
| 18 | Rell1 | 201.6663 | -1.13143 | 0.14234 | 7.94869 | 1.88E-15 | 1.58E-12 | 420.5699 | 127.4846 | 191.2413 | 67.3696 |
| 19 | Il9r | 837.1027 | -2.19899 | 0.27708 | 7.93634 | 2.08E-15 | 1.65E-12 | 2074.102 | 684.8854 | 434.226 | 155.198 |
| 20 | Pdcd1lg2 | 209.8793 | -1.38012 | 0.17701 | 7.79707 | 6.34E-15 | 4.37E-12 | 170.4108 | 362.0827 | 64.35483 | 242.669 |
| 21 | Dusp3 | 708.0337 | 0.674319 | 0.08649 | -7.7967 | 6.36E-15 | 4.37E-12 | 352.2656 | 932.4256 | 562.2219 | 985.222 |
| 22 | Ccr6 | 974.1534 | -1.56728 | 0.20127 | 7.78691 | 6.87E-15 | 4.52E-12 | 1238.454 | 1796.21 | 411.5823 | 450.367 |
| 23 | Myo1f | 144.9685 | -1.53095 | 0.20091 | 7.61997 | 2.54E-14 | 1.60E-11 | 407.3018 | 18.25405 | 139.4086 | 14.9098 |
| 24 | Gm15920 | 73.18057 | 4.515319 | 0.59593 | -7.5769 | 3.54E-14 | 2.14E-11 | 1.595187 | 4.310023 | 72.87643 | 213.941 |
| 25 | Gm10036 | 68.59307 | 3.284261 | 0.4377 | -7.5034 | 6.22E-14 | 3.62E-11 | 9.856863 | 9.774205 | 114.6733 | 140.068 |
| 26 | Ehd3 | 167.0023 | -1.58249 | 0.21199 | 7.46485 | 8.34E-14 | 4.67E-11 | 438.6983 | 58.49175 | 144.4347 | 26.3845 |
| 27 | Scml4 | 473.7565 | -1.09789 | 0.14776 | 7.43036 | 1.08E-13 | 5.85E-11 | 1203.435 | 80.52926 | 559.7032 | 51.3588 |
| 28 | Sdc1 | 136.4381 | -1.33926 | 0.18243 | 7.34139 | 2.11E-13 | 1.10E-10 | 304.9092 | 64.43709 | 119.5387 | 56.8676 |
| 29 | Gpr171 | 1485.693 | -0.92317 | 0.12948 | 7.12976 | 1.01E-12 | 5.07E-10 | 1658.814 | 2330.591 | 871.8149 | 1081.55 |
| 30 | Pde7a | 2152.833 | -0.59777 | 0.08471 | 7.05685 | 1.70E-12 | 8.32E-10 | 3861.933 | 1251.829 | 2549.685 | 947.884 |
| 31 | Slco5a1 | 73.05885 | -2.06727 | 0.29578 | 6.98921 | 2.76E-12 | 1.30E-09 | 203.6684 | 36.26973 | 47.02055 | 5.27671 |
| 32 | Mb21d1 | 118.0569 | 1.492924 | 0.21371 | -6.9856 | 2.84E-12 | 1.30E-09 | 30.20043 | 97.58221 | 86.37955 | 258.066 |
| 33 | Rbpms | 113.7187 | 1.067539 | 0.1534 | -6.9593 | 3.42E-12 | 1.52E-09 | 114.9035 | 34.63995 | 242.033 | 63.2983 |
| 34 | Gm6969 | 57.58026 | -4.14821 | 0.59925 | 6.92236 | 4.44E-12 | 1.92E-09 | 168.4737 | 55.49357 | 6.353779 | 0 |
| 35 | Rps13-ps1 | 108.1333 | -4.9408 | 0.71757 | 6.88543 | 5.76E-12 | 2.42E-09 | 209.7466 | 218.4002 | 2.195302 | 2.19121 |
| 36 | Irf7 | 1046.817 | 1.012218 | 0.1475 | -6.8624 | 6.77E-12 | 2.77E-09 | 1017.043 | 244.6674 | 2061.222 | 864.333 |
| 37 | Hmgn2-ps1 | 88.90926 | 2.287094 | 0.33728 | -6.781 | 1.19E-11 | 4.75E-09 | 22.02156 | 24.63402 | 114.722 | 194.259 |
| 38 | Kctd14 | 290.8186 | 1.123582 | 0.16792 | -6.6911 | 2.22E-11 | 8.38E-09 | 155.4419 | 208.1909 | 340.5362 | 459.105 |
| 39 | 6330403A02 | 38.88587 | 2.12636 | 0.31869 | -6.6722 | 2.52E-11 | 9.30E-09 | 24.47554 | 4.836162 | 111.1581 | 15.0737 |
| 40 | Gm5806 | 50.29347 | 4.339636 | 0.65451 | -6.6303 | 3.35E-11 | 1.21E-08 | 1.331853 | 1.049371 | 59.00819 | 139.784 |
| 41 | Leprotl1 | 553.2531 | 0.997061 | 0.15054 | -6.6232 | 3.52E-11 | 1.24E-08 | 328.0553 | 381.5504 | 658.7951 | 844.612 |
| 42 | Amd2 | 49.31742 | 2.737904 | 0.41537 | -6.5915 | 4.35E-11 | 1.50E-08 | 5.30795 | 22.84561 | 40.16983 | 128.946 |
| 43 | Elk3 | 633.8462 | -0.55604 | 0.08566 | 6.49122 | 8.51E-11 | 2.86E-08 | 1092.017 | 292.7896 | 742.0572 | 408.521 |
| 44 | Slc26a11 | 299.5808 | -0.84809 | 0.13079 | 6.48414 | 8.92E-11 | 2.94E-08 | 454.4013 | 275.0386 | 251.1281 | 217.755 |
| 45 | Btla | 6695.415 | -0.62063 | 0.09781 | 6.34519 | 2.22E-10 | 7.15E-08 | 12383.78 | 3739.1 | 8043.244 | 2615.53 |
| 46 | Six5 | 11.96757 | 3.574513 | 0.56587 | -6.3169 | 2.67E-10 | 8.42E-08 | 1.575054 | 0.585938 | 26.50708 | 19.2022 |
| 47 | St6gal1 | 2972.185 | -0.87053 | 0.13923 | 6.25222 | 4.05E-10 | 1.25E-07 | 6042.535 | 1623.506 | 3292.458 | 930.239 |
| 48 | Rgl1 | 620.1177 | 0.781589 | 0.12532 | -6.2366 | 4.47E-10 | 1.35E-07 | 748.6018 | 187.4552 | 1290.335 | 254.078 |
| 49 | Dmpk | 214.5662 | 1.955021 | 0.31378 | -6.2306 | 4.65E-10 | 1.38E-07 | 147.0748 | 9.942798 | 594.5257 | 106.722 |
| 50 | Ints4 | 1576.786 | 0.482158 | 0.07786 | -6.1928 | 5.91E-10 | 1.72E-07 | 1587.2 | 1196.839 | 2218.305 | 1304.8 |
| 51 | Vps37b | 2171.97 | -0.60268 | 0.09791 | 6.15544 | 7.49E-10 | 2.14E-07 | 4427.443 | 727.0142 | 2911.539 | 621.885 |
| 52 | Abcb1b | 174.522 | -1.7899 | 0.29219 | 6.12574 | 9.03E-10 | 2.53E-07 | 81.3503 | 451.1266 | 22.45042 | 143.16 |
| 53 | Fer | 31.37153 | 2.00431 | 0.32822 | -6.1067 | 1.02E-09 | 2.80E-07 | 20.88903 | 3.272835 | 86.74451 | 14.5797 |
| 54 | Anxa6 | 2249.53 | -0.61453 | 0.10137 | 6.06255 | 1.34E-09 | 3.62E-07 | 4644.446 | 720.0288 | 3029.803 | 603.844 |
| 55 | Serpine2 | 52.75259 | 1.744203 | 0.2887 | -6.0416 | 1.53E-09 | 4.05E-07 | 33.56027 | 10.54314 | 115.6149 | 51.2921 |
| 56 | Pip5k1b | 111.7106 | -1.18145 | 0.19639 | 6.01585 | 1.79E-09 | 4.67E-07 | 211.7971 | 80.03192 | 92.73161 | 62.2819 |
| 57 | 5430427O19 | 768.8054 | -0.8656 | 0.144 | 6.01115 | 1.84E-09 | 4.73E-07 | 635.3995 | 1366.912 | 347.251 | 725.659 |
| 58 | Abcb1a | 41.14935 | -1.84134 | 0.30806 | 5.9773 | 2.27E-09 | 5.72E-07 | 71.1489 | 59.36171 | 19.09504 | 14.9917 |
| 59 | Antxr2 | 205.728 | -0.74521 | 0.12558 | 5.93392 | 2.96E-09 | 7.34E-07 | 435.1436 | 62.19882 | 259.1495 | 66.4202 |
| 60 | Galnt6 | 419.7679 | -0.56294 | 0.09624 | 5.84965 | 4.93E-09 | 1.20E-06 | 653.8039 | 292.321 | 441.7925 | 291.154 |
| 61 | Plekhg3 | 303.9683 | -0.8212 | 0.14063 | 5.83955 | 5.23E-09 | 1.26E-06 | 712.447 | 48.27405 | 402.0024 | 53.1497 |
| 62 | Nav2 | 44.65232 | 2.065093 | 0.3565 | -5.7926 | 6.93E-09 | 1.64E-06 | 26.45714 | 14.51888 | 117.4091 | 20.2242 |
| 63 | Fam69a | 611.8732 | -0.51571 | 0.08908 | 5.78904 | 7.08E-09 | 1.65E-06 | 797.8157 | 629.6696 | 557.5365 | 462.471 |
| 64 | Pvrl1 | 212.6365 | 1.137124 | 0.19865 | -5.7242 | 1.04E-08 | 2.38E-06 | 81.34554 | 245.483 | 181.5871 | 342.13 |
| 65 | Acap1 | 4005.878 | -0.58727 | 0.10268 | 5.71943 | 1.07E-08 | 2.40E-06 | 7354.849 | 1991.01 | 4887.824 | 1789.83 |
| 66 | B4galnt1 | 2398.634 | -0.47208 | 0.08256 | 5.71821 | 1.08E-08 | 2.40E-06 | 3613.036 | 1538.328 | 2601.984 | 1841.19 |
| 67 | Zap70 | 28.55081 | 1.695737 | 0.29758 | -5.6984 | 1.21E-08 | 2.64E-06 | 23.63688 | 2.083244 | 78.07331 | 10.4098 |
| 68 | Map4k3 | 299.9677 | 0.697746 | 0.12248 | -5.697 | 1.22E-08 | 2.64E-06 | 289.6948 | 163.5122 | 470.4141 | 276.25 |
| 69 | Ntng2 | 57.51625 | 1.217711 | 0.21401 | -5.69 | 1.27E-08 | 2.71E-06 | 55.55851 | 17.48913 | 130.6025 | 26.4149 |

| | | | | | | | | | | | |
|---|---|---|---|---|---|---|---|---|---|---|---|
| 70 | Emid1 | 309.3381 | -0.89026 | 0.15653 | 5.68745 | 1.29E-08 | 2.71E-06 | 752.5526 | 45.37495 | 404.1125 | 35.3125 |
| 71 | Gpr137b | 459.4167 | -0.62716 | 0.11201 | 5.59906 | 2.16E-08 | 4.47E-06 | 738.5556 | 398.3505 | 477.346 | 223.414 |
| 72 | Cnr2 | 2777.857 | 0.748761 | 0.13379 | -5.5966 | 2.19E-08 | 4.47E-06 | 3264.533 | 1180.727 | 5502.427 | 1163.74 |
| 73 | Fancc | 320.9376 | -0.7158 | 0.12813 | 5.58676 | 2.31E-08 | 4.67E-06 | 279.9925 | 522.5938 | 169.595 | 311.569 |
| 74 | Prss12 | 8.212996 | -3.46753 | 0.62148 | 5.57942 | 2.41E-08 | 4.81E-06 | 24.00609 | 2.554427 | 1.410042 | 4.88142 |
| 75 | Tmem26 | 308.646 | 0.624351 | 0.11236 | -5.5569 | 2.75E-08 | 5.40E-06 | 366.4001 | 87.25683 | 566.4185 | 214.509 |
| 76 | Plekhg5 | 40.9186 | 1.335154 | 0.2436 | -5.481 | 4.23E-08 | 8.21E-06 | 40.74596 | 6.263013 | 104.369 | 12.2965 |
| 77 | Kif11 | 178.6085 | -0.93346 | 0.17162 | 5.43925 | 5.35E-08 | 1.03E-05 | 255.5126 | 201.8629 | 133.073 | 123.985 |
| 78 | Pglyrp2 | 136.1108 | -1.00125 | 0.1851 | 5.40927 | 6.33E-08 | 1.20E-05 | 353.5334 | 7.920472 | 175.5693 | 7.4201 |
| 79 | Entpd5 | 763.7515 | -0.65199 | 0.12067 | 5.40285 | 6.56E-08 | 1.23E-05 | 1309.849 | 502.8766 | 831.8318 | 410.448 |
| 80 | Gm6548 | 181.3102 | 1.121409 | 0.20993 | -5.3418 | 9.20E-08 | 1.70E-05 | 138.1153 | 57.49982 | 303.8047 | 225.821 |
| 81 | Cep97 | 232.1293 | -0.8607 | 0.16355 | 5.26259 | 1.42E-07 | 2.53E-05 | 451.6416 | 134.5637 | 247.8283 | 94.4837 |
| 82 | Tgfa | 7.056969 | 3.251337 | 0.61938 | -5.2493 | 1.53E-07 | 2.69E-05 | 1.898118 | 0.696259 | 23.58518 | 2.04831 |
| 83 | 4930427A07 | 154.8037 | -0.82333 | 0.1575 | 5.22738 | 1.72E-07 | 2.99E-05 | 327.1404 | 58.99021 | 184.3854 | 48.6989 |
| 84 | Egr2 | 6710.806 | 1.253927 | 0.24244 | -5.1722 | 2.31E-07 | 3.98E-05 | 758.9964 | 10104.09 | 1843.241 | 14136.9 |
| 85 | Slc29a4 | 56.157 | 1.072139 | 0.20805 | -5.1532 | 2.56E-07 | 4.35E-05 | 51.14307 | 26.72279 | 108.1822 | 38.5799 |
| 86 | Rasgrf2 | 94.55426 | 0.930644 | 0.18126 | -5.1343 | 2.83E-07 | 4.76E-05 | 123.5815 | 6.493621 | 236.9828 | 11.1591 |
| 87 | Slc22a15 | 25.88676 | -2.20231 | 0.43177 | 5.10065 | 3.38E-07 | 5.57E-05 | 51.2123 | 31.81064 | 10.08524 | 10.4389 |
| 88 | Xxylt1 | 557.363 | -0.55804 | 0.10959 | 5.09198 | 3.54E-07 | 5.77E-05 | 502.7103 | 774.4297 | 341.9827 | 610.329 |
| 89 | Adk | 221.1175 | 0.684846 | 0.13497 | -5.074 | 3.90E-07 | 6.27E-05 | 186.5539 | 156.4787 | 301.5542 | 239.883 |
| 90 | Sspn | 185.8443 | 0.989118 | 0.19569 | -5.0545 | 4.31E-07 | 6.87E-05 | 158.7848 | 108.3003 | 318.3898 | 157.902 |
| 91 | Hook1 | 874.9783 | -0.58694 | 0.11756 | 4.99262 | 5.96E-07 | 9.39E-05 | 764.2485 | 1278.909 | 507.4641 | 949.292 |
| 92 | Rasa4 | 309.7848 | 0.738135 | 0.14791 | -4.9905 | 6.02E-07 | 9.40E-05 | 306.6126 | 158.9897 | 512.8601 | 260.677 |
| 93 | Sypl | 3618.109 | 0.425426 | 0.08528 | -4.9885 | 6.09E-07 | 9.40E-05 | 2301.044 | 4214.316 | 3091.512 | 4865.56 |
| 94 | Fam101b | 333.6741 | 0.77154 | 0.1553 | -4.968 | 6.77E-07 | 0.000103 | 468.3459 | 22.16445 | 802.2004 | 41.9855 |
| 95 | Bcar3 | 7625.999 | 0.339276 | 0.06854 | -4.9497 | 7.43E-07 | 0.000112 | 4130.141 | 10425.59 | 5226.83 | 10721.4 |
| 96 | Twsg1 | 4449.384 | 0.499658 | 0.10196 | -4.9004 | 9.56E-07 | 0.000143 | 1429.945 | 7480.191 | 2024.044 | 6863.36 |
| 97 | Hopx | 174.0787 | 0.893202 | 0.18231 | -4.8993 | 9.62E-07 | 0.000143 | 147.7686 | 69.36807 | 275.8305 | 203.348 |
| 98 | Epn2 | 90.97733 | 0.963556 | 0.1975 | -4.8788 | 1.07E-06 | 0.000157 | 75.94101 | 49.05292 | 149.1058 | 89.8096 |
| 99 | Abca8b | 16.50569 | 2.034715 | 0.41786 | -4.8694 | 1.12E-06 | 0.000163 | 12.29817 | 0 | 52.68083 | 1.04375 |
| 100 | Epb4.1 | 1774.983 | -0.59627 | 0.12289 | 4.85205 | 1.22E-06 | 0.000176 | 2794.386 | 1367.112 | 1844.348 | 1094.08 |
| 101 | Hal | 12.15324 | 2.771482 | 0.57211 | -4.8443 | 1.27E-06 | 0.000181 | 2.68745 | 4.189735 | 25.17581 | 16.56 |
| 102 | Gad1 | 278.101 | 0.848644 | 0.17538 | -4.8389 | 1.31E-06 | 0.000185 | 172.7883 | 269.9753 | 312.8804 | 356.76 |
| 103 | Stc1 | 13.99575 | 2.401376 | 0.50412 | -4.7635 | 1.90E-06 | 0.000267 | 7.652873 | 0.343146 | 44.84051 | 3.14645 |
| 104 | Ighv7-2 | 4.070748 | 3.934835 | 0.8306 | -4.7373 | 2.17E-06 | 0.000301 | 0 | 0 | 13.50934 | 2.77365 |
| 105 | Gm5815 | 5.319328 | -3.79985 | 0.80652 | 4.71143 | 2.46E-06 | 0.000338 | 13.60738 | 7.669931 | 0 | 0 |
| 106 | Bahd1 | 1031.349 | 0.323806 | 0.06918 | -4.681 | 2.86E-06 | 0.000389 | 1082.685 | 821.2061 | 1355.317 | 866.187 |
| 107 | Ift57 | 99.31614 | 1.04052 | 0.22248 | -4.677 | 2.91E-06 | 0.000393 | 45.03967 | 52.5604 | 94.29607 | 205.368 |
| 108 | Daam1 | 174.797 | -0.96842 | 0.20722 | 4.67338 | 2.96E-06 | 0.000397 | 98.02659 | 410.0408 | 49.52069 | 141.6 |
| 109 | Oas1a | 192.0071 | 0.735455 | 0.15792 | -4.6571 | 3.21E-06 | 0.000423 | 220.6267 | 42.94856 | 368.3039 | 136.149 |
| 110 | Gm6377 | 1739.559 | -0.63058 | 0.1358 | 4.64331 | 3.43E-06 | 0.000447 | 1773.054 | 2206.38 | 1142.268 | 1836.53 |
| 111 | Rab31 | 252.1037 | 0.757547 | 0.16323 | -4.6409 | 3.47E-06 | 0.000449 | 258.9376 | 130.5891 | 440.7093 | 178.179 |
| 112 | Nid1 | 277.4476 | 1.529485 | 0.32992 | -4.6359 | 3.55E-06 | 0.000453 | 257.2484 | 25.52767 | 771.0168 | 55.9976 |
| 113 | Pde2a | 1764.975 | 0.813485 | 0.1755 | -4.6353 | 3.56E-06 | 0.000453 | 2214.262 | 514.0531 | 3914.34 | 417.243 |
| 114 | Ndc80 | 168.7029 | -1.0287 | 0.2221 | 4.63173 | 3.63E-06 | 0.000457 | 368.8772 | 81.73256 | 178.9828 | 45.219 |
| 115 | Gm3608 | 72.71769 | -2.12007 | 0.4588 | 4.62092 | 3.82E-06 | 0.000478 | 117.8018 | 129.6811 | 24.0696 | 19.3183 |
| 116 | Clec16a | 864.2326 | 0.426644 | 0.09245 | -4.6149 | 3.93E-06 | 0.000484 | 619.0094 | 898.2662 | 832.8653 | 1106.79 |
| 117 | Slc9c1 | 4.202622 | 3.947565 | 0.85542 | -4.6148 | 3.94E-06 | 0.000484 | 0 | 0 | 15.72248 | 1.08801 |
| 118 | Stat4 | 930.6265 | -0.66049 | 0.14321 | 4.61203 | 3.99E-06 | 0.000487 | 871.219 | 1263.955 | 548.9277 | 1038.4 |
| 119 | Pear1 | 536.0533 | 0.605001 | 0.13146 | -4.6023 | 4.18E-06 | 0.000503 | 621.511 | 211.6469 | 946.4019 | 364.653 |
| 120 | 2810417H13 | 168.9216 | -0.72698 | 0.15797 | 4.60195 | 4.19E-06 | 0.000503 | 313.643 | 108.7704 | 188.5532 | 64.7199 |
| 121 | Lamp3 | 44.42644 | -1.16509 | 0.25359 | 4.59432 | 4.34E-06 | 0.000517 | 77.64978 | 39.11082 | 33.92625 | 27.0189 |
| 122 | Vps26b | 123.6154 | 0.755927 | 0.16463 | -4.5917 | 4.40E-06 | 0.00052 | 97.14448 | 76.74581 | 165.1382 | 155.433 |
| 123 | Rap1gap | 189.8781 | 1.102253 | 0.24092 | -4.5752 | 4.76E-06 | 0.000558 | 152.451 | 80.29713 | 331.6749 | 195.089 |
| 124 | Btnl1 | 462.1324 | 1.50556 | 0.32934 | -4.5714 | 4.84E-06 | 0.000564 | 33.70414 | 690.7572 | 100.6318 | 1023.44 |
| 125 | Itih5 | 131.8847 | -1.05628 | 0.23143 | 4.56414 | 5.02E-06 | 0.000579 | 346.3732 | 5.346113 | 164.8664 | 10.9532 |
| 126 | Nnt | 495.1172 | 0.508277 | 0.11156 | -4.5561 | 5.21E-06 | 0.000598 | 486.7926 | 350.8205 | 694.8474 | 448.008 |
| 127 | Ift81 | 20.69824 | 1.526011 | 0.33561 | -4.547 | 5.44E-06 | 0.000619 | 15.11842 | 9.397846 | 45.45589 | 12.8208 |
| 128 | Pdia4 | 3991.437 | 0.415135 | 0.09153 | -4.5355 | 5.75E-06 | 0.000649 | 3896.904 | 3195.009 | 5199.313 | 3674.52 |
| 129 | Cytip | 8203.16 | -0.4175 | 0.09282 | 4.49774 | 6.87E-06 | 0.00077 | 14811.92 | 3520.703 | 11080.97 | 3399.04 |
| 130 | Fyn | 4308.202 | 0.775136 | 0.17293 | -4.4825 | 7.38E-06 | 0.000821 | 1094.074 | 5918.358 | 1882.606 | 8337.77 |
| 131 | Il18 | 126.6362 | 0.886803 | 0.19823 | -4.4737 | 7.69E-06 | 0.000849 | 168.9988 | 4.738025 | 314.2087 | 18.5995 |
| 132 | Gnaq | 997.6091 | 0.424998 | 0.09534 | -4.4579 | 8.28E-06 | 0.000907 | 624.6192 | 1156.017 | 840.1699 | 1369.63 |
| 133 | Rasgef1b | 11997.18 | -0.3302 | 0.07409 | 4.45662 | 8.33E-06 | 0.000907 | 12896.55 | 13378.36 | 10253.11 | 11460.7 |
| 134 | Top2a | 763.1014 | -0.57331 | 0.12883 | 4.45028 | 8.58E-06 | 0.000927 | 1002.767 | 815.3482 | 672.982 | 561.308 |
| 135 | Qrfp | 19.16935 | -1.64791 | 0.37202 | 4.42967 | 9.44E-06 | 0.001013 | 49.06283 | 5.594436 | 15.04616 | 6.97399 |
| 136 | Sec16b | 39.35039 | 1.135652 | 0.25671 | -4.424 | 9.69E-06 | 0.001033 | 46.3342 | 3.493477 | 103.3445 | 4.2294 |
| 137 | Whrn | 770.5132 | 0.541278 | 0.12241 | -4.4218 | 9.79E-06 | 0.001036 | 851.1784 | 381.2769 | 1239.761 | 609.837 |
| 138 | Usp18 | 193.0904 | 0.732708 | 0.16582 | -4.4188 | 9.92E-06 | 0.001043 | 125.6339 | 91.5483 | 210.1824 | 344.997 |
| 139 | Ndrg1 | 85.75889 | -0.787 | 0.17817 | 4.41704 | 1.00E-05 | 0.001044 | 151.8223 | 48.084 | 87.40053 | 55.7287 |
| 140 | Folr4 | 24.04767 | 1.553856 | 0.35271 | -4.4055 | 1.06E-05 | 0.001094 | 19.63998 | 3.002362 | 59.81507 | 13.7333 |
| 141 | Capn2 | 364.0678 | 0.657008 | 0.1494 | -4.3978 | 1.09E-05 | 0.001124 | 464.252 | 114.4028 | 734.671 | 142.946 |
| 142 | Hcrtr2 | 69.79758 | 0.985039 | 0.22409 | -4.3957 | 1.10E-05 | 0.001124 | 84.66441 | 11.71723 | 168.881 | 13.9277 |

| | | | | | | | | | | | |
|---|---|---|---|---|---|---|---|---|---|---|---|
| 143 | Rhbdf1 | 131.3186 | -1.1233 | 0.25557 | 4.3953 | 1.11E-05 | 0.001124 | 265.824 | 84.00307 | 119.9297 | 55.5178 |
| 144 | Tgfb3 | 161.194 | -0.77505 | 0.17648 | 4.39159 | 1.13E-05 | 0.001135 | 180.451 | 225.1897 | 104.5379 | 134.597 |
| 145 | Sdc4 | 1926.589 | -0.49989 | 0.11398 | 4.38597 | 1.15E-05 | 0.001157 | 2864.187 | 1673.018 | 2022.323 | 1146.83 |
| 146 | 5330426P16 | 59.80492 | -0.87812 | 0.20053 | 4.37893 | 1.19E-05 | 0.001188 | 103.758 | 50.46275 | 56.18183 | 28.8171 |
| 147 | Gm14292 | 6.351251 | -3.41276 | 0.77993 | 4.37574 | 1.21E-05 | 0.001197 | 10.6498 | 13.38797 | 0 | 1.36723 |
| 148 | Hepacam2 | 150.3691 | 0.731031 | 0.16734 | -4.3685 | 1.25E-05 | 0.001229 | 216.8338 | 5.586687 | 361.1238 | 17.9323 |
| 149 | Kctd10 | 712.171 | -0.41988 | 0.09622 | 4.36368 | 1.28E-05 | 0.001249 | 998.136 | 581.3843 | 745.2272 | 523.937 |
| 150 | Ube2e2 | 6.10933 | -2.78156 | 0.63775 | 4.36155 | 1.29E-05 | 0.001253 | 19.18118 | 2.424173 | 2.150888 | 0.68108 |
| 151 | Mprip | 2102.274 | -0.30645 | 0.07038 | 4.3541 | 1.34E-05 | 0.001288 | 2752.915 | 1842.586 | 2225.006 | 1588.59 |
| 152 | Il10rb | 1151.206 | 0.369342 | 0.08539 | -4.3255 | 1.52E-05 | 0.001458 | 756.3454 | 1311.169 | 977.3763 | 1559.93 |
| 153 | Hspbap1 | 201.5165 | -0.60541 | 0.14022 | 4.31774 | 1.58E-05 | 0.001498 | 204.3778 | 252.9393 | 133.7808 | 214.968 |
| 154 | Pik3r6 | 11.01847 | 2.018168 | 0.46752 | -4.3167 | 1.58E-05 | 0.001498 | 6.91648 | 2.090993 | 30.39968 | 4.66671 |
| 155 | Rgs13 | 69.21763 | -1.19281 | 0.2775 | 4.29846 | 1.72E-05 | 0.001617 | 88.81836 | 102.6339 | 37.88504 | 47.5332 |
| 156 | Slc4a11 | 67.63383 | -0.9698 | 0.22682 | 4.27572 | 1.91E-05 | 0.001765 | 89.58101 | 74.14553 | 44.88204 | 61.9267 |
| 157 | Yes1 | 79.55911 | 1.110151 | 0.25969 | -4.2749 | 1.91E-05 | 0.001765 | 18.41492 | 110.2865 | 40.21131 | 149.324 |
| 158 | Bspry | 289.1905 | 0.988532 | 0.23126 | -4.2745 | 1.92E-05 | 0.001765 | 70.74028 | 367.2504 | 142.7751 | 575.996 |
| 159 | Rgs2 | 5999.457 | -0.61687 | 0.14435 | 4.27356 | 1.92E-05 | 0.001765 | 12966.69 | 1426.998 | 8431.259 | 1172.88 |
| 160 | Manea | 1558.941 | -0.51615 | 0.12083 | 4.27158 | 1.94E-05 | 0.00177 | 2243.286 | 1600.333 | 1565.87 | 826.276 |
| 161 | Ipcef1 | 185.2735 | -1.03742 | 0.24299 | 4.26938 | 1.96E-05 | 0.001777 | 470.7409 | 26.77112 | 226.5671 | 17.0147 |
| 162 | Eps8l1 | 315.2604 | -1.30705 | 0.30724 | 4.2542 | 2.10E-05 | 0.00189 | 721.729 | 185.9975 | 283.1771 | 70.1379 |
| 163 | Sh3bp5 | 4944.66 | 0.28807 | 0.06777 | -4.2505 | 2.13E-05 | 0.00191 | 5604.592 | 3454.752 | 6845.262 | 3874.03 |
| 164 | Fam167a | 3256.628 | 0.388273 | 0.09158 | -4.2395 | 2.24E-05 | 0.00199 | 1530.906 | 4994.245 | 2004.796 | 4496.57 |
| 165 | Tnik | 179.4568 | -0.62788 | 0.14813 | 4.23865 | 2.25E-05 | 0.00199 | 220.9201 | 213.1267 | 142.5023 | 141.278 |
| 166 | Cdh24 | 107.4303 | -0.81839 | 0.19345 | 4.23052 | 2.33E-05 | 0.002052 | 92.22568 | 193.7134 | 51.46734 | 92.3147 |
| 167 | Ncf2 | 2449.462 | 0.334499 | 0.07923 | -4.222 | 2.42E-05 | 0.002118 | 3218.22 | 1044.528 | 4059.425 | 1475.68 |
| 168 | Tmem154 | 537.5289 | 0.674832 | 0.15989 | -4.2207 | 2.44E-05 | 0.002119 | 642.5037 | 172.6757 | 1029.241 | 305.695 |
| 169 | Gm11346 | 134.7267 | -0.82684 | 0.19602 | 4.21825 | 2.46E-05 | 0.002129 | 331.7204 | 11.03315 | 185.8115 | 10.3417 |
| 170 | Ncf1 | 3485.571 | -0.42894 | 0.10179 | 4.2141 | 2.51E-05 | 0.002149 | 3973.166 | 4001.574 | 2948.412 | 3019.13 |
| 171 | Clic4 | 6423.972 | 0.378701 | 0.08988 | -4.2136 | 2.51E-05 | 0.002149 | 2990.322 | 9094.155 | 3890.912 | 9720.5 |
| 172 | 9430038I01F | 323.624 | 0.527098 | 0.12583 | -4.1889 | 2.80E-05 | 0.002383 | 249.969 | 303.4884 | 360.6699 | 380.369 |
| 173 | Slc35d2 | 143.771 | 0.6016 | 0.14429 | -4.1693 | 3.06E-05 | 0.002583 | 181.7085 | 45.9159 | 276.1108 | 71.3487 |
| 174 | Rpl30-ps9 | 4.6564 | -3.16184 | 0.76264 | 4.14592 | 3.38E-05 | 0.002824 | 9.897129 | 8.393872 | 0.334598 | 0 |
| 175 | Ifi27l2a | 960.5956 | 0.889646 | 0.21462 | -4.1451 | 3.40E-05 | 0.002824 | 1060.678 | 225.777 | 1984.263 | 571.665 |
| 176 | Ryr2 | 58.08955 | -1.28186 | 0.30972 | 4.13876 | 3.49E-05 | 0.002888 | 122.6313 | 43.17404 | 48.96301 | 17.5899 |
| 177 | Spag5 | 89.44948 | -0.70721 | 0.17199 | 4.11205 | 3.92E-05 | 0.003226 | 146.3363 | 71.27276 | 89.55914 | 50.6298 |
| 178 | Tor3a | 1148.055 | 0.380814 | 0.09267 | -4.1092 | 3.97E-05 | 0.003248 | 823.4909 | 1057.784 | 1072.522 | 1638.42 |
| 179 | Hist1h2al | 3.252417 | -3.35755 | 0.81809 | 4.10411 | 4.06E-05 | 0.003303 | 8.261675 | 4.747991 | 0 | 0 |
| 180 | mt-Tn | 471.6366 | 0.549765 | 0.13425 | -4.0952 | 4.22E-05 | 0.003415 | 610.5796 | 180.3094 | 896.249 | 199.408 |
| 181 | Gm8093 | 5.357401 | 3.061455 | 0.7487 | -4.089 | 4.33E-05 | 0.003487 | 1.25199 | 0.343146 | 17.83547 | 1.99899 |
| 182 | Galc | 102.8294 | 0.803089 | 0.19716 | -4.0733 | 4.64E-05 | 0.003712 | 68.75826 | 84.99057 | 120.5746 | 136.994 |
| 183 | Orai2 | 862.8805 | -0.44435 | 0.10922 | 4.06844 | 4.73E-05 | 0.003737 | 1621.225 | 320.9334 | 1190.526 | 318.837 |
| 184 | AI504432 | 506.9052 | -1.08981 | 0.26788 | 4.0683 | 4.74E-05 | 0.003737 | 898.1899 | 463.5367 | 414.235 | 251.659 |
| 185 | Ddx25 | 63.56512 | 0.770001 | 0.18928 | -4.0681 | 4.74E-05 | 0.003737 | 88.22333 | 3.242936 | 151.1295 | 11.6647 |
| 186 | Nlrc3 | 211.6837 | -1.05881 | 0.26198 | 4.04159 | 5.31E-05 | 0.004142 | 451.9824 | 106.5908 | 213.309 | 74.8527 |
| 187 | Scpep1 | 1358.91 | -0.33225 | 0.08234 | 4.03521 | 5.46E-05 | 0.004234 | 1662.936 | 1371.52 | 1320.545 | 1080.64 |
| 188 | Nedd4 | 2092.746 | 0.760479 | 0.18875 | -4.029 | 5.60E-05 | 0.004325 | 2667.158 | 218.7511 | 4547.004 | 938.071 |
| 189 | Zfp365 | 38.54705 | 1.034786 | 0.25707 | -4.0254 | 5.69E-05 | 0.004371 | 32.47455 | 18.34997 | 67.55519 | 35.8085 |
| 190 | Abhd10 | 240.2811 | -0.51746 | 0.12861 | 4.02362 | 5.73E-05 | 0.004381 | 247.0955 | 242.7681 | 172.4844 | 298.776 |
| 191 | Bmi1 | 1892.532 | 0.315686 | 0.07879 | -4.0066 | 6.16E-05 | 0.004686 | 1323.728 | 2225.031 | 1647.425 | 2373.94 |
| 192 | Smoc1 | 18.79261 | -1.71223 | 0.42818 | 3.99881 | 6.37E-05 | 0.004818 | 28.18152 | 24.54654 | 7.87276 | 14.5696 |
| 193 | Fam149b | 552.5638 | -0.44091 | 0.11062 | 3.98584 | 6.72E-05 | 0.005063 | 1026.336 | 219.6219 | 755.157 | 209.14 |
| 194 | E330020D12 | 901.5681 | 0.721698 | 0.18112 | -3.9845 | 6.76E-05 | 0.005066 | 508.1889 | 1154.427 | 843.3738 | 1100.28 |
| 195 | Cers6 | 120.5132 | -0.65559 | 0.16459 | 3.98322 | 6.80E-05 | 0.005069 | 160.9115 | 135.9773 | 101.8569 | 83.307 |
| 196 | Iqsec2 | 135.566 | 0.905491 | 0.2282 | -3.9679 | 7.25E-05 | 0.005379 | 50.64694 | 137.4923 | 95.89029 | 258.235 |
| 197 | Aspm | 98.74662 | -0.69416 | 0.17509 | 3.96456 | 7.35E-05 | 0.005429 | 159.2269 | 83.38697 | 97.94714 | 54.4254 |
| 198 | Dtd2 | 80.44714 | 0.941315 | 0.23754 | -3.9628 | 7.41E-05 | 0.005442 | 39.51183 | 87.42874 | 77.06946 | 117.779 |
| 199 | Ltk | 221.1665 | 0.832054 | 0.21112 | -3.9411 | 8.11E-05 | 0.005931 | 286.9564 | 31.25901 | 514.3394 | 52.111 |
| 200 | Ighg2b | 102.6915 | -2.05453 | 0.52151 | 3.93958 | 8.16E-05 | 0.005939 | 230.046 | 106.1186 | 47.69955 | 26.9017 |
| 201 | Ncapg | 68.17973 | -0.82306 | 0.20919 | 3.93457 | 8.33E-05 | 0.006036 | 101.843 | 69.07697 | 56.97907 | 44.8199 |
| 202 | Ggnbp1 | 108.3168 | 0.704124 | 0.17938 | -3.9253 | 8.66E-05 | 0.006243 | 118.1609 | 49.4898 | 192.9905 | 72.6259 |
| 203 | Gpr146 | 94.14612 | -0.77884 | 0.19855 | 3.92264 | 8.76E-05 | 0.006282 | 146.864 | 85.90749 | 85.48237 | 58.3306 |
| 204 | Gm11223 | 7.104997 | 2.871884 | 0.73693 | -3.8971 | 9.74E-05 | 0.006918 | 1.19226 | 0.585938 | 16.74992 | 9.89187 |
| 205 | Dennd4a | 6281.637 | -0.60116 | 0.15434 | 3.8951 | 9.82E-05 | 0.006942 | 5751.694 | 10321.8 | 3778.861 | 5274.2 |
| 206 | Rpl30-ps10 | 3.001663 | -3.19624 | 0.82142 | 3.89113 | 9.98E-05 | 0.007024 | 7.51622 | 4.17202 | 0 | 0.31841 |
| 207 | Limk2 | 1182.277 | -0.26831 | 0.06902 | 3.88727 | 0.000101 | 0.007104 | 1434.244 | 1096.881 | 1191.185 | 1006.8 |
| 208 | Cep57l1 | 115.5097 | -0.63732 | 0.16427 | 3.87973 | 0.000105 | 0.007289 | 202.3326 | 75.45875 | 129.9439 | 54.3036 |
| 209 | Pkd1l1 | 6.62536 | 3.177537 | 0.81921 | -3.8788 | 0.000105 | 0.007289 | 0.949059 | 0 | 21.90693 | 3.64545 |
| 210 | Sh3bp2 | 1927.563 | -0.2247 | 0.05795 | 3.87731 | 0.000106 | 0.007299 | 1931.23 | 1983 | 1652.497 | 2143.53 |
| 211 | A930033H14 | 8.00832 | 2.191278 | 0.56685 | -3.8657 | 0.000111 | 0.00762 | 2.797179 | 3.493477 | 15.85865 | 9.88397 |
| 212 | Foxm1 | 245.8735 | -0.54021 | 0.14005 | 3.85721 | 0.000115 | 0.007834 | 277.559 | 281.0063 | 190.6823 | 234.246 |
| 213 | Pisd-ps2 | 286.5423 | -0.77906 | 0.202 | 3.85675 | 0.000115 | 0.007834 | 515.6233 | 211.6079 | 298.2411 | 120.697 |
| 214 | Plekha6 | 16.27013 | -1.47601 | 0.38389 | 3.84486 | 0.000121 | 0.008185 | 42.84407 | 6.062304 | 14.80691 | 1.36723 |
| 215 | Ccdc88b | 555.2821 | -0.35976 | 0.09359 | 3.84382 | 0.000121 | 0.008185 | 936.2453 | 283.2012 | 729.3637 | 272.318 |

| | | | | | | | | | | | |
|---|---|---|---|---|---|---|---|---|---|---|---|
| 216 | Krt222 | 100.6473 | 0.81694 | 0.21263 | -3.8421 | 0.000122 | 0.008205 | 117.6071 | 32.86883 | 208.2614 | 43.8517 |
| 217 | A630023P12 | 50.85308 | 1.099418 | 0.28632 | -3.8398 | 0.000123 | 0.008247 | 60.21758 | 2.911974 | 131.8189 | 8.46386 |
| 218 | Mical1 | 1424.822 | -0.357 | 0.093 | 3.83869 | 0.000124 | 0.008247 | 2296.964 | 954.1186 | 1791.277 | 656.927 |
| 219 | Cish | 1003.913 | 2.432913 | 0.63646 | -3.8225 | 0.000132 | 0.008768 | 14.95576 | 1831.471 | 131.4269 | 2037.8 |
| 220 | Igkv4-57-1 | 127.9356 | 0.640309 | 0.1676 | -3.8204 | 0.000133 | 0.008807 | 164.1913 | 39.67904 | 256.6519 | 51.2203 |
| 221 | Dirc2 | 125.643 | -0.5645 | 0.14787 | 3.81749 | 0.000135 | 0.008872 | 197.5361 | 94.02271 | 133.3687 | 77.6444 |
| 222 | Uhrf1 | 441.6401 | -0.46549 | 0.12227 | 3.80705 | 0.000141 | 0.009175 | 482.289 | 531.1431 | 349.3072 | 403.821 |
| 223 | Lgals3bp | 803.5846 | 0.558085 | 0.14708 | -3.7944 | 0.000148 | 0.009615 | 863.7509 | 300.3335 | 1275.213 | 775.041 |
| 224 | Lrrc32 | 1233.594 | 2.151193 | 0.56757 | -3.7902 | 0.000151 | 0.009734 | 38.95867 | 1453.513 | 225.2531 | 3216.65 |
| 225 | Ighg2c | 66.12465 | -2.89802 | 0.76498 | 3.78837 | 0.000152 | 0.009734 | 167.0284 | 79.78988 | 11.42657 | 6.25372 |
| 226 | Mybl1 | 36.11303 | -1.31778 | 0.34948 | 3.77072 | 0.000163 | 0.010351 | 62.26257 | 43.74806 | 24.03878 | 14.4027 |
| 227 | Casc5 | 120.8898 | -0.95063 | 0.25211 | 3.77072 | 0.000163 | 0.010351 | 213.9769 | 109.8242 | 109.1921 | 50.566 |
| 228 | Dennd3 | 1243.183 | -0.48995 | 0.13013 | 3.7651 | 0.000166 | 0.010534 | 2716.205 | 142.3129 | 1930.314 | 183.899 |
| 229 | Adm | 475.8729 | 0.925442 | 0.24585 | -3.7643 | 0.000167 | 0.010534 | 97.76705 | 741.6553 | 189.3369 | 874.732 |
| 230 | Cacna2d2 | 13.16521 | -1.53688 | 0.41014 | 3.74722 | 0.000179 | 0.011229 | 35.63758 | 2.444106 | 11.75111 | 2.82804 |
| 231 | Skap1 | 19.22004 | -1.30861 | 0.34961 | 3.74306 | 0.000182 | 0.011369 | 44.40688 | 8.361755 | 17.50022 | 6.61132 |
| 232 | Ado | 1058.126 | 0.6851 | 0.18332 | -3.7371 | 0.000186 | 0.011593 | 315.5058 | 1512.822 | 510.6977 | 1893.48 |
| 233 | Dna2 | 70.98579 | -0.73782 | 0.19764 | 3.7332 | 0.000189 | 0.011727 | 118.214 | 57.01064 | 70.56425 | 38.1542 |
| 234 | Epha2 | 284.006 | -0.64958 | 0.17442 | 3.72428 | 0.000196 | 0.0121 | 439.5629 | 200.0845 | 279.2655 | 217.111 |
| 235 | Dclre1a | 91.2442 | 0.670757 | 0.18029 | -3.7205 | 0.000199 | 0.012235 | 71.42852 | 58.64595 | 113.9828 | 120.919 |
| 236 | Gpd1l | 2543.193 | -0.42224 | 0.1142 | 3.69755 | 0.000218 | 0.013339 | 4693.481 | 948.3841 | 3499.099 | 1031.81 |
| 237 | Ahnak2 | 12.03813 | 2.288064 | 0.61945 | -3.6937 | 0.000221 | 0.013488 | 1.301988 | 11.93565 | 9.996413 | 24.9185 |
| 238 | Ckap4 | 460.0014 | 0.366208 | 0.09922 | -3.6907 | 0.000224 | 0.013593 | 588.2347 | 198.1105 | 757.4922 | 296.168 |
| 239 | Capg | 5337.819 | 0.724717 | 0.19644 | -3.6892 | 0.000225 | 0.013617 | 3323.026 | 4227.541 | 5527.578 | 8273.13 |
| 240 | Rsu1 | 436.1412 | 0.445329 | 0.12121 | -3.6741 | 0.000239 | 0.014373 | 650.0823 | 93.37519 | 886.2304 | 114.877 |
| 241 | Klrb1f | 10.46633 | -1.79946 | 0.48988 | 3.67325 | 0.000239 | 0.014373 | 31.62231 | 0.931301 | 8.586371 | 0.72534 |
| 242 | Rundc3b | 75.94964 | -1.01635 | 0.27675 | 3.67242 | 0.00024 | 0.014373 | 34.26294 | 172.3219 | 16.45913 | 80.7546 |
| 243 | Lztfl1 | 1235.979 | -0.39982 | 0.10903 | 3.66725 | 0.000245 | 0.014609 | 910.7191 | 2115.959 | 689.5991 | 1227.64 |
| 244 | Nr2f6 | 78.4889 | 0.760764 | 0.20776 | -3.6617 | 0.000251 | 0.014872 | 52.09892 | 72.59992 | 88.75605 | 100.501 |
| 245 | Dsn1 | 169.4503 | -0.52147 | 0.14291 | 3.649 | 0.000263 | 0.015564 | 217.4207 | 185.312 | 150.9032 | 124.165 |
| 246 | Rtp4 | 199.8168 | 0.612771 | 0.16871 | -3.6322 | 0.000281 | 0.016549 | 182.7218 | 78.06439 | 280.1963 | 258.285 |
| 247 | Nrip1 | 913.3527 | -0.53288 | 0.14698 | 3.62564 | 0.000288 | 0.016909 | 898.5021 | 1308.135 | 619.388 | 827.386 |
| 248 | Mki67 | 840.0968 | -0.79453 | 0.21931 | 3.62284 | 0.000291 | 0.017028 | 1429.355 | 742.1782 | 817.0144 | 371.839 |
| 249 | Sema4d | 3104.353 | -0.53012 | 0.14644 | 3.61994 | 0.000295 | 0.017153 | 5528.43 | 1870.976 | 3818.352 | 1199.65 |
| 250 | Cr2 | 28210.46 | 0.476731 | 0.13176 | -3.6182 | 0.000297 | 0.017203 | 16096.13 | 32723.89 | 22440.79 | 41581 |
| 251 | Ttc28 | 601.9041 | 0.934961 | 0.25889 | -3.6115 | 0.000304 | 0.017588 | 499.6586 | 332.478 | 969.3513 | 606.128 |
| 252 | Cdc6 | 43.48643 | -0.91923 | 0.25467 | 3.6095 | 0.000307 | 0.017655 | 76.38849 | 29.56875 | 39.77783 | 28.2106 |
| 253 | Scimp | 64.39342 | -0.90677 | 0.25132 | 3.60803 | 0.000309 | 0.017688 | 155.9463 | 11.74713 | 82.41591 | 7.46436 |
| 254 | Tacc2 | 34.77998 | -1.5206 | 0.42177 | 3.60525 | 0.000312 | 0.017767 | 18.23188 | 58.24191 | 5.711817 | 56.9343 |
| 255 | Pim2 | 504.6694 | -0.41222 | 0.11435 | 3.60492 | 0.000312 | 0.017767 | 642.8488 | 446.2278 | 482.4448 | 447.156 |
| 256 | Amigo2 | 529.4001 | 0.479539 | 0.13321 | -3.5999 | 0.000318 | 0.018046 | 295.8625 | 452.6381 | 413.6759 | 955.424 |
| 257 | Pxk | 6180.69 | -0.21958 | 0.06113 | 3.59182 | 0.000328 | 0.018544 | 7676.398 | 5914.816 | 6590.967 | 4540.58 |
| 258 | Ptk2b | 6394.189 | -0.37489 | 0.10441 | 3.59048 | 0.00033 | 0.018571 | 10590.65 | 3751.453 | 8159.131 | 3075.52 |
| 259 | Ticrr | 106.6915 | -0.75157 | 0.20979 | 3.58258 | 0.00034 | 0.019071 | 133.1361 | 126.6503 | 78.72947 | 88.2501 |
| 260 | Sbf1 | 3478.227 | -0.35582 | 0.09985 | 3.56359 | 0.000366 | 0.02043 | 3664.086 | 3903.695 | 2860.325 | 3484.8 |
| 261 | Polm | 1122.178 | 0.451802 | 0.12695 | -3.5588 | 0.000373 | 0.020652 | 1417.555 | 495.9448 | 1942.706 | 632.507 |
| 262 | Echdc3 | 21.06421 | 1.207005 | 0.33958 | -3.5544 | 0.000379 | 0.020928 | 18.30158 | 10.48265 | 42.92599 | 12.5466 |
| 263 | Rogdi | 683.4643 | -0.4499 | 0.12682 | 3.54747 | 0.000389 | 0.021407 | 981.7262 | 513.7748 | 717.6203 | 520.736 |
| 264 | Sertad4 | 21.02831 | -1.57587 | 0.4449 | 3.54203 | 0.000397 | 0.021773 | 21.4053 | 34.71374 | 6.445539 | 21.5487 |
| 265 | Dcaf12 | 1051.198 | -0.2486 | 0.07024 | 3.53922 | 0.000401 | 0.02185 | 1451.589 | 731.148 | 1221.03 | 801.025 |
| 266 | Apbb1ip | 1524.906 | -0.23 | 0.06499 | 3.5392 | 0.000401 | 0.02185 | 2198.945 | 919.9587 | 1873.271 | 1107.45 |
| 267 | Ryk | 31.68886 | 1.187625 | 0.33696 | -3.5245 | 0.000424 | 0.023018 | 15.88468 | 22.25483 | 37.61204 | 51.0039 |
| 268 | Klf13 | 2581 | -0.36446 | 0.10344 | 3.52344 | 0.000426 | 0.023026 | 4635.683 | 1049.166 | 3598.199 | 1040.95 |
| 269 | Arap1 | 4039.551 | 0.274248 | 0.07795 | -3.5184 | 0.000434 | 0.023266 | 4916.294 | 2465.661 | 5947.518 | 2828.73 |
| 270 | Ctla4 | 75.52809 | -0.61576 | 0.17501 | 3.51836 | 0.000434 | 0.023266 | 151.623 | 24.74546 | 98.57839 | 27.1656 |
| 271 | Prc1 | 154.9353 | -0.54487 | 0.15489 | 3.51786 | 0.000435 | 0.023266 | 169.6349 | 184.4724 | 116.1917 | 149.442 |
| 272 | Gm15228 | 12.3667 | 2.087245 | 0.59356 | -3.5165 | 0.000437 | 0.023266 | 6.959687 | 0 | 35.26757 | 7.23952 |
| 273 | Cldn10 | 34.55796 | -0.90709 | 0.25799 | 3.51599 | 0.000438 | 0.023266 | 57.69622 | 22.50316 | 30.33096 | 27.7015 |
| 274 | Ptger4 | 1082.648 | -0.4171 | 0.11888 | 3.50856 | 0.000451 | 0.023842 | 1606.653 | 797.1485 | 1201.942 | 724.85 |
| 275 | Bach1 | 1284.219 | -0.32468 | 0.09285 | 3.49688 | 0.000471 | 0.024824 | 1283.01 | 1469.5 | 1023.644 | 1360.72 |
| 276 | Stk38 | 1627.553 | -0.30151 | 0.08653 | 3.48436 | 0.000493 | 0.025925 | 2345.42 | 1159.864 | 1901.706 | 1103.22 |
| 277 | Stil | 42.29777 | -0.92319 | 0.26528 | 3.48013 | 0.000501 | 0.026247 | 71.28096 | 38.09134 | 37.28582 | 22.533 |
| 278 | Kremen1 | 485.2043 | -0.32911 | 0.09476 | 3.47298 | 0.000515 | 0.026862 | 533.6048 | 557.3306 | 424.457 | 425.425 |
| 279 | Pdlim1 | 563.9337 | 0.552684 | 0.15919 | -3.4719 | 0.000517 | 0.026875 | 389.7098 | 542.9627 | 573.9618 | 749.101 |
| 280 | Gpr155 | 305.8732 | -0.52091 | 0.15011 | 3.47013 | 0.00052 | 0.026963 | 547.6128 | 167.3027 | 380.7816 | 127.796 |
| 281 | Smc2 | 768.78 | -0.42835 | 0.12354 | 3.46738 | 0.000526 | 0.027147 | 636.0366 | 1078.321 | 472.4729 | 888.289 |
| 282 | Ifnar2 | 927.5591 | 0.411328 | 0.11888 | -3.4601 | 0.00054 | 0.027795 | 532.7237 | 1124.936 | 709.0308 | 1343.55 |
| 283 | Pkib | 6966.541 | 0.905211 | 0.26172 | -3.4587 | 0.000543 | 0.027849 | 877.4985 | 11271.67 | 1668.877 | 14048.1 |
| 284 | Cenpe | 128.3162 | -0.70808 | 0.20482 | 3.45713 | 0.000546 | 0.027916 | 213.6954 | 113.5339 | 130.1578 | 55.8776 |
| 285 | Npr2 | 27.41786 | 1.415527 | 0.40959 | -3.4559 | 0.000548 | 0.027945 | 17.4464 | 8.733703 | 49.05871 | 34.4326 |
| 286 | Trpm4 | 48.12327 | -0.77994 | 0.22576 | 3.45475 | 0.000551 | 0.027975 | 78.57167 | 40.5432 | 45.49611 | 27.8821 |
| 287 | Ptpn22 | 995.3853 | -0.36334 | 0.10527 | 3.45164 | 0.000557 | 0.028204 | 920.7848 | 1538.535 | 715.2325 | 806.989 |
| 288 | Bub1 | 53.39784 | -0.77426 | 0.22438 | 3.45066 | 0.000559 | 0.028212 | 78.30741 | 55.9668 | 45.37712 | 33.94 |

| | | | | | | | | | | |
|---|---|---|---|---|---|---|---|---|---|---|
| 289 | Zbtb18 | 2040.615 | -0.21499 | 0.06247 | 3.44122 | 0.000579 | 0.029118 | 2505.53 | 1763.895 | 2158.837 | 1734.2 |
| 290 | Slamf7 | 2448.85 | 0.38382 | 0.1117 | -3.4361 | 0.00059 | 0.029495 | 1519.819 | 2821.787 | 1984.44 | 3469.35 |
| 291 | Kmo | 1319.686 | -0.42157 | 0.1227 | 3.43561 | 0.000591 | 0.029495 | 2411.543 | 577.1397 | 1798.145 | 491.914 |
| 292 | Oasl1 | 667.148 | 0.648057 | 0.18876 | -3.4332 | 0.000596 | 0.029502 | 563.6693 | 332.158 | 887.6781 | 885.087 |
| 293 | Smagp | 133.8016 | 0.713809 | 0.20791 | -3.4332 | 0.000596 | 0.029502 | 172.8557 | 20.30516 | 284.652 | 57.3934 |
| 294 | Zfp937 | 50.81691 | -0.85939 | 0.25065 | 3.42865 | 0.000607 | 0.029893 | 80.86818 | 50.72438 | 44.02574 | 27.6494 |
| 295 | Pcp4 | 10.21369 | 1.730268 | 0.50476 | -3.4279 | 0.000608 | 0.029893 | 8.804196 | 0.465651 | 31.22223 | 0.36267 |
| 296 | Zbtb32 | 138.8144 | 0.925384 | 0.27035 | -3.4229 | 0.00062 | 0.030346 | 27.89939 | 215.5729 | 54.17298 | 257.612 |
| 297 | Tpm1 | 404.8804 | -0.35945 | 0.10509 | 3.42049 | 0.000625 | 0.030519 | 591.5048 | 271.6763 | 460.7679 | 295.573 |
| 298 | Atl2 | 1420.816 | 0.287819 | 0.08436 | -3.4117 | 0.000646 | 0.031422 | 1204.782 | 1491.735 | 1470.04 | 1516.71 |
| 299 | Reep6 | 39.37197 | 1.898925 | 0.55755 | -3.4058 | 0.00066 | 0.032 | 1.515324 | 63.20651 | 8.933956 | 83.8321 |
| 300 | H2-Eb2 | 3090.926 | -0.35386 | 0.10446 | 3.38734 | 0.000706 | 0.034126 | 2530.114 | 3764.569 | 1977.097 | 4091.93 |
| 301 | Gm14290 | 124.9969 | 0.654472 | 0.19328 | -3.3862 | 0.000709 | 0.03416 | 150.2728 | 50.16433 | 237.8942 | 61.6561 |
| 302 | Fam129b | 69.32233 | -0.87594 | 0.25878 | 3.38491 | 0.000712 | 0.034211 | 50.52069 | 119.9057 | 26.86119 | 80.0017 |
| 303 | Camk2d | 3278.58 | -0.27975 | 0.0828 | 3.37868 | 0.000728 | 0.034885 | 3092.734 | 4190.634 | 2546.468 | 3284.48 |
| 304 | H2-Q7 | 8744.561 | 0.384853 | 0.11421 | -3.3696 | 0.000753 | 0.035936 | 10370.02 | 5402.879 | 13554.88 | 5650.47 |
| 305 | Ccdc176 | 13.615 | -1.58546 | 0.47125 | 3.36438 | 0.000767 | 0.036497 | 27.7098 | 11.18334 | 8.415838 | 7.15101 |
| 306 | Spag9 | 4881.798 | -0.50752 | 0.15088 | 3.36363 | 0.000769 | 0.036497 | 4029.641 | 7415.745 | 2826.822 | 5254.98 |
| 307 | Slc26a10 | 100.0734 | -0.55048 | 0.16383 | 3.35995 | 0.00078 | 0.036761 | 186.8361 | 35.68932 | 126.8817 | 50.8865 |
| 308 | Tma7-ps | 6.022643 | -2.48066 | 0.73831 | 3.35991 | 0.00078 | 0.036761 | 9.104187 | 11.55043 | 0.696431 | 2.73953 |
| 309 | Gm14137 | 188.8643 | 0.715064 | 0.213 | -3.3571 | 0.000788 | 0.037017 | 242.0704 | 44.59827 | 400.6384 | 68.1502 |
| 310 | Ccdc69 | 201.4377 | -0.42859 | 0.12801 | 3.34819 | 0.000813 | 0.038005 | 422.8352 | 37.95887 | 314.0508 | 30.9061 |
| 311 | Chst12 | 163.8159 | -0.50383 | 0.15048 | 3.34813 | 0.000814 | 0.038005 | 289.243 | 76.84573 | 203.9096 | 85.2654 |
| 312 | Lgmn | 2767.183 | 0.381573 | 0.11456 | -3.3307 | 0.000866 | 0.040315 | 3314.439 | 1642.519 | 4323.265 | 1788.51 |
| 313 | Rragb | 135.897 | -0.60573 | 0.1819 | 3.33003 | 0.000868 | 0.040315 | 96.49749 | 215.1091 | 62.6345 | 169.347 |
| 314 | Grpel2 | 1431.531 | 0.231254 | 0.06946 | -3.3292 | 0.000871 | 0.040315 | 862.0787 | 1930.793 | 1011.117 | 1922.13 |
| 315 | Tdg | 1410.792 | -0.36679 | 0.1105 | 3.31921 | 0.000903 | 0.041529 | 1037.231 | 2086.235 | 803.2436 | 1716.46 |
| 316 | Eif3j2 | 155.2338 | 1.582112 | 0.47665 | -3.3192 | 0.000903 | 0.041529 | 27.48221 | 129.2264 | 92.05556 | 372.171 |
| 317 | Ero1lb | 3964.26 | 0.38013 | 0.11474 | -3.3129 | 0.000923 | 0.042246 | 5529.784 | 1513.334 | 7204.158 | 1609.76 |
| 318 | Cacna1h | 180.772 | 0.930038 | 0.28081 | -3.312 | 0.000926 | 0.042246 | 231.0041 | 13.95509 | 447.2887 | 30.8402 |
| 319 | Pdxk | 2362.817 | 0.331383 | 0.10017 | -3.3081 | 0.000939 | 0.042697 | 1382.831 | 3219.631 | 1740.059 | 3108.75 |
| 320 | St14 | 379.1493 | -0.33745 | 0.10209 | 3.30528 | 0.000949 | 0.042996 | 622.2295 | 195.7615 | 491.5649 | 207.041 |
| 321 | mt-Rnr2 | 10701.18 | 0.420538 | 0.12727 | -3.3044 | 0.000952 | 0.043008 | 13763.09 | 4999.128 | 18449.43 | 5593.06 |
| 322 | Ermap | 41.30414 | -0.91847 | 0.27894 | 3.29273 | 0.000992 | 0.044693 | 80.22157 | 23.02973 | 41.91154 | 20.0537 |
| 323 | Lurap1 | 67.28434 | 0.921103 | 0.27986 | -3.2913 | 0.000997 | 0.04478 | 65.48845 | 34.30346 | 126.2902 | 43.0553 |
| 324 | Grap2 | 1344.32 | -0.5992 | 0.18244 | 3.28445 | 0.001022 | 0.045666 | 2668.831 | 563.5033 | 1753.705 | 391.241 |
| 325 | Relt | 323.4974 | 0.338152 | 0.10296 | -3.2842 | 0.001023 | 0.045666 | 346.8666 | 237.9736 | 439.5456 | 269.604 |
| 326 | Ptprs | 658.0212 | -0.69322 | 0.21132 | 3.28042 | 0.001037 | 0.046141 | 498.346 | 1099.625 | 306.2811 | 727.833 |
| 327 | Gopc | 972.5086 | -0.2371 | 0.07236 | 3.27674 | 0.00105 | 0.04652 | 1056.943 | 1005.119 | 895.1705 | 932.802 |
| 328 | Pacs1 | 1338.945 | -0.27913 | 0.0852 | 3.27605 | 0.001053 | 0.04652 | 1923.309 | 986.4582 | 1583.543 | 862.47 |
| 329 | Adap1 | 257.1892 | -0.62364 | 0.1904 | 3.27532 | 0.001055 | 0.04652 | 421.0357 | 170.4356 | 272.0748 | 165.211 |
| 330 | Vill | 87.12433 | 0.65985 | 0.20149 | -3.2748 | 0.001057 | 0.04652 | 110.6347 | 18.02898 | 176.3077 | 43.526 |
| 331 | Slc1a4 | 697.9053 | 2.172468 | 0.66394 | -3.2721 | 0.001068 | 0.046786 | 10.87966 | 1264.871 | 83.1875 | 1432.68 |
| 332 | Prkar2a | 281.2494 | -0.47053 | 0.14389 | 3.26997 | 0.001076 | 0.046786 | 225.7167 | 374.0442 | 163.0404 | 362.196 |
| 333 | Gramd1a | 2115.582 | -0.2635 | 0.08058 | 3.26995 | 0.001076 | 0.046786 | 3162.581 | 1227.785 | 2633.984 | 1437.98 |
| 334 | Gm28042 | 273.907 | -0.55407 | 0.16945 | 3.26972 | 0.001077 | 0.046786 | 483.1164 | 157.4654 | 328.3133 | 126.733 |
| 335 | mt-Tl1 | 55.05891 | 0.877117 | 0.2683 | -3.2691 | 0.001079 | 0.046786 | 56.22956 | 30.46571 | 104.7385 | 28.8019 |
| 336 | Gm15411 | 109.829 | -0.76056 | 0.23284 | 3.26641 | 0.001089 | 0.046985 | 270.8087 | 5.536856 | 158.7905 | 4.18008 |
| 337 | Crim1 | 196.679 | -0.76194 | 0.23327 | 3.2663 | 0.00109 | 0.046985 | 304.1307 | 182.5722 | 177.6094 | 122.404 |
| 338 | Arid5a | 2514.84 | 0.283587 | 0.08685 | -3.2651 | 0.001094 | 0.047042 | 2486.618 | 2535.605 | 3027.918 | 2009.22 |
| 339 | Rgs3 | 155.7535 | -0.51877 | 0.15902 | 3.26237 | 0.001105 | 0.04737 | 325.1349 | 35.85017 | 226.5626 | 35.4661 |
| 340 | Tbc1d30 | 405.2691 | 1.818983 | 0.55801 | -3.2598 | 0.001115 | 0.047667 | 10.67312 | 733.5194 | 48.20575 | 828.678 |
| 341 | Prdx6 | 1367.759 | -0.51237 | 0.15722 | 3.259 | 0.001118 | 0.047667 | 1640.733 | 1365.141 | 1146.898 | 1318.27 |
| 342 | Prkcb | 3752.022 | -0.47537 | 0.14604 | 3.25512 | 0.001133 | 0.048061 | 6753.541 | 1918.948 | 4846.517 | 1489.08 |
| 343 | Kcnk5 | 477.0305 | -0.39204 | 0.12046 | 3.25456 | 0.001136 | 0.048061 | 603.1247 | 419.0122 | 458.1325 | 427.853 |
| 344 | Cdk5rap1 | 208.6818 | 0.79493 | 0.2443 | -3.2539 | 0.001138 | 0.048061 | 166.538 | 161.3282 | 292.2634 | 214.598 |
| 345 | Gpr176 | 22.86018 | 1.591804 | 0.48927 | -3.2534 | 0.00114 | 0.048061 | 8.081815 | 13.77057 | 27.7667 | 41.8216 |
| 346 | Pitpnm2 | 2888.738 | 0.443298 | 0.1363 | -3.2523 | 0.001145 | 0.048061 | 2373.775 | 2920.544 | 3234.259 | 3026.38 |
| 347 | Tusc1 | 22.25976 | 1.267868 | 0.38988 | -3.2519 | 0.001146 | 0.048061 | 12.93751 | 11.01322 | 32.97399 | 32.1143 |
| 348 | Zfp874b | 231.4086 | 0.463225 | 0.14266 | -3.2471 | 0.001166 | 0.048698 | 151.0311 | 294.1865 | 209.0786 | 271.338 |
| 349 | Mybl2 | 133.8623 | -0.48033 | 0.14795 | 3.24659 | 0.001168 | 0.048698 | 156.0463 | 132.2775 | 111.6264 | 135.499 |
| 350 | Mid1 | 612.1837 | 0.922176 | 0.28442 | -3.2423 | 0.001186 | 0.049164 | 325.8053 | 468.2186 | 629.188 | 1025.52 |
| 351 | Cdk17 | 1824.782 | -0.30277 | 0.09351 | 3.23793 | 0.001204 | 0.049759 | 1866.493 | 2090.356 | 1511.826 | 1830.45 |
| 352 | Evi5 | 296.6386 | 0.491196 | 0.15175 | -3.2369 | 0.001208 | 0.049759 | 282.9501 | 167.8226 | 398.6698 | 337.112 |
| 353 | Foxp4 | 1380.352 | -0.27475 | 0.08489 | 3.23655 | 0.00121 | 0.049759 | 1681.445 | 1159.045 | 1388.614 | 1292.3 |
| 354 | Bend3 | 1563.104 | 0.488584 | 0.15103 | -3.2351 | 0.001216 | 0.049872 | 352.3605 | 2797.87 | 495.7872 | 2606.4 |

**Table S2B. List of genes differentially expressed in 3hrs activated B cells of Runx1 cK/O versus Cre control mice. P-values were generated by Wald test and p-adj are BH adjusted (DESeq2).**

| | Associated. Gene.Name | baseMean | log2Fold Change | lfcSE | stat | pvalue | padj | Cre_0 baseMean | Cre_3 baseMean | KO_0 baseMean | KO_3 baseMean |
|---|---|---|---|---|---|---|---|---|---|---|---|
| 1 | Dmwd | 740.7569 | 4.38735 | 0.22123 | -19.8315 | 1.59E-87 | 2.07E-83 | 150.7769 | 26.31985 | 2207.437 | 578.4938 |
| 2 | Cd200 | 782.1183 | 1.8355 | 0.12411 | -14.789 | 1.73E-49 | 1.12E-45 | 596.1455 | 258.2163 | 1347.269 | 926.8421 |
| 3 | Cd47 | 2638.36 | 1.22588 | 0.0913 | -13.4273 | 4.18E-41 | 1.50E-37 | 3201.507 | 1066.609 | 3785.217 | 2500.105 |
| 4 | Dab2ip | 274.2097 | 2.52044 | 0.18781 | -13.4202 | 4.61E-41 | 1.50E-37 | 168.6841 | 37.46776 | 669.1077 | 221.5793 |
| 5 | Gm9855 | 84.55802 | 4.44799 | 0.37375 | -11.901 | 1.17E-32 | 3.05E-29 | 7.69247 | 8.596798 | 102.4293 | 219.5135 |
| 6 | Irf7 | 1046.817 | 1.80791 | 0.15527 | -11.6435 | 2.48E-31 | 5.37E-28 | 1017.043 | 244.6674 | 2061.222 | 864.3334 |
| 7 | Usp18 | 193.0904 | 1.89087 | 0.16648 | -11.358 | 6.77E-30 | 1.26E-26 | 125.6339 | 91.5483 | 210.1824 | 344.997 |
| 8 | Itsn1 | 548.242 | 1.52488 | 0.13589 | -11.2216 | 3.19E-29 | 5.20E-26 | 451.714 | 340.0083 | 413.4089 | 987.8367 |
| 9 | Cables1 | 52.78564 | -2.82557 | 0.25474 | 11.0921 | 1.37E-28 | 1.98E-25 | 11.97511 | 170.1792 | 5.650224 | 23.33802 |
| 10 | Nr4a2 | 701.8206 | 1.5225 | 0.14028 | -10.8534 | 1.92E-27 | 2.50E-24 | 865.5796 | 266.7121 | 905.1257 | 769.8651 |
| 11 | Zbp1 | 786.5503 | 1.07929 | 0.1016 | -10.6233 | 2.32E-26 | 2.75E-23 | 815.9223 | 464.4567 | 880.7793 | 985.0428 |
| 12 | Nedd4 | 2092.746 | 2.06758 | 0.19668 | -10.5123 | 7.58E-26 | 8.22E-23 | 2667.158 | 218.7511 | 4547.004 | 938.0707 |
| 13 | Slc2a6 | 506.8291 | -1.51361 | 0.14493 | 10.4437 | 1.57E-25 | 1.57E-22 | 182.4591 | 1326.727 | 56.21433 | 461.9164 |
| 14 | Gadd45a | 449.2861 | 1.24201 | 0.1225 | -10.1389 | 3.71E-24 | 3.45E-21 | 658.7294 | 120.3924 | 728.8127 | 289.2097 |
| 15 | Ccr6 | 974.1534 | -1.96759 | 0.20041 | 9.81793 | 9.43E-23 | 8.18E-20 | 1238.454 | 1796.21 | 411.5823 | 450.367 |
| 16 | Saraf | 1143.446 | 1.18245 | 0.12238 | -9.66227 | 4.36E-22 | 3.55E-19 | 1119.065 | 401.3009 | 2137.335 | 916.0844 |
| 17 | Trib1 | 1080.816 | 0.8896 | 0.09439 | -9.42444 | 4.32E-21 | 3.31E-18 | 1415.759 | 452.1496 | 1617.781 | 837.5724 |
| 18 | Rtp4 | 199.8168 | 1.70688 | 0.18134 | -9.41249 | 4.85E-21 | 3.50E-18 | 182.7218 | 78.06439 | 280.1963 | 258.2848 |
| 19 | Gm15920 | 73.18057 | 5.0987 | 0.54307 | -9.3886 | 6.08E-21 | 4.17E-18 | 1.595187 | 4.310023 | 72.87643 | 213.9406 |
| 20 | Ifnar1 | 4324.171 | 0.91812 | 0.09808 | -9.36129 | 7.88E-21 | 5.13E-18 | 2694.542 | 2673.909 | 6863.977 | 5064.256 |
| 21 | Ift57 | 99.31614 | 1.93551 | 0.20779 | -9.31491 | 1.22E-20 | 7.57E-18 | 45.03967 | 52.5604 | 94.29607 | 205.3684 |
| 22 | Eya1 | 91.24181 | -1.95416 | 0.21557 | 9.0652 | 1.24E-19 | 7.36E-17 | 155.8559 | 134.2627 | 40.99364 | 33.85509 |
| 23 | Tmem26 | 308.646 | 1.27507 | 0.14086 | -9.05224 | 1.40E-19 | 7.93E-17 | 366.4001 | 87.25683 | 566.4185 | 214.5086 |
| 24 | Tsc22d1 | 1338.839 | 1.07771 | 0.1193 | -9.03341 | 1.66E-19 | 9.03E-17 | 2352.069 | 306.2864 | 2046.361 | 650.639 |
| 25 | Ptpn22 | 995.3853 | -0.92708 | 0.1032 | 8.98347 | 2.62E-19 | 1.37E-16 | 920.7848 | 1538.535 | 715.2325 | 806.9891 |
| 26 | Daam1 | 174.797 | -1.52245 | 0.1699 | 8.96073 | 3.23E-19 | 1.62E-16 | 98.02659 | 410.0408 | 49.52069 | 141.5999 |
| 27 | Lgals3bp | 803.5846 | 1.35623 | 0.15261 | -8.88689 | 6.28E-19 | 3.03E-16 | 863.7509 | 300.3335 | 1275.213 | 775.0411 |
| 28 | Hmgn2-ps1 | 88.90926 | 2.87973 | 0.32928 | -8.74546 | 2.22E-18 | 1.03E-15 | 22.02156 | 24.63402 | 114.722 | 194.2595 |
| 29 | Klhdc2 | 1182.986 | -1.1297 | 0.12986 | 8.69957 | 3.33E-18 | 1.50E-15 | 466.4035 | 2586.131 | 502.6944 | 1176.714 |
| 30 | Mb21d1 | 118.0569 | 1.40087 | 0.16184 | -8.65605 | 4.88E-18 | 2.12E-15 | 30.20043 | 97.58221 | 86.37955 | 258.0656 |
| 31 | Phf11b | 369.3161 | 1.02066 | 0.11804 | -8.64686 | 5.29E-18 | 2.22E-15 | 516.7144 | 127.0823 | 573.8346 | 259.6331 |
| 32 | Gm6548 | 181.3102 | 1.93636 | 0.22472 | -8.61691 | 6.88E-18 | 2.80E-15 | 138.1153 | 57.49982 | 303.8047 | 225.8209 |
| 33 | Gm5806 | 50.29347 | 5.67533 | 0.65934 | -8.60757 | 7.46E-18 | 2.94E-15 | 1.331853 | 1.049371 | 59.00819 | 139.7845 |
| 34 | Gpr171 | 1485.693 | -1.10151 | 0.12834 | 8.58294 | 9.25E-18 | 3.54E-15 | 1658.814 | 2330.591 | 871.8149 | 1081.552 |
| 35 | Dmpk | 214.5662 | 3.2533 | 0.38253 | -8.50463 | 1.82E-17 | 6.78E-15 | 147.0748 | 9.942798 | 594.5257 | 106.7217 |
| 36 | Amigo2 | 529.4001 | 1.07372 | 0.12631 | -8.50066 | 1.89E-17 | 6.82E-15 | 295.8625 | 452.6381 | 413.6759 | 955.4239 |
| 37 | Gm10036 | 68.59307 | 3.60738 | 0.43661 | -8.26227 | 1.43E-16 | 5.03E-14 | 9.856863 | 9.774205 | 114.6733 | 140.0679 |
| 38 | Oas1a | 192.0071 | 1.61697 | 0.19716 | -8.20145 | 2.37E-16 | 8.14E-14 | 220.6267 | 42.94856 | 368.3039 | 136.1492 |
| 39 | Lrrk2 | 6056.895 | -1.85132 | 0.23111 | 8.01045 | 1.14E-15 | 3.82E-13 | 8945.458 | 7822.367 | 5342.392 | 2117.363 |
| 40 | Hsh2d | 399.2783 | 0.78765 | 0.09971 | -7.89929 | 2.80E-15 | 9.13E-13 | 459.0128 | 244.9712 | 470.2532 | 422.8759 |
| 41 | Amd2 | 49.31742 | 2.43974 | 0.31087 | -7.84817 | 4.22E-15 | 1.34E-12 | 5.30795 | 22.84561 | 40.16983 | 128.9463 |
| 42 | Pisd-ps1 | 1992.179 | -0.76312 | 0.0975 | 7.82684 | 5.00E-15 | 1.55E-12 | 3654.134 | 1434.486 | 2036.677 | 843.4177 |
| 43 | Hopx | 174.0787 | 1.52598 | 0.19608 | -7.78239 | 7.12E-15 | 2.16E-12 | 147.7686 | 69.36807 | 275.8305 | 203.3477 |
| 44 | Manea | 1558.941 | -0.94744 | 0.12288 | 7.71014 | 1.26E-14 | 3.72E-12 | 2243.286 | 1600.333 | 1565.87 | 826.2763 |
| 45 | Rbms1 | 1290.266 | 0.59103 | 0.07718 | -7.6581 | 1.89E-14 | 5.46E-12 | 1620.96 | 733.2996 | 1701.718 | 1105.085 |
| 46 | St8sia6 | 268.5616 | -1.3131 | 0.17187 | 7.63992 | 2.17E-14 | 6.15E-12 | 423.4261 | 356.9067 | 152.131 | 141.7824 |
| 47 | Leprotl1 | 553.2531 | 1.13368 | 0.14863 | -7.62777 | 2.39E-14 | 6.62E-12 | 328.0553 | 381.5504 | 658.7951 | 844.6118 |
| 48 | Ifitm3 | 139.173 | 2.01171 | 0.26412 | -7.61663 | 2.60E-14 | 7.06E-12 | 124.6616 | 43.26 | 208.5415 | 180.229 |
| 49 | Lztfl1 | 1235.979 | -0.78287 | 0.10486 | 7.46552 | 8.30E-14 | 2.20E-11 | 910.7191 | 2115.959 | 689.5991 | 1227.639 |
| 50 | Cst7 | 29.65852 | -2.4298 | 0.32879 | 7.39009 | 1.47E-13 | 3.82E-11 | 0.625995 | 96.37865 | 4.38348 | 17.24596 |
| 51 | Il9r | 837.1027 | -2.08083 | 0.28258 | 7.36378 | 1.79E-13 | 4.56E-11 | 2074.102 | 684.8854 | 434.226 | 155.1978 |
| 52 | Klhdc1 | 127.124 | -1.24153 | 0.16873 | 7.35825 | 1.86E-13 | 4.67E-11 | 140.0048 | 177.6927 | 116.2238 | 74.57484 |
| 53 | Oasl1 | 667.148 | 1.3972 | 0.1906 | -7.33039 | 2.29E-13 | 5.64E-11 | 563.6693 | 332.158 | 887.6781 | 885.0867 |
| 54 | Oas3 | 251.9439 | 2.17912 | 0.29865 | -7.29652 | 2.95E-13 | 7.12E-11 | 221.768 | 64.1875 | 417.4723 | 304.3476 |
| 55 | Lgals9 | 1512.211 | 0.77636 | 0.10746 | -7.22472 | 5.02E-13 | 1.19E-10 | 2164.56 | 496.0205 | 2536.989 | 851.2731 |
| 56 | Ppp1r16b | 7093.408 | -0.94721 | 0.13178 | 7.18752 | 6.60E-13 | 1.53E-10 | 6034.002 | 10449.16 | 6492.522 | 5397.944 |
| 57 | Asph | 149.2955 | 1.23993 | 0.17324 | -7.15714 | 8.24E-13 | 1.88E-10 | 93.46409 | 117.2274 | 107.498 | 278.9926 |
| 58 | Rhobtb1 | 73.7577 | 2.07789 | 0.29176 | -7.12202 | 1.06E-12 | 2.35E-10 | 38.45599 | 11.58919 | 191.984 | 53.00157 |
| 59 | Gm6969 | 57.58026 | -4.94325 | 0.70543 | 7.00744 | 2.43E-12 | 5.21E-10 | 168.4737 | 55.49357 | 6.353779 | 0 |
| 60 | Pydc3 | 721.8846 | 1.63566 | 0.23344 | -7.00671 | 2.44E-12 | 5.21E-10 | 898.9619 | 136.6368 | 1418.137 | 433.8022 |
| 61 | Epb4.1l2 | 3609.435 | -0.58642 | 0.08395 | 6.98531 | 2.84E-12 | 5.97E-10 | 3107.285 | 5042.191 | 2933.406 | 3354.858 |
| 62 | Rps13-ps1 | 108.1333 | -5.00298 | 0.71695 | 6.97816 | 2.99E-12 | 6.18E-10 | 209.7466 | 218.4002 | 2.195302 | 2.191212 |
| 63 | Tor3a | 1148.055 | 0.62911 | 0.09028 | -6.96825 | 3.21E-12 | 6.53E-10 | 823.4909 | 1057.784 | 1072.522 | 1638.422 |
| 64 | Kctd14 | 290.8186 | 1.12771 | 0.16282 | -6.92605 | 4.33E-12 | 8.67E-10 | 155.4419 | 208.1909 | 340.5362 | 459.1053 |
| 65 | Abcb1b | 174.522 | -1.62408 | 0.23567 | 6.89121 | 5.53E-12 | 1.09E-09 | 81.3503 | 451.1266 | 22.45042 | 143.1605 |
| 66 | Sdc3 | 1189.424 | 1.75097 | 0.25484 | -6.87091 | 6.38E-12 | 1.24E-09 | 1753.576 | 161.1638 | 2285.214 | 557.7425 |
| 67 | Pole2 | 145.5458 | -1.14361 | 0.16726 | 6.83729 | 8.07E-12 | 1.55E-09 | 70.41973 | 311.8965 | 59.99919 | 139.8676 |
| 68 | Slc2a1 | 1070.411 | 0.71687 | 0.10531 | -6.80706 | 9.96E-12 | 1.88E-09 | 1263.865 | 661.2475 | 1266.8 | 1089.732 |
| 69 | Gpr137b | 459.4167 | -0.82889 | 0.12201 | 6.79339 | 1.10E-11 | 2.04E-09 | 738.5556 | 398.3505 | 477.346 | 223.4144 |

| | | | | | | | | | | | |
|---|---|---|---|---|---|---|---|---|---|---|---|
| 70 | Isg15 | 138.9684 | 1.69858 | 0.2537 | -6.69526 | 2.15E-11 | 3.95E-09 | 87.05666 | 74.75543 | 144.6161 | 249.4455 |
| 71 | Il1r1 | 53.55464 | -2.07235 | 0.31016 | 6.6815 | 2.37E-11 | 4.28E-09 | 7.135937 | 163.5207 | 6.083706 | 37.47822 |
| 72 | Lilrb4 | 774.4591 | -1.11179 | 0.16836 | 6.60367 | 4.01E-11 | 7.15E-09 | 279.4214 | 1784.891 | 213.5896 | 819.9346 |
| 73 | 5430427O1! | 768.8054 | -0.90829 | 0.13814 | 6.5749 | 4.87E-11 | 8.57E-09 | 635.3995 | 1366.912 | 347.251 | 725.659 |
| 74 | Gsn | 690.0347 | -1.00284 | 0.15298 | 6.55535 | 5.55E-11 | 9.64E-09 | 1604.784 | 249.1434 | 781.8312 | 124.3801 |
| 75 | Adck3 | 1925.11 | 0.61906 | 0.0957 | -6.46893 | 9.87E-11 | 1.69E-08 | 2409.915 | 1127.601 | 2429.263 | 1733.66 |
| 76 | Stat1 | 3506.387 | 0.57229 | 0.08858 | -6.46078 | 1.04E-10 | 1.76E-08 | 4365.804 | 2045.797 | 4566.645 | 3047.303 |
| 77 | Ly6a | 1968.773 | 1.37237 | 0.2125 | -6.45822 | 1.06E-10 | 1.77E-08 | 2193.264 | 608.4431 | 3474.503 | 1598.881 |
| 78 | Isg20 | 219.3116 | 0.95206 | 0.14746 | -6.45647 | 1.07E-10 | 1.77E-08 | 229.2643 | 114.5102 | 309.7904 | 223.6815 |
| 79 | Fancc | 320.9376 | -0.74091 | 0.11544 | 6.41804 | 1.38E-10 | 2.25E-08 | 279.9925 | 522.5938 | 169.595 | 311.569 |
| 80 | Cdh24 | 107.4303 | -1.0662 | 0.16625 | 6.41309 | 1.43E-10 | 2.29E-08 | 92.22568 | 193.7134 | 51.46734 | 92.31472 |
| 81 | Mx2 | 187.2386 | 1.49027 | 0.23311 | -6.39292 | 1.63E-10 | 2.58E-08 | 257.8132 | 37.55261 | 344.8007 | 108.7878 |
| 82 | Muc1 | 21.79827 | -2.93175 | 0.46032 | 6.36897 | 1.90E-10 | 2.99E-08 | 29.27763 | 39.43222 | 14.24372 | 4.239527 |
| 83 | Evi5 | 296.6386 | 0.9993 | 0.15723 | -6.35567 | 2.08E-10 | 3.22E-08 | 282.9501 | 167.8226 | 398.6698 | 337.1121 |
| 84 | Mcoln2 | 758.936 | 0.59695 | 0.0941 | -6.34404 | 2.24E-10 | 3.43E-08 | 944.5898 | 437.2732 | 991.7192 | 662.1616 |
| 85 | Gp49a | 639.5648 | -1.16677 | 0.18537 | 6.29409 | 3.09E-10 | 4.68E-08 | 184.7405 | 1555.898 | 130.9582 | 686.6622 |
| 86 | Serpine2 | 52.75259 | 2.1956 | 0.34904 | -6.29038 | 3.17E-10 | 4.74E-08 | 33.56027 | 10.54314 | 115.6149 | 51.2921 |
| 87 | Rab3b | 22.54583 | 2.45708 | 0.39105 | -6.28328 | 3.31E-10 | 4.91E-08 | 5.101835 | 10.60737 | 11.16362 | 63.3105 |
| 88 | Arhgap26 | 1222.626 | -0.91787 | 0.14618 | 6.27917 | 3.40E-10 | 4.98E-08 | 1795.21 | 1052.518 | 1488.857 | 553.9169 |
| 89 | Sema7a | 1065.568 | 0.54403 | 0.08681 | -6.26676 | 3.69E-10 | 5.33E-08 | 784.7047 | 1051.557 | 890.7711 | 1535.238 |
| 90 | Dennd4a | 6281.637 | -0.96087 | 0.15399 | 6.23983 | 4.38E-10 | 6.27E-08 | 5751.694 | 10321.8 | 3778.861 | 5274.197 |
| 91 | Gm4955 | 801.8467 | 1.49331 | 0.24077 | -6.20213 | 5.57E-10 | 7.88E-08 | 1013.993 | 208.7341 | 1384.935 | 599.724 |
| 92 | Pxk | 6180.69 | -0.38119 | 0.0617 | 6.17812 | 6.49E-10 | 9.08E-08 | 7676.398 | 5914.816 | 6590.967 | 4540.579 |
| 93 | Rundc3b | 75.94964 | -1.08077 | 0.17512 | 6.17154 | 6.76E-10 | 9.37E-08 | 34.26294 | 172.3219 | 16.45913 | 80.75463 |
| 94 | Fam53b | 1566.91 | -0.81578 | 0.13285 | 6.14073 | 8.21E-10 | 1.13E-07 | 1572.865 | 2039.759 | 1499.698 | 1155.32 |
| 95 | Clec2i | 1865.326 | -0.6333 | 0.10331 | 6.13021 | 8.78E-10 | 1.19E-07 | 2524.595 | 1720.54 | 2109.059 | 1107.108 |
| 96 | Cybb | 4924.185 | -0.9959 | 0.16356 | 6.08882 | 1.14E-09 | 1.53E-07 | 8681.278 | 2836.128 | 6766.297 | 1413.036 |
| 97 | Irgm1 | 1074.3 | 0.78182 | 0.12903 | -6.05931 | 1.37E-09 | 1.82E-07 | 848.0025 | 910.0029 | 969.0947 | 1570.1 |
| 98 | Six5 | 11.96757 | 3.81851 | 0.6315 | -6.04678 | 1.48E-09 | 1.94E-07 | 1.575054 | 0.585938 | 26.50708 | 19.2022 |
| 99 | St3gal5 | 133.823 | 0.89117 | 0.14748 | -6.04282 | 1.51E-09 | 1.97E-07 | 100.3776 | 105.7315 | 132.0634 | 197.1195 |
| 100 | Ski | 1225.733 | 0.40209 | 0.06669 | -6.02907 | 1.65E-09 | 2.13E-07 | 1075.962 | 914.6709 | 1704.562 | 1207.735 |
| 101 | Txndc16 | 2237.042 | -0.82218 | 0.13661 | 6.01827 | 1.76E-09 | 2.25E-07 | 3334.508 | 1396.666 | 3430.215 | 786.7803 |
| 102 | Plcxd2 | 483.2408 | -0.95766 | 0.1597 | 5.99653 | 2.02E-09 | 2.55E-07 | 500.2545 | 891.4289 | 85.55695 | 455.7228 |
| 103 | Ifi27 | 702.9707 | 0.821 | 0.13776 | -5.95977 | 2.53E-09 | 3.13E-07 | 803.0807 | 344.7175 | 1052.399 | 611.685 |
| 104 | Ifi27l2a | 960.5956 | 1.31553 | 0.22119 | -5.94755 | 2.72E-09 | 3.34E-07 | 1060.678 | 225.777 | 1984.263 | 571.665 |
| 105 | Ncf2 | 2449.462 | 0.49532 | 0.08359 | -5.92567 | 3.11E-09 | 3.75E-07 | 3218.22 | 1044.528 | 4059.425 | 1475.676 |
| 106 | 5031414D1! | 1238.402 | -0.49896 | 0.08436 | 5.91454 | 3.33E-09 | 3.98E-07 | 1839.622 | 752.5967 | 1828.409 | 532.9808 |
| 107 | Abcb1a | 41.14935 | -1.90014 | 0.32165 | 5.90746 | 3.47E-09 | 4.11E-07 | 71.1489 | 59.36171 | 19.09504 | 14.99173 |
| 108 | Slco5a1 | 73.05885 | -2.46713 | 0.41783 | 5.90463 | 3.53E-09 | 4.15E-07 | 203.6684 | 36.26973 | 47.02055 | 5.27671 |
| 109 | Runx1 | 1198.168 | 0.82903 | 0.14084 | -5.88616 | 3.95E-09 | 4.60E-07 | 1431.487 | 596.1719 | 1701.704 | 1063.311 |
| 110 | Gbp6 | 149.6296 | 1.37246 | 0.23403 | -5.86437 | 4.51E-09 | 5.20E-07 | 169.3039 | 60.3174 | 208.7898 | 160.1071 |
| 111 | Vps26b | 123.6154 | 0.98936 | 0.16913 | -5.84978 | 4.92E-09 | 5.62E-07 | 97.14448 | 76.74581 | 165.1382 | 155.4332 |
| 112 | L1cam | 111.5093 | -1.26109 | 0.21658 | 5.82282 | 5.79E-09 | 6.55E-07 | 199.4727 | 75.95983 | 139.5774 | 31.02722 |
| 113 | Hnrnpll | 33.03917 | 1.80068 | 0.31122 | -5.7858 | 7.22E-09 | 8.10E-07 | 24.01289 | 16.58994 | 30.23561 | 61.31824 |
| 114 | Lyl1 | 1072.835 | 0.75641 | 0.13138 | -5.7575 | 8.54E-09 | 9.50E-07 | 1228.697 | 609.2621 | 1420.459 | 1032.922 |
| 115 | Map4k3 | 299.9677 | 0.7575 | 0.13268 | -5.70941 | 1.13E-08 | 1.23E-06 | 289.6948 | 163.5122 | 470.4141 | 276.2495 |
| 116 | Dnah8 | 556.2316 | -1.94137 | 0.3402 | 5.70649 | 1.15E-08 | 1.24E-06 | 899.2069 | 674.4717 | 485.5364 | 165.7114 |
| 117 | Ddx60 | 228.4181 | 1.74564 | 0.30762 | -5.67473 | 1.39E-08 | 1.48E-06 | 340.0618 | 20.76749 | 479.4022 | 73.44081 |
| 118 | Dclre1a | 91.2442 | 1.03656 | 0.18317 | -5.65896 | 1.52E-08 | 1.60E-06 | 71.42852 | 58.64595 | 113.9828 | 120.9195 |
| 119 | Ptpn7 | 719.9187 | 0.63388 | 0.11202 | -5.65842 | 1.53E-08 | 1.60E-06 | 734.2948 | 495.7834 | 879.5692 | 770.0276 |
| 120 | Ifih1 | 142.6112 | 1.25669 | 0.22357 | -5.62097 | 1.90E-08 | 1.98E-06 | 169.2187 | 56.72023 | 206.8175 | 137.6885 |
| 121 | Mcu | 233.8621 | 0.65938 | 0.11747 | -5.61334 | 1.98E-08 | 2.05E-06 | 232.3599 | 164.4468 | 277.9194 | 260.7224 |
| 122 | St6gal1 | 2972.185 | -0.79762 | 0.14213 | 5.61193 | 2.00E-08 | 2.05E-06 | 6042.535 | 1623.506 | 3292.458 | 930.2393 |
| 123 | Cd81 | 1721.878 | 0.78034 | 0.14131 | -5.52204 | 3.35E-08 | 3.41E-06 | 2099.429 | 747.8039 | 2750.409 | 1289.872 |
| 124 | Ifi47 | 618.8432 | 0.84347 | 0.15342 | -5.49784 | 3.84E-08 | 3.85E-06 | 456.5826 | 533.4565 | 522.9265 | 962.4072 |
| 125 | Spin1 | 1694.208 | -0.44042 | 0.08017 | 5.49326 | 3.95E-08 | 3.92E-06 | 1888.371 | 1828.198 | 1714.75 | 1345.513 |
| 126 | Gm3608 | 72.71769 | -2.53758 | 0.46251 | 5.48652 | 4.10E-08 | 4.04E-06 | 117.8018 | 129.6811 | 24.0696 | 19.31827 |
| 127 | BC017643 | 1480.564 | -0.71505 | 0.13061 | 5.47468 | 4.38E-08 | 4.29E-06 | 2252.581 | 1166.334 | 1794.453 | 708.8887 |
| 128 | Pear1 | 536.0533 | 0.78028 | 0.14265 | -5.4698 | 4.51E-08 | 4.38E-06 | 621.511 | 211.6469 | 946.4019 | 364.6535 |
| 129 | Mical1 | 1424.822 | -0.53758 | 0.09853 | 5.45626 | 4.86E-08 | 4.69E-06 | 2296.964 | 954.1186 | 1791.277 | 656.9271 |
| 130 | Psen2 | 929.1939 | 0.60892 | 0.11182 | -5.4456 | 5.16E-08 | 4.94E-06 | 929.0957 | 646.2265 | 1152.966 | 988.4871 |
| 131 | Parp12 | 39.20666 | 1.72591 | 0.31716 | -5.44167 | 5.28E-08 | 5.02E-06 | 47.55656 | 12.15298 | 54.75168 | 42.36543 |
| 132 | Ifit3 | 181.9063 | 1.28429 | 0.2372 | -5.41434 | 6.15E-08 | 5.80E-06 | 250.4984 | 46.28145 | 316.4038 | 114.4417 |
| 133 | Dhtkd1 | 50.11996 | -1.42043 | 0.26297 | 5.40143 | 6.61E-08 | 6.19E-06 | 55.02145 | 77.05572 | 40.31732 | 28.08534 |
| 134 | Zfp36l2 | 5248.21 | 0.82123 | 0.15272 | -5.37735 | 7.56E-08 | 6.98E-06 | 8538.571 | 1167.608 | 9214.88 | 2071.783 |
| 135 | Smagp | 133.8016 | 1.48628 | 0.27752 | -5.3556 | 8.53E-08 | 7.82E-06 | 172.8557 | 20.30516 | 284.652 | 57.3934 |
| 136 | Irf5 | 3126.734 | 0.47256 | 0.08878 | -5.32282 | 1.02E-07 | 9.30E-06 | 3266.609 | 2304.631 | 3734.529 | 3201.165 |
| 137 | Ifit1 | 72.67536 | 1.62273 | 0.30514 | -5.31804 | 1.05E-07 | 9.48E-06 | 45.20418 | 41.10215 | 73.48782 | 130.9073 |
| 138 | Slc9a3r1 | 968.1224 | 0.52497 | 0.09909 | -5.29782 | 1.17E-07 | 1.05E-05 | 1008.34 | 723.3942 | 1098.403 | 1042.353 |
| 139 | Ptpn14 | 326.1321 | 1.86245 | 0.35238 | -5.28532 | 1.25E-07 | 1.12E-05 | 700.5816 | 12.74114 | 539.3294 | 51.87622 |
| 140 | Rell1 | 201.6663 | -0.91714 | 0.17355 | 5.28443 | 1.26E-07 | 1.12E-05 | 420.5699 | 127.4846 | 191.2413 | 67.36961 |
| 141 | Il10 | 100.8163 | 1.59002 | 0.30123 | -5.27835 | 1.30E-07 | 1.15E-05 | 73.84632 | 56.25124 | 99.00511 | 174.1626 |
| 142 | Trim30d | 1087.523 | 1.09874 | 0.20825 | -5.27599 | 1.32E-07 | 1.15E-05 | 1251.215 | 529.8008 | 1422.081 | 1146.995 |

| | | | | | | | | | | | |
|---|---|---|---|---|---|---|---|---|---|---|---|
| 143 | Whrn | 770.5132 | 0.67644 | 0.12823 | -5.27505 | 1.33E-07 | 1.15E-05 | 851.1784 | 381.2769 | 1239.761 | 609.8368 |
| 144 | Slfn1 | 40.1616 | 1.94526 | 0.36946 | -5.26511 | 1.40E-07 | 1.21E-05 | 29.23779 | 14.4912 | 55.75386 | 61.16354 |
| 145 | Plekhm1 | 1918.909 | -0.50392 | 0.09596 | 5.25146 | 1.51E-07 | 1.29E-05 | 3068.719 | 1140.833 | 2662.509 | 803.574 |
| 146 | Fam21 | 2692.044 | -0.34458 | 0.0658 | 5.23675 | 1.63E-07 | 1.39E-05 | 2419.177 | 3355.115 | 2351.34 | 2642.543 |
| 147 | Tnfrsf13c | 4888.305 | 0.47905 | 0.09191 | -5.21226 | 1.87E-07 | 1.57E-05 | 2912.359 | 5450.502 | 3585.9 | 7604.461 |
| 148 | Sfxn3 | 637.5344 | -0.51775 | 0.09933 | 5.21214 | 1.87E-07 | 1.57E-05 | 856.0431 | 419.6096 | 981.3389 | 293.1461 |
| 149 | Mov10 | 448.5023 | 0.51709 | 0.09929 | -5.20782 | 1.91E-07 | 1.59E-05 | 472.7911 | 328.5079 | 523.6021 | 469.1079 |
| 150 | Gm7592 | 191.7882 | 0.96412 | 0.18543 | -5.19927 | 2.00E-07 | 1.66E-05 | 217.454 | 97.74458 | 260.3172 | 191.6371 |
| 151 | Ublcp1 | 1129.293 | 0.5207 | 0.10037 | -5.18791 | 2.13E-07 | 1.75E-05 | 1637.854 | 418.7633 | 1858.196 | 602.357 |
| 152 | Btla | 6695.415 | -0.51331 | 0.09933 | 5.1677 | 2.37E-07 | 1.94E-05 | 12383.78 | 3739.1 | 8043.244 | 2615.533 |
| 153 | Glrx | 128.5064 | 0.91349 | 0.17819 | -5.12654 | 2.95E-07 | 2.40E-05 | 159.2337 | 57.25674 | 190.0722 | 107.463 |
| 154 | Dos | 131.6539 | 0.88688 | 0.17307 | -5.12434 | 2.99E-07 | 2.41E-05 | 60.74206 | 131.2835 | 90.13052 | 244.4594 |
| 155 | Trim30a | 2904.783 | 0.96875 | 0.18944 | -5.11386 | 3.16E-07 | 2.54E-05 | 3603.758 | 1324.344 | 4078.551 | 2612.477 |
| 156 | Amica1 | 152.824 | 1.03363 | 0.20276 | -5.09779 | 3.44E-07 | 2.75E-05 | 257.1445 | 37.65519 | 237.2504 | 79.24601 |
| 157 | Gimap4 | 4441.047 | 0.73902 | 0.14508 | -5.0939 | 3.51E-07 | 2.78E-05 | 2972.416 | 4048.132 | 3963.477 | 6780.162 |
| 158 | Tmc6 | 1937.5 | 0.47996 | 0.09444 | -5.08226 | 3.73E-07 | 2.94E-05 | 1914.607 | 1619.451 | 1953.056 | 2262.885 |
| 159 | Oas2 | 484.9155 | 1.4146 | 0.2799 | -5.05392 | 4.33E-07 | 3.40E-05 | 620.0567 | 73.70384 | 1043.376 | 202.5253 |
| 160 | Map3k8 | 1166.311 | -0.54992 | 0.10924 | 5.03426 | 4.80E-07 | 3.74E-05 | 1484.875 | 1073.221 | 1375.23 | 731.9164 |
| 161 | Fam83d | 47.85914 | -1.48978 | 0.29603 | 5.03253 | 4.84E-07 | 3.75E-05 | 80.7243 | 44.73737 | 50.90122 | 15.07367 |
| 162 | Acad8 | 106.2769 | 0.93093 | 0.18505 | -5.03078 | 4.88E-07 | 3.76E-05 | 70.33056 | 83.60098 | 109.3814 | 161.7948 |
| 163 | Irf9 | 1849.262 | 0.45872 | 0.09123 | -5.02847 | 4.94E-07 | 3.79E-05 | 1998.032 | 1354.944 | 2180.714 | 1863.36 |
| 164 | Neat1 | 4235.338 | -0.51496 | 0.10271 | 5.01385 | 5.34E-07 | 4.06E-05 | 3710.535 | 5681.569 | 3578.225 | 3971.024 |
| 165 | 1700017B05 | 2102.743 | 0.46067 | 0.0919 | -5.01264 | 5.37E-07 | 4.06E-05 | 2318.199 | 1382.47 | 2805.83 | 1904.474 |
| 166 | Ifit2 | 290.2247 | 1.34579 | 0.26869 | -5.00865 | 5.48E-07 | 4.13E-05 | 359.2175 | 82.65597 | 503.5903 | 215.435 |
| 167 | Tjp3 | 169.6555 | -0.82079 | 0.16391 | 5.00749 | 5.51E-07 | 4.13E-05 | 226.1977 | 161.2043 | 200.1078 | 91.11233 |
| 168 | Rap1gap | 189.8781 | 1.25097 | 0.25027 | -4.99849 | 5.78E-07 | 4.30E-05 | 152.451 | 80.29713 | 331.6749 | 195.0893 |
| 169 | Ckap4 | 460.0014 | 0.57469 | 0.11546 | -4.97745 | 6.44E-07 | 4.77E-05 | 588.2347 | 198.1105 | 757.4922 | 296.1683 |
| 170 | 1700016P03 | 73.83288 | 1.51522 | 0.30516 | -4.96539 | 6.86E-07 | 5.04E-05 | 30.27241 | 55.31262 | 46.63189 | 163.1146 |
| 171 | Igtp | 441.5015 | 0.62143 | 0.12539 | -4.95612 | 7.19E-07 | 5.26E-05 | 397.7312 | 358.5774 | 455.0787 | 554.6189 |
| 172 | Rab37 | 144.2027 | 0.8086 | 0.16333 | -4.95085 | 7.39E-07 | 5.37E-05 | 142.8377 | 100.9399 | 155.8178 | 177.2153 |
| 173 | Gypc | 333.1138 | 0.69367 | 0.14065 | -4.93192 | 8.14E-07 | 5.88E-05 | 274.4039 | 270.7424 | 348.0627 | 439.2462 |
| 174 | Pydc4 | 1305.641 | 1.51837 | 0.30792 | -4.93102 | 8.18E-07 | 5.88E-05 | 1922.768 | 122.7253 | 2812.094 | 364.9779 |
| 175 | Gyk | 453.2265 | -0.70133 | 0.14257 | 4.91912 | 8.69E-07 | 6.19E-05 | 174.5701 | 918.5601 | 157.4227 | 562.3531 |
| 176 | Gm15382 | 35.10391 | -1.34175 | 0.27286 | 4.91731 | 8.77E-07 | 6.21E-05 | 34.1564 | 56.12917 | 28.63074 | 21.49935 |
| 177 | Kbtbd11 | 859.1567 | 1.0141 | 0.20666 | -4.90716 | 9.24E-07 | 6.50E-05 | 1431.52 | 84.88468 | 1746.74 | 173.4826 |
| 178 | Arc | 34.7061 | 1.58515 | 0.32325 | -4.90381 | 9.40E-07 | 6.58E-05 | 41.63235 | 13.73002 | 39.88091 | 43.58114 |
| 179 | Socs5 | 605.4593 | -0.6763 | 0.13819 | 4.89405 | 9.88E-07 | 6.88E-05 | 430.9839 | 936.998 | 469.3728 | 584.4824 |
| 180 | Junb | 11581.11 | 0.87157 | 0.17861 | -4.87982 | 1.06E-06 | 7.35E-05 | 18434.23 | 2639.741 | 20389.76 | 4860.714 |
| 181 | Capg | 5337.819 | 0.95618 | 0.19622 | -4.87301 | 1.10E-06 | 7.57E-05 | 3323.026 | 4227.541 | 5527.578 | 8273.132 |
| 182 | Nckap1 | 26.95324 | 1.79764 | 0.36941 | -4.86618 | 1.14E-06 | 7.80E-05 | 6.077149 | 19.44806 | 12.59796 | 69.6898 |
| 183 | Elk3 | 633.8462 | 0.47773 | 0.09822 | -4.86398 | 1.15E-06 | 7.84E-05 | 1092.017 | 292.7896 | 742.0572 | 408.5213 |
| 184 | Atf3 | 2697.363 | 0.89627 | 0.18471 | -4.85225 | 1.22E-06 | 8.28E-05 | 3914.076 | 619.5119 | 5095.046 | 1160.819 |
| 185 | Itga6 | 439.676 | -0.90112 | 0.1858 | 4.8501 | 1.23E-06 | 8.33E-05 | 510.36 | 519.2084 | 453.4604 | 275.675 |
| 186 | Dhx58 | 430.1777 | 0.96816 | 0.19968 | -4.84853 | 1.24E-06 | 8.35E-05 | 527.3331 | 183.7791 | 647.1237 | 362.475 |
| 187 | Nek7 | 2054.656 | 0.40105 | 0.08281 | -4.84305 | 1.28E-06 | 8.54E-05 | 1908.793 | 1818.48 | 2090.26 | 2401.091 |
| 188 | Fam69a | 611.8732 | -0.44251 | 0.09142 | 4.84022 | 1.30E-06 | 8.61E-05 | 797.8157 | 629.6696 | 557.5365 | 462.4711 |
| 189 | Srgap3 | 44.04232 | -1.70049 | 0.35139 | 4.83926 | 1.30E-06 | 8.61E-05 | 73.32008 | 37.64563 | 54.52602 | 10.67756 |
| 190 | Oasl2 | 204.2372 | 1.23547 | 0.25577 | -4.83038 | 1.36E-06 | 8.96E-05 | 242.3238 | 63.53888 | 357.6379 | 153.4482 |
| 191 | Plin3 | 297.6177 | -0.69601 | 0.14438 | 4.82079 | 1.43E-06 | 9.33E-05 | 295.9803 | 374.7132 | 288.0987 | 231.6786 |
| 192 | Btnl2 | 346.46 | 0.70188 | 0.14561 | -4.82029 | 1.43E-06 | 9.33E-05 | 277.0635 | 289.8525 | 346.0581 | 472.8658 |
| 193 | Pltp | 87.65011 | -1.3733 | 0.28501 | 4.81835 | 1.45E-06 | 9.35E-05 | 162.3076 | 49.89496 | 119.5329 | 18.86503 |
| 194 | Zfp608 | 2998.11 | -0.8331 | 0.17292 | 4.81787 | 1.45E-06 | 9.35E-05 | 2494.69 | 4585.485 | 2353.556 | 2558.708 |
| 195 | Egln3 | 22.95046 | -1.66996 | 0.34702 | 4.81227 | 1.49E-06 | 9.57E-05 | 23.57036 | 40.84648 | 15.19598 | 12.18902 |
| 196 | Epsti1 | 633.5069 | 0.65938 | 0.13721 | -4.80568 | 1.54E-06 | 9.84E-05 | 1091.112 | 118.863 | 1136.168 | 187.8852 |
| 197 | Pacsin1 | 428.3699 | 0.67859 | 0.14159 | -4.79269 | 1.65E-06 | 0.0001 | 526.1031 | 222.1891 | 608.3097 | 356.8777 |
| 198 | Lphn2 | 237.5704 | -1.15525 | 0.24217 | 4.77051 | 1.84E-06 | 0.00012 | 100.5672 | 539.2684 | 71.82494 | 238.6211 |
| 199 | Ehd4 | 1273.027 | 0.78859 | 0.16624 | -4.74371 | 2.10E-06 | 0.00013 | 1712.107 | 389.6792 | 2311.932 | 678.3913 |
| 200 | Tmem154 | 537.5289 | 0.81242 | 0.17229 | -4.71528 | 2.41E-06 | 0.00015 | 642.5037 | 172.6757 | 1029.241 | 305.6949 |
| 201 | Sdc4 | 1926.589 | -0.54428 | 0.11576 | 4.70163 | 2.58E-06 | 0.00016 | 2864.187 | 1673.018 | 2022.323 | 1146.827 |
| 202 | C330027C09 | 132.302 | -0.81435 | 0.17343 | 4.69557 | 2.66E-06 | 0.00016 | 171.2735 | 150.5858 | 122.2063 | 85.14212 |
| 203 | Rps6ka1 | 1283.27 | 0.41484 | 0.0886 | -4.68237 | 2.84E-06 | 0.00017 | 1322.439 | 1097.893 | 1245.519 | 1467.23 |
| 204 | Aff3 | 2581.216 | -0.72461 | 0.1548 | 4.68096 | 2.86E-06 | 0.00017 | 3989.235 | 1823.438 | 3413.268 | 1098.921 |
| 205 | Ctns | 404.8013 | -0.59333 | 0.12705 | 4.66998 | 3.01E-06 | 0.00018 | 361.9647 | 557.4858 | 330.6097 | 369.1449 |
| 206 | Xaf1 | 398.5604 | 1.16093 | 0.24869 | -4.66827 | 3.04E-06 | 0.00018 | 456.9152 | 170.4056 | 579.272 | 387.6488 |
| 207 | Gfod1 | 274.0259 | 0.81224 | 0.17419 | -4.66283 | 3.12E-06 | 0.00019 | 301.8292 | 143.8955 | 396.3521 | 254.0268 |
| 208 | Slc25a20 | 363.5084 | 0.71165 | 0.15331 | -4.64184 | 3.45E-06 | 0.00021 | 416.2528 | 218.7836 | 458.7286 | 360.2687 |
| 209 | Rsl1 | 198.234 | -0.57984 | 0.12492 | 4.64179 | 3.45E-06 | 0.00021 | 125.6651 | 341.8521 | 97.12333 | 228.2954 |
| 210 | Ablim1 | 10258.82 | -0.52607 | 0.1141 | 4.61071 | 4.01E-06 | 0.00024 | 12260.06 | 10673.59 | 10701.75 | 7399.903 |
| 211 | Bcl2l11 | 2991.385 | 0.48721 | 0.10568 | -4.61023 | 4.02E-06 | 0.00024 | 1944.933 | 3273.133 | 2155.482 | 4591.994 |
| 212 | Slc44a2 | 4018.062 | -0.40193 | 0.08724 | 4.60736 | 4.08E-06 | 0.00024 | 5949.668 | 2691.304 | 5394.418 | 2036.856 |
| 213 | Spn | 203.1635 | 0.76304 | 0.16576 | -4.60325 | 4.16E-06 | 0.00024 | 219.7011 | 130.9172 | 236.3951 | 225.6408 |
| 214 | Cmpk2 | 241.665 | 1.15121 | 0.25026 | -4.60006 | 4.22E-06 | 0.00025 | 396.6652 | 52.12202 | 399.2517 | 118.621 |
| 215 | Gimap3 | 1852.946 | 0.57858 | 0.12609 | -4.58864 | 4.46E-06 | 0.00026 | 1720.592 | 1490.235 | 1970.959 | 2229.996 |

| | | | | | | | | | | | |
|---|---|---|---|---|---|---|---|---|---|---|---|
| 216 | Rara | 1511.476 | 0.40281 | 0.08794 | -4.58075 | 4.63E-06 | 0.00027 | 1503.917 | 1369.652 | 1358.664 | 1813.668 |
| 217 | Fgr | 504.1348 | 0.66581 | 0.14554 | -4.57466 | 4.77E-06 | 0.00027 | 811.2669 | 131.9676 | 860.8404 | 212.4641 |
| 218 | Fosl2 | 2081.551 | 1.15514 | 0.25374 | -4.55239 | 5.30E-06 | 0.0003 | 390.1249 | 2232.674 | 645.7286 | 5057.679 |
| 219 | Rgcc | 216.5396 | 1.12443 | 0.24733 | -4.54623 | 5.46E-06 | 0.00031 | 460.8565 | 29.38976 | 310.6838 | 65.22833 |
| 220 | Mapre2 | 1906.168 | 0.37993 | 0.08366 | -4.54157 | 5.58E-06 | 0.00031 | 1257.628 | 2141.207 | 1436.676 | 2789.164 |
| 221 | Ssfa2 | 3937.782 | 0.55599 | 0.12265 | -4.53318 | 5.81E-06 | 0.00033 | 1546.069 | 5024.911 | 1778.81 | 7401.336 |
| 222 | Phf11a | 61.11398 | 1.35312 | 0.29891 | -4.52676 | 5.99E-06 | 0.00033 | 85.87908 | 12.15076 | 114.369 | 32.05711 |
| 223 | Nr4a1 | 15030.67 | 0.72888 | 0.16109 | -4.52466 | 6.05E-06 | 0.00034 | 20021.9 | 7564.559 | 19944.1 | 12592.14 |
| 224 | Tgfb1 | 4336.828 | 0.62542 | 0.13839 | -4.51928 | 6.20E-06 | 0.00034 | 3273.817 | 4153.268 | 3496.148 | 6424.08 |
| 225 | Nrip1 | 913.3527 | -0.6545 | 0.14511 | 4.51042 | 6.47E-06 | 0.00036 | 898.5021 | 1308.135 | 619.388 | 827.3856 |
| 226 | Iqsec2 | 135.566 | 0.90019 | 0.19962 | -4.50945 | 6.50E-06 | 0.00036 | 50.64694 | 137.4923 | 95.89029 | 258.2346 |
| 227 | Pde7a | 2152.833 | -0.40021 | 0.08934 | 4.47979 | 7.47E-06 | 0.00041 | 3861.933 | 1251.829 | 2549.685 | 947.884 |
| 228 | Adk | 221.1175 | 0.61802 | 0.13895 | -4.44767 | 8.68E-06 | 0.00047 | 186.5539 | 156.4787 | 301.5542 | 239.883 |
| 229 | Itgb3bp | 121.8906 | 0.82671 | 0.18686 | -4.42423 | 9.68E-06 | 0.00053 | 143.4585 | 63.05744 | 167.8526 | 113.1938 |
| 230 | Cenpe | 128.3162 | -0.995 | 0.22575 | 4.40761 | 1.05E-05 | 0.00056 | 213.6954 | 113.5339 | 130.1578 | 55.8776 |
| 231 | Mki67 | 840.0968 | -0.97964 | 0.22232 | 4.40644 | 1.05E-05 | 0.00057 | 1429.355 | 742.1782 | 817.0144 | 371.8393 |
| 232 | Gripap1 | 3748.843 | 0.44329 | 0.1007 | -4.40198 | 1.07E-05 | 0.00057 | 3228.16 | 3536.671 | 3415.516 | 4815.026 |
| 233 | Eif2ak2 | 641.1723 | 1.31236 | 0.29816 | -4.40155 | 1.07E-05 | 0.00057 | 604.8263 | 275.8881 | 979.4748 | 704.4998 |
| 234 | Gphn | 399.3972 | 0.5975 | 0.13598 | -4.39402 | 1.11E-05 | 0.00059 | 372.2465 | 329.0788 | 399.0983 | 497.1653 |
| 235 | Fam46a | 441.6413 | 0.77475 | 0.17661 | -4.38679 | 1.15E-05 | 0.00061 | 571.8793 | 173.0649 | 724.7414 | 296.8794 |
| 236 | Rasa4 | 309.7848 | 0.6914 | 0.15786 | -4.37986 | 1.19E-05 | 0.00063 | 306.6126 | 158.9897 | 512.8601 | 260.6767 |
| 237 | Tpp1 | 2143.887 | -0.34873 | 0.0797 | 4.37562 | 1.21E-05 | 0.00064 | 1752.28 | 2896.859 | 1651.759 | 2274.649 |
| 238 | Slfn8 | 1027.603 | 1.14144 | 0.26106 | -4.37236 | 1.23E-05 | 0.00064 | 1347.828 | 390.6072 | 1494.22 | 877.7561 |
| 239 | Tgfb3 | 161.194 | -0.73922 | 0.1696 | 4.35864 | 1.31E-05 | 0.00068 | 180.451 | 225.1897 | 104.5379 | 134.5975 |
| 240 | D8Ertd82e | 1337.383 | -0.42374 | 0.09722 | 4.3585 | 1.31E-05 | 0.00068 | 814.8838 | 2227.569 | 647.679 | 1659.4 |
| 241 | Daxx | 1206.356 | 0.33289 | 0.07641 | -4.35648 | 1.32E-05 | 0.00068 | 1083.195 | 1105.749 | 1242.193 | 1394.286 |
| 242 | Fbxo48 | 42.73536 | -1.47509 | 0.33884 | 4.3534 | 1.34E-05 | 0.00069 | 54.08346 | 61.55861 | 34.39056 | 20.90883 |
| 243 | Slc37a3 | 428.2352 | -0.4869 | 0.11223 | 4.33846 | 1.43E-05 | 0.00074 | 393.8848 | 575.6344 | 333.3086 | 410.113 |
| 244 | Il18 | 126.6362 | 1.72632 | 0.39896 | -4.32706 | 1.51E-05 | 0.00077 | 168.9988 | 4.738025 | 314.2087 | 18.5995 |
| 245 | Foxn3 | 1630.607 | -1.27767 | 0.29561 | 4.32213 | 1.55E-05 | 0.00079 | 1801.401 | 2593.65 | 1086.176 | 1041.2 |
| 246 | Dusp5 | 1807.613 | 0.79301 | 0.18474 | -4.29257 | 1.77E-05 | 0.00089 | 2123.268 | 989.6614 | 2393.166 | 1724.356 |
| 247 | Sema4d | 3104.353 | -0.63653 | 0.14852 | 4.2858 | 1.82E-05 | 0.00092 | 5528.43 | 1870.976 | 3818.352 | 1199.653 |
| 249 | Camk2d | 3278.58 | -0.35076 | 0.08201 | 4.27676 | 1.90E-05 | 0.00095 | 3092.734 | 4190.634 | 2546.468 | 3284.482 |
| 250 | Casp8 | 1341.309 | 0.3653 | 0.08548 | -4.27349 | 1.92E-05 | 0.00096 | 1684.518 | 781.4675 | 1892.574 | 1006.675 |
| 251 | Ptpn1 | 5060.422 | -0.37076 | 0.08683 | 4.27009 | 1.95E-05 | 0.00097 | 3373.642 | 7580.672 | 3428.721 | 5858.653 |
| 252 | Ide | 3879.819 | -0.85756 | 0.20089 | 4.26884 | 1.96E-05 | 0.00098 | 1153.773 | 8704.888 | 894.5652 | 4766.05 |
| 253 | Myo1g | 5445.197 | -0.23289 | 0.05473 | 4.25493 | 2.09E-05 | 0.00104 | 6314.102 | 5068.882 | 6086.17 | 4311.636 |
| 254 | Aftph | 2575.715 | -0.19747 | 0.0465 | 4.24622 | 2.17E-05 | 0.00107 | 2373.62 | 3046.495 | 2227.25 | 2655.496 |
| 255 | Zfp597 | 421.7306 | -0.64891 | 0.15329 | 4.2332 | 2.30E-05 | 0.00113 | 408.4838 | 567.2338 | 351.6462 | 359.5587 |
| 256 | Kcnh4 | 359.1391 | -0.67199 | 0.15884 | 4.23059 | 2.33E-05 | 0.00114 | 76.59393 | 777.297 | 96.16623 | 486.4991 |
| 257 | Vill | 87.12433 | 1.16683 | 0.27623 | -4.22417 | 2.40E-05 | 0.00117 | 110.6347 | 18.02898 | 176.3077 | 43.52603 |
| 258 | Serpini1 | 290.2643 | 0.83857 | 0.19875 | -4.21917 | 2.45E-05 | 0.00119 | 530.2481 | 37.66334 | 525.9131 | 67.2325 |
| 259 | Arhgap31 | 275.8913 | -1.01124 | 0.23989 | 4.21546 | 2.49E-05 | 0.00121 | 555.6837 | 122.7333 | 365.3306 | 59.81746 |
| 260 | Eps8l1 | 315.2604 | -1.35139 | 0.32076 | 4.21309 | 2.52E-05 | 0.00121 | 721.729 | 185.9975 | 283.1771 | 70.13786 |
| 261 | Pik3ip1 | 952.5659 | -0.63954 | 0.15206 | 4.20582 | 2.60E-05 | 0.00125 | 943.4732 | 1248.034 | 819.0228 | 799.734 |
| 262 | Rin3 | 1258.854 | 0.51615 | 0.12279 | -4.20344 | 2.63E-05 | 0.00126 | 2263.976 | 247.3766 | 2169.157 | 354.9065 |
| 263 | BC094916 | 409.5891 | 1.10412 | 0.26296 | -4.19881 | 2.68E-05 | 0.00128 | 489.7458 | 185.1701 | 558.9172 | 404.5233 |
| 264 | Ubxn11 | 330.9738 | 0.60091 | 0.14314 | -4.19798 | 2.69E-05 | 0.00128 | 349.2256 | 231.6574 | 391.2154 | 351.7966 |
| 265 | Trim12c | 1167.472 | 0.38581 | 0.09195 | -4.19589 | 2.72E-05 | 0.00129 | 1248.724 | 906.3423 | 1330.733 | 1184.088 |
| 266 | Pacsin2 | 1806.326 | 0.34193 | 0.08152 | -4.19465 | 2.73E-05 | 0.00129 | 1159.605 | 2118.414 | 1262.076 | 2685.208 |
| 267 | Gm12250 | 195.5429 | 0.76529 | 0.18304 | -4.18105 | 2.90E-05 | 0.00136 | 80.97951 | 221.4511 | 101.4073 | 378.3337 |
| 268 | Osbpl9 | 3211.049 | -0.25207 | 0.06064 | 4.15661 | 3.23E-05 | 0.00151 | 3186.269 | 3342.07 | 3510.482 | 2805.376 |
| 269 | Pxdc1 | 1373.459 | 0.77758 | 0.18771 | -4.14249 | 3.44E-05 | 0.0016 | 1260.213 | 1002.967 | 1500.8 | 1729.855 |
| 270 | Scpep1 | 1358.91 | -0.34523 | 0.08342 | 4.13873 | 3.49E-05 | 0.00162 | 1662.936 | 1371.52 | 1320.545 | 1080.64 |
| 271 | Fam214a | 1406.175 | -0.82177 | 0.19882 | 4.13331 | 3.58E-05 | 0.00166 | 2662.42 | 570.6492 | 2071.311 | 320.3207 |
| 272 | Top2a | 763.1014 | -0.53733 | 0.13002 | 4.1327 | 3.59E-05 | 0.00166 | 1002.767 | 815.3482 | 672.982 | 561.3083 |
| 273 | Nusap1 | 143.7237 | -0.69364 | 0.16793 | 4.13039 | 3.62E-05 | 0.00167 | 214.9866 | 124.3564 | 159.071 | 76.48098 |
| 274 | Ddx26b | 888.2362 | -0.64641 | 0.15678 | 4.1231 | 3.74E-05 | 0.00171 | 1379.761 | 583.7954 | 1218.239 | 371.1484 |
| 275 | Akt3 | 804.1361 | -0.61951 | 0.1505 | 4.11633 | 3.85E-05 | 0.00174 | 1291.021 | 531.9071 | 1049.114 | 344.5021 |
| 276 | Stk38l | 622.7715 | 0.50247 | 0.12223 | -4.11087 | 3.94E-05 | 0.00178 | 429.6427 | 664.6018 | 454.623 | 942.2186 |
| 277 | Ehd3 | 167.0023 | -1.10898 | 0.27008 | 4.10616 | 4.02E-05 | 0.00181 | 438.6983 | 58.49175 | 144.4347 | 26.3845 |
| 278 | Epn2 | 90.97733 | 0.87934 | 0.21438 | -4.10168 | 4.10E-05 | 0.00184 | 75.94101 | 49.05292 | 149.1058 | 89.80956 |
| 279 | Pyhin1 | 2615.327 | 0.81612 | 0.19936 | -4.09359 | 4.25E-05 | 0.00189 | 4256.554 | 588.0804 | 4573.142 | 1043.532 |
| 280 | Cers6 | 120.5132 | -0.69553 | 0.17025 | 4.08528 | 4.40E-05 | 0.00196 | 160.9115 | 135.9773 | 101.8569 | 83.30701 |
| 281 | Kremen1 | 485.2043 | -0.38424 | 0.09408 | 4.0841 | 4.42E-05 | 0.00196 | 533.6048 | 557.3306 | 424.457 | 425.4248 |
| 282 | Fam134b | 1930.379 | 0.38239 | 0.0938 | -4.07678 | 4.57E-05 | 0.00202 | 2492.529 | 1134.151 | 2614.09 | 1480.748 |
| 283 | 4931428F04 | 94.28003 | -0.73808 | 0.18154 | 4.06558 | 4.79E-05 | 0.00211 | 94.95316 | 121.3609 | 88.40498 | 72.40105 |
| 284 | Hs3st3b1 | 62.44958 | 0.90552 | 0.22288 | -4.06288 | 4.85E-05 | 0.00213 | 54.70835 | 42.30988 | 72.02396 | 80.75614 |
| 285 | Tgtp2 | 119.8311 | 0.96915 | 0.23869 | -4.06026 | 4.90E-05 | 0.00214 | 136.5099 | 63.49765 | 150.9785 | 128.3385 |
| 286 | Fbxw7 | 1337.824 | 0.28494 | 0.07022 | -4.0577 | 4.96E-05 | 0.00216 | 1115.727 | 1373.603 | 1189.831 | 1672.137 |
| 287 | Ifnlr1 | 31.17607 | -1.15458 | 0.28524 | 4.0477 | 5.17E-05 | 0.00225 | 32.03815 | 52.86742 | 16.26849 | 23.53024 |
| 288 | Npr2 | 27.41786 | 1.77601 | 0.43917 | -4.04398 | 5.26E-05 | 0.00227 | 17.4464 | 8.733703 | 49.05871 | 34.43263 |
| 289 | Serpina3g | 1709.411 | 0.72011 | 0.17833 | -4.03805 | 5.39E-05 | 0.00232 | 2047.089 | 866.307 | 2487.857 | 1436.39 |

| | | | | | | | | | | | |
|---|---|---|---|---|---|---|---|---|---|---|---|
| 290 | Rgs13 | 69.21763 | -1.08465 | 0.26881 | 4.03509 | 5.46E-05 | 0.00235 | 88.81836 | 102.6339 | 37.88504 | 47.53325 |
| 291 | Orc5 | 349.0271 | 0.59176 | 0.14674 | -4.03266 | 5.51E-05 | 0.00236 | 339.6973 | 270.1311 | 379.1863 | 407.0937 |
| 292 | Rbpms | 113.7187 | 0.83628 | 0.20756 | -4.02909 | 5.60E-05 | 0.00239 | 114.9035 | 34.63995 | 242.033 | 63.29831 |
| 293 | Pced1b | 362.3042 | 0.46475 | 0.1154 | -4.02713 | 5.65E-05 | 0.0024 | 409.3844 | 240.3774 | 466.488 | 332.9669 |
| 294 | Ccdc23 | 254.6775 | 0.89271 | 0.22177 | -4.02546 | 5.69E-05 | 0.00241 | 232.3984 | 181.0702 | 265.7022 | 339.5393 |
| 295 | Casc5 | 120.8898 | -1.08332 | 0.26934 | 4.02209 | 5.77E-05 | 0.00244 | 213.9769 | 109.8242 | 109.1921 | 50.56598 |
| 296 | Abhd17b | 1635.484 | 0.42752 | 0.10647 | -4.01552 | 5.93E-05 | 0.0025 | 2205.463 | 712.0029 | 2665.948 | 958.5234 |
| 297 | Mybl1 | 36.11303 | -1.49688 | 0.37361 | 4.00654 | 6.16E-05 | 0.00259 | 62.26257 | 43.74806 | 24.03878 | 14.40272 |
| 298 | Kif11 | 178.6085 | -0.69524 | 0.17392 | 3.99758 | 6.40E-05 | 0.00266 | 255.5126 | 201.8629 | 133.073 | 123.9855 |
| 299 | Pdlim7 | 262.4579 | 0.72686 | 0.18185 | -3.99703 | 6.41E-05 | 0.00266 | 408.4941 | 91.13732 | 397.8421 | 152.3582 |
| 300 | S1pr1 | 5371.4 | 0.4585 | 0.11471 | -3.99694 | 6.42E-05 | 0.00266 | 7925.479 | 1733.829 | 9438.884 | 2387.41 |
| 301 | Bhlhe40 | 3559.502 | -0.38442 | 0.09635 | 3.99002 | 6.61E-05 | 0.00273 | 4466.413 | 3317.437 | 3913.206 | 2540.953 |
| 302 | Trim25 | 3468.975 | 0.60524 | 0.15175 | -3.98834 | 6.65E-05 | 0.00274 | 4878.812 | 1452.223 | 5328.753 | 2216.112 |
| 303 | Pik3cd | 4677.931 | -0.43716 | 0.10962 | 3.98813 | 6.66E-05 | 0.00274 | 6703.007 | 3574.275 | 5796.42 | 2638.021 |
| 304 | Ncf1 | 3485.571 | -0.40547 | 0.10171 | 3.9867 | 6.70E-05 | 0.00274 | 3973.166 | 4001.574 | 2948.412 | 3019.13 |
| 305 | Faim3 | 11858.32 | 0.4805 | 0.12128 | -3.96175 | 7.44E-05 | 0.00304 | 14698.29 | 6074.942 | 18168.08 | 8491.974 |
| 306 | Tm6sf1 | 918.3987 | 0.8897 | 0.22492 | -3.95556 | 7.64E-05 | 0.00311 | 1761.758 | 93.21768 | 1642.423 | 176.1955 |
| 307 | Gm14446 | 73.74643 | 1.59093 | 0.40328 | -3.94502 | 7.98E-05 | 0.00324 | 110.7205 | 4.86053 | 162.179 | 17.2257 |
| 309 | Smap2 | 5132.538 | 0.23034 | 0.05854 | -3.93504 | 8.32E-05 | 0.00336 | 7331.804 | 2575.188 | 7598.85 | 3024.31 |
| 310 | Hepacam2 | 150.3691 | 1.47619 | 0.37544 | -3.93194 | 8.43E-05 | 0.0034 | 216.8338 | 5.586687 | 361.1238 | 17.93228 |
| 311 | Cd48 | 1182.885 | 0.38524 | 0.09806 | -3.92873 | 8.54E-05 | 0.00342 | 1603.036 | 570.839 | 1808.928 | 748.7389 |
| 312 | Sertad1 | 357.2175 | 0.74622 | 0.18996 | -3.92818 | 8.56E-05 | 0.00342 | 283.3804 | 290.2061 | 364.8933 | 490.3904 |
| 313 | Pdcd1lg2 | 209.8793 | -0.57154 | 0.1456 | 3.92542 | 8.66E-05 | 0.00345 | 170.4108 | 362.0827 | 64.35483 | 242.669 |
| 314 | Parp9 | 1054.555 | 0.40621 | 0.10352 | -3.92388 | 8.71E-05 | 0.00346 | 1159.833 | 771.6869 | 1264.416 | 1022.283 |
| 315 | Ralgds | 1090.608 | -0.38635 | 0.09851 | 3.92208 | 8.78E-05 | 0.00347 | 1816.546 | 573.534 | 1532.496 | 439.8574 |
| 316 | 2810417H1: | 168.9216 | -0.73816 | 0.18833 | 3.91959 | 8.87E-05 | 0.0035 | 313.643 | 108.7704 | 188.5532 | 64.71993 |
| 317 | Bank1 | 7289.674 | 0.42842 | 0.1094 | -3.91615 | 9.00E-05 | 0.00353 | 10979.64 | 2797.414 | 11614.99 | 3766.65 |
| 318 | Tyk2 | 1166.868 | -0.38349 | 0.09793 | 3.916 | 9.00E-05 | 0.00353 | 1720.905 | 857.7009 | 1431.899 | 656.9678 |
| 319 | Ap1s3 | 922.9543 | 0.36749 | 0.09391 | -3.91327 | 9.11E-05 | 0.00356 | 1116.737 | 713.4791 | 941.2047 | 920.3966 |
| 320 | Tnik | 179.4568 | -0.57853 | 0.14792 | 3.9112 | 9.18E-05 | 0.00357 | 220.9201 | 213.1267 | 142.5023 | 141.2783 |
| 321 | Ppcdc | 1148.796 | 0.4363 | 0.11157 | -3.91068 | 9.20E-05 | 0.00357 | 1503.009 | 620.0257 | 1631.213 | 840.9383 |
| 322 | Iqgap1 | 18150.2 | -0.59395 | 0.1519 | 3.91024 | 9.22E-05 | 0.00357 | 17512.04 | 24169.97 | 14956.38 | 15962.41 |
| 323 | Lrig1 | 62.78883 | 1.11825 | 0.28687 | -3.89806 | 9.70E-05 | 0.00375 | 21.97156 | 19.18423 | 167.0451 | 42.95444 |
| 324 | Mid1 | 612.1837 | 1.09981 | 0.2827 | -3.89038 | 0.0001 | 0.00386 | 325.8053 | 468.2186 | 629.188 | 1025.523 |
| 325 | Tnfaip8 | 1171.646 | 0.52073 | 0.134 | -3.88612 | 0.0001 | 0.0039 | 1535.017 | 666.4474 | 1527.641 | 957.4774 |
| 326 | Slc27a1 | 149.0839 | -0.64954 | 0.16715 | 3.88601 | 0.0001 | 0.0039 | 214.3518 | 127.4255 | 173.2697 | 81.28853 |
| 327 | Mndal | 3956.413 | 0.49667 | 0.12788 | -3.88394 | 0.0001 | 0.00392 | 6609.676 | 1080.396 | 6607.973 | 1527.607 |
| 328 | Fer | 31.37153 | 1.8692 | 0.48184 | -3.87928 | 0.0001 | 0.00398 | 20.88903 | 3.272835 | 86.74451 | 14.57974 |
| 329 | Tpst2 | 1890.706 | 0.8146 | 0.21033 | -3.87286 | 0.00011 | 0.00406 | 644.8272 | 2194.549 | 831.6804 | 3891.769 |
| 330 | 3300002I08 | 20.71766 | -1.63816 | 0.42302 | 3.87255 | 0.00011 | 0.00406 | 34.02226 | 25.94833 | 15.38788 | 7.512177 |
| 331 | Stat5b | 2434.582 | -0.31433 | 0.0812 | 3.87117 | 0.00011 | 0.00408 | 2070.95 | 3096.174 | 2082.497 | 2488.708 |
| 332 | Jarid2 | 3014.53 | 0.36742 | 0.09514 | -3.86185 | 0.00011 | 0.00422 | 3276.265 | 2565.667 | 2904.412 | 3311.777 |
| 333 | Phlda1 | 67.32452 | 1.08736 | 0.28183 | -3.85821 | 0.00011 | 0.00426 | 75.73447 | 27.8703 | 106.0543 | 59.63899 |
| 334 | Sla | 2539.641 | 0.73683 | 0.19106 | -3.85663 | 0.00011 | 0.00427 | 4525.58 | 239.3477 | 4991.135 | 402.5017 |
| 335 | Clcn7 | 1194.793 | -0.29832 | 0.07737 | 3.85606 | 0.00012 | 0.00427 | 908.4163 | 1659.251 | 862.6366 | 1348.868 |
| 336 | Gcnt1 | 511.9043 | -0.61389 | 0.15933 | 3.85286 | 0.00012 | 0.00432 | 726.6931 | 460.6442 | 561.3289 | 298.9508 |
| 337 | Rab11fip4 | 104.5889 | 0.95824 | 0.24886 | -3.85049 | 0.00012 | 0.00435 | 77.09796 | 76.75313 | 113.1674 | 151.3372 |
| 338 | Gla | 1682.287 | 0.5505 | 0.14305 | -3.84829 | 0.00012 | 0.00437 | 2954.731 | 305.1481 | 3021.367 | 447.9018 |
| 339 | Mx1 | 359.0244 | 1.2647 | 0.32868 | -3.84783 | 0.00012 | 0.00437 | 472.6262 | 112.6603 | 570.9416 | 279.8696 |
| 340 | Dsn1 | 169.4503 | -0.56589 | 0.14727 | 3.84245 | 0.00012 | 0.00446 | 217.4207 | 185.312 | 150.9032 | 124.1651 |
| 341 | Ly6c1 | 42.4342 | 1.22618 | 0.31934 | -3.83969 | 0.00012 | 0.00449 | 48.5823 | 13.45068 | 75.87154 | 31.83227 |
| 342 | Igkc | 34145.39 | 0.57721 | 0.15041 | -3.83764 | 0.00012 | 0.00452 | 43382.34 | 13005.99 | 60729.61 | 19463.62 |
| 343 | Trim34a | 774.0071 | 0.80506 | 0.20989 | -3.8356 | 0.00013 | 0.00454 | 1200.057 | 172.6125 | 1419.782 | 303.5764 |
| 344 | Arid5a | 2514.84 | -0.33526 | 0.08745 | 3.83388 | 0.00013 | 0.00456 | 2486.618 | 2535.605 | 3027.918 | 2009.219 |
| 345 | Smim3 | 121.5806 | -0.7241 | 0.18895 | 3.8322 | 0.00013 | 0.00458 | 140.3761 | 153.1284 | 100.4313 | 92.38671 |
| 346 | Wdr81 | 1627.256 | -0.46375 | 0.12108 | 3.83004 | 0.00013 | 0.00461 | 1667.275 | 2007.924 | 1380.048 | 1453.777 |
| 347 | Cln8 | 100.1456 | 1.29923 | 0.33937 | -3.82842 | 0.00013 | 0.00463 | 121.2323 | 17.12827 | 217.0522 | 45.16964 |
| 348 | Arhgef10 | 67.00201 | 1.30094 | 0.34022 | -3.82379 | 0.00013 | 0.0047 | 121.3047 | 9.42331 | 112.9223 | 24.35778 |
| 349 | Zmiz2 | 6639.594 | -0.26762 | 0.06999 | 3.82342 | 0.00013 | 0.0047 | 5831.354 | 8586.184 | 5010.82 | 7130.017 |
| 350 | Apbb1ip | 1524.906 | 0.26668 | 0.0698 | -3.82068 | 0.00013 | 0.00473 | 2198.945 | 919.9587 | 1873.271 | 1107.448 |
| 351 | 3110043O2: | 367.719 | 0.4426 | 0.1159 | -3.81878 | 0.00013 | 0.00475 | 333.1235 | 323.9367 | 374.6388 | 439.177 |
| 353 | Gm15561 | 9.387069 | -2.03238 | 0.53225 | 3.8185 | 0.00013 | 0.00475 | 3.006904 | 25.98667 | 2.864498 | 5.690204 |
| 354 | St3gal6 | 568.0559 | 0.42787 | 0.11208 | -3.81748 | 0.00013 | 0.00476 | 828.016 | 269.2953 | 812.834 | 362.0784 |
| 355 | Atg16l2 | 865.8304 | -0.32777 | 0.08592 | 3.81471 | 0.00014 | 0.0048 | 1280.175 | 538.577 | 1215.707 | 428.8628 |
| 356 | Acp5 | 1165.867 | 0.79502 | 0.2085 | -3.81294 | 0.00014 | 0.00481 | 2205.94 | 113.6265 | 2144.108 | 199.7955 |
| 357 | Bst2 | 554.9526 | 0.6857 | 0.17984 | -3.8129 | 0.00014 | 0.00481 | 434.6805 | 460.27 | 580.7368 | 744.1232 |
| 358 | Gimap6 | 10880.44 | 0.43977 | 0.11542 | -3.81011 | 0.00014 | 0.00485 | 14562.87 | 5132.873 | 16854.25 | 6971.762 |
| 359 | 4632428N0! | 1148.2 | 0.48777 | 0.12818 | -3.80523 | 0.00014 | 0.00493 | 959.2258 | 1022.64 | 1174.325 | 1436.61 |
| 360 | B3gnt7 | 2381.571 | 0.81483 | 0.21421 | -3.80384 | 0.00014 | 0.00495 | 724.2836 | 2775.917 | 1101.384 | 4924.698 |
| 361 | Prr7 | 104.542 | 1.24832 | 0.32886 | -3.79594 | 0.00015 | 0.00509 | 147.9881 | 15.34096 | 216.8814 | 37.95756 |
| 362 | Gm15228 | 12.3667 | 2.85958 | 0.75435 | -3.79077 | 0.00015 | 0.00519 | 6.959687 | 0 | 35.26757 | 7.239522 |
| 363 | H60b | 42.84086 | 1.57744 | 0.41655 | -3.78695 | 0.00015 | 0.00525 | 51.99465 | 7.945936 | 86.29325 | 25.1296 |
| 364 | Atp6v1a | 1006.896 | -0.37261 | 0.09853 | 3.78164 | 0.00016 | 0.00535 | 1084.067 | 1102.882 | 989.3868 | 851.2488 |

| | | | | | | | | | | | |
|---|---|---|---|---|---|---|---|---|---|---|---|
| 365 | Hook1 | 874.9783 | -0.42666 | 0.11307 | 3.77351 | 0.00016 | 0.00552 | 764.2485 | 1278.909 | 507.4641 | 949.2919 |
| 366 | Slc35g1 | 81.21143 | 1.08537 | 0.28789 | -3.77009 | 0.00016 | 0.00558 | 22.66047 | 82.26936 | 42.05837 | 177.8575 |
| 367 | Plod2 | 27.32701 | -1.15422 | 0.30654 | 3.76536 | 0.00017 | 0.00567 | 2.980649 | 71.58448 | 2.891733 | 31.8512 |
| 368 | Gm24270 | 71.88717 | 2.89227 | 0.76876 | -3.76227 | 0.00017 | 0.00572 | 12.46084 | 16.59547 | 27.07381 | 231.4186 |
| 369 | Wdhd1 | 532.3956 | -0.3848 | 0.10236 | 3.75937 | 0.00017 | 0.00578 | 572.7969 | 601.8363 | 495.1426 | 459.8065 |
| 370 | Dhx40 | 925.2769 | 0.3279 | 0.08741 | -3.75143 | 0.00018 | 0.00595 | 1383.736 | 469.5544 | 1256.61 | 591.2075 |
| 371 | Chchd10 | 405.6794 | 0.81099 | 0.21714 | -3.73494 | 0.00019 | 0.00633 | 417.4341 | 246.9372 | 521.6732 | 436.6732 |
| 372 | Ubald2 | 1189.773 | 0.51816 | 0.13884 | -3.73196 | 0.00019 | 0.00639 | 1264.04 | 784.6586 | 1583.113 | 1127.282 |
| 373 | Pla2g7 | 184.8272 | 1.49798 | 0.40171 | -3.72901 | 0.00019 | 0.00645 | 303.7442 | 14.14362 | 377.5196 | 43.90123 |
| 374 | Ifi35 | 947.9968 | 0.54468 | 0.14626 | -3.72396 | 0.0002 | 0.00656 | 645.9652 | 937.1625 | 837.8796 | 1370.98 |
| 375 | Ddit4 | 584.3362 | 0.77898 | 0.20929 | -3.72201 | 0.0002 | 0.0066 | 401.0773 | 528.4757 | 494.3965 | 913.3953 |
| 376 | Tgif2 | 742.9866 | 0.43023 | 0.11564 | -3.72058 | 0.0002 | 0.00662 | 917.3005 | 464.8046 | 961.0827 | 628.7586 |
| 377 | Oas1c | 195.2274 | 0.73046 | 0.19646 | -3.71821 | 0.0002 | 0.00666 | 321.8514 | 42.49066 | 344.3985 | 72.16909 |
| 378 | Map3k14 | 1284.673 | 0.24972 | 0.06717 | -3.71764 | 0.0002 | 0.00666 | 1455.795 | 1015.457 | 1457.695 | 1209.746 |
| 379 | Plbd1 | 667.6168 | 0.5297 | 0.14257 | -3.71538 | 0.0002 | 0.00671 | 901.2758 | 294.1065 | 1049.896 | 425.1889 |
| 380 | Cetn4 | 26.14246 | 1.31481 | 0.35505 | -3.70313 | 0.00021 | 0.00702 | 2.867311 | 26.4789 | 6.906256 | 68.31739 |
| 381 | Pisd-ps2 | 286.5423 | -0.79063 | 0.21372 | 3.69943 | 0.00022 | 0.00711 | 515.6233 | 211.6079 | 298.2411 | 120.6968 |
| 382 | Zfp219 | 269.0837 | 0.53723 | 0.14525 | -3.69861 | 0.00022 | 0.00711 | 179.6454 | 275.8762 | 220.8496 | 399.9636 |
| 383 | Igkv4-63 | 68.03531 | 1.06699 | 0.28877 | -3.69499 | 0.00022 | 0.00719 | 90.07824 | 15.33543 | 133.24 | 33.48752 |
| 384 | Agpat4 | 209.1676 | 0.7329 | 0.19843 | -3.69358 | 0.00022 | 0.00722 | 162.4364 | 190.9191 | 163.6367 | 319.6781 |
| 385 | Cacna1s | 34.80759 | 1.88461 | 0.51204 | -3.68059 | 0.00023 | 0.00757 | 53.52037 | 3.48351 | 66.45353 | 15.77296 |
| 386 | Gm23935 | 766.6136 | 2.74707 | 0.74644 | -3.68025 | 0.00023 | 0.00757 | 150.0514 | 210.0174 | 297.0911 | 2409.294 |
| 387 | Ryk | 31.68886 | 1.15544 | 0.31404 | -3.67932 | 0.00023 | 0.00757 | 15.88468 | 22.25483 | 37.61204 | 51.00391 |
| 388 | Tap1 | 3979.396 | 0.34453 | 0.09386 | -3.67071 | 0.00024 | 0.00782 | 4500.607 | 2899.827 | 4830.905 | 3686.244 |
| 389 | Pik3ap1 | 9426.474 | 0.23125 | 0.06303 | -3.66883 | 0.00024 | 0.00785 | 7932.95 | 10085.38 | 7849.284 | 11838.28 |
| 390 | Sema4b | 2802.762 | 0.26733 | 0.07293 | -3.66534 | 0.00025 | 0.00794 | 2537.871 | 2758.267 | 2591.955 | 3322.955 |
| 391 | Ogfrl1 | 358.3871 | 0.54333 | 0.14864 | -3.65529 | 0.00026 | 0.00823 | 440.3207 | 196.4145 | 510.2426 | 286.5706 |
| 392 | Phf11d | 89.41974 | 0.92427 | 0.25287 | -3.65507 | 0.00026 | 0.00823 | 116.1151 | 25.64243 | 165.7682 | 50.15327 |
| 393 | Prr5 | 780.0408 | 0.55476 | 0.15229 | -3.64281 | 0.00027 | 0.00859 | 1153.309 | 289.8097 | 1248.358 | 428.6857 |
| 394 | Trim35 | 1165.24 | 0.32116 | 0.08826 | -3.63871 | 0.00027 | 0.0087 | 1609.284 | 549.3422 | 1815.75 | 686.5828 |
| 395 | Ccr9 | 101.3951 | -0.71553 | 0.19673 | 3.63718 | 0.00028 | 0.00873 | 36.39092 | 216.645 | 20.80239 | 131.742 |
| 396 | Rsad2 | 202.0045 | 1.31909 | 0.36282 | -3.63568 | 0.00028 | 0.00876 | 241.1232 | 70.50106 | 312.017 | 184.3767 |
| 397 | Btg2 | 4951.865 | 0.51836 | 0.14272 | -3.63209 | 0.00028 | 0.00886 | 8325.684 | 1023.227 | 8989.09 | 1469.46 |
| 398 | Galc | 102.8294 | 0.69007 | 0.19012 | -3.62961 | 0.00028 | 0.00889 | 68.75826 | 84.99057 | 120.5746 | 136.9942 |
| 399 | Trafd1 | 870.8615 | 0.53204 | 0.14658 | -3.62957 | 0.00028 | 0.00889 | 1276.966 | 347.8828 | 1354.484 | 504.114 |
| 400 | Pttg1 | 391.8666 | 0.70491 | 0.19423 | -3.62918 | 0.00028 | 0.00889 | 381.6311 | 263.1386 | 490.499 | 432.1977 |
| 401 | Folr4 | 24.04767 | 1.80253 | 0.49673 | -3.62877 | 0.00028 | 0.00889 | 19.63998 | 3.002362 | 59.81507 | 13.73327 |
| 402 | Ankrd12 | 2435.516 | -0.79809 | 0.2201 | 3.626 | 0.00029 | 0.00897 | 1949.854 | 3860.392 | 1732.093 | 2199.724 |
| 403 | Tusc1 | 22.25976 | 1.43136 | 0.39556 | -3.61853 | 0.0003 | 0.00921 | 12.93751 | 11.01322 | 32.97399 | 32.11433 |
| 404 | Adam10 | 3179.921 | -0.22359 | 0.0618 | 3.61781 | 0.0003 | 0.00921 | 2677.994 | 3937.416 | 2734.952 | 3369.322 |
| 405 | Hfe | 140.2159 | 1.14282 | 0.31596 | -3.61694 | 0.0003 | 0.00922 | 201.0155 | 36.61397 | 239.8246 | 83.40944 |
| 406 | Cmtm7 | 1129.881 | 0.57525 | 0.15914 | -3.61481 | 0.0003 | 0.00927 | 1194.532 | 834.293 | 1242.414 | 1248.284 |
| 407 | Bsn | 83.46358 | -1.11338 | 0.30929 | 3.59975 | 0.00032 | 0.0098 | 51.35146 | 178.2547 | 23.72429 | 80.52383 |
| 408 | Etv5 | 187.6708 | 0.67363 | 0.1873 | -3.59644 | 0.00032 | 0.00989 | 253.212 | 76.94208 | 296.331 | 124.1981 |
| 409 | Sidt1 | 261.5674 | 0.73023 | 0.20305 | -3.59629 | 0.00032 | 0.00989 | 411.2018 | 56.07602 | 484.6994 | 94.2923 |
| 410 | Prr11 | 42.36655 | -1.18399 | 0.33002 | 3.58767 | 0.00033 | 0.0102 | 70.73079 | 34.53738 | 49.63215 | 14.56588 |
| 411 | Hrsp12 | 156.0755 | -0.74383 | 0.20752 | 3.58433 | 0.00034 | 0.01031 | 192.5955 | 168.1068 | 163.8368 | 99.76299 |
| 412 | Car2 | 181.2809 | 0.8162 | 0.2278 | -3.58291 | 0.00034 | 0.01034 | 232.0029 | 79.27546 | 271.4129 | 142.4322 |
| 413 | Cyth1 | 3616.967 | -0.37088 | 0.10354 | 3.58205 | 0.00034 | 0.01035 | 5111.713 | 2471.262 | 4976.516 | 1908.377 |
| 414 | Ank | 409.6267 | -0.33461 | 0.09344 | 3.58092 | 0.00034 | 0.01037 | 276.7901 | 639.3827 | 216.5468 | 505.7872 |
| 416 | Cep128 | 222.0309 | -0.58184 | 0.16306 | 3.56819 | 0.00036 | 0.01086 | 348.8213 | 125.4386 | 330.3694 | 83.49416 |
| 417 | Ppp1r15a | 7931.558 | 0.45735 | 0.1282 | -3.5676 | 0.00036 | 0.01086 | 12496.38 | 2656.948 | 12918.28 | 3654.624 |
| 418 | Igkv15-103 | 628.4502 | 1.45598 | 0.41003 | -3.55093 | 0.00038 | 0.01154 | 500.1441 | 197.734 | 1237.621 | 578.3018 |
| 419 | Scaper | 513.0516 | -0.42803 | 0.12065 | 3.54777 | 0.00039 | 0.01166 | 600.3866 | 528.6058 | 531.0666 | 392.1473 |
| 420 | Stat3 | 7883.128 | -0.40481 | 0.1142 | 3.54465 | 0.00039 | 0.01177 | 3113.095 | 14362.77 | 3222.066 | 10834.58 |
| 421 | Itpkb | 2866.119 | 0.42342 | 0.11972 | -3.53681 | 0.0004 | 0.01209 | 3495.269 | 2008.062 | 3265.346 | 2695.799 |
| 422 | Lta | 214.5966 | 0.7007 | 0.19838 | -3.53212 | 0.00041 | 0.01228 | 356.275 | 49.24035 | 372.4733 | 80.39763 |
| 423 | Sfn | 931.3321 | 0.68671 | 0.19455 | -3.52968 | 0.00042 | 0.01237 | 854.1652 | 692.5195 | 1057.597 | 1121.047 |
| 424 | Fosb | 18101.76 | 2.49742 | 0.70857 | -3.52459 | 0.00042 | 0.01258 | 35028.2 | 46.48078 | 36590.06 | 742.3037 |
| 425 | Agpat3 | 1678.116 | 0.26416 | 0.07499 | -3.5225 | 0.00043 | 0.01265 | 1765.883 | 1381.356 | 1902.866 | 1662.359 |
| 426 | Rps27a-ps2 | 69.04074 | 1.41901 | 0.40317 | -3.51962 | 0.00043 | 0.01276 | 41.74954 | 39.62699 | 82.395 | 112.3914 |
| 427 | Gm2a | 1741.838 | -0.41348 | 0.11758 | 3.51666 | 0.00044 | 0.01285 | 3373.537 | 523.5177 | 2677.063 | 393.2363 |
| 428 | Hvcn1 | 13502.69 | -0.25928 | 0.07373 | 3.51651 | 0.00044 | 0.01285 | 11962.46 | 16077.11 | 12541.26 | 13429.92 |
| 430 | Nod1 | 324.6139 | 1.37126 | 0.39027 | -3.51357 | 0.00044 | 0.01297 | 658.5712 | 12.76329 | 590.1874 | 36.93381 |
| 431 | Fry | 838.8 | -0.8969 | 0.25532 | 3.51291 | 0.00044 | 0.01297 | 1555.254 | 350.1035 | 1264.807 | 185.0349 |
| 432 | Npepps | 1321.754 | -0.22051 | 0.0628 | 3.51134 | 0.00045 | 0.01302 | 1106.334 | 1650.573 | 1114.316 | 1415.794 |
| 433 | Dgkd | 9452.327 | -0.3105 | 0.08852 | 3.50761 | 0.00045 | 0.01317 | 13099.92 | 7371.088 | 11398.41 | 5939.889 |
| 434 | Arhgap11a | 505.4558 | -0.43411 | 0.12381 | 3.50634 | 0.00045 | 0.0132 | 743.0433 | 393.6229 | 594.0991 | 291.058 |
| 435 | 03-Mar | 160.8991 | 0.60022 | 0.17132 | -3.50362 | 0.00046 | 0.01331 | 90.9169 | 166.5909 | 133.6125 | 252.4761 |
| 436 | Ryr2 | 58.08955 | -1.23519 | 0.35279 | 3.50119 | 0.00046 | 0.01339 | 122.6313 | 43.17404 | 48.96301 | 17.58987 |
| 437 | Ddx58 | 1749.601 | 0.66149 | 0.18895 | -3.50084 | 0.00046 | 0.01339 | 2647.696 | 563.3788 | 2890.795 | 896.5327 |
| 438 | Tjp1 | 19.76812 | 1.49041 | 0.42583 | -3.49999 | 0.00047 | 0.0134 | 18.83798 | 6.606159 | 33.50636 | 20.12198 |
| 439 | Slc22a15 | 25.88676 | -1.5203 | 0.43453 | 3.49872 | 0.00047 | 0.01344 | 51.2123 | 31.81064 | 10.08524 | 10.43886 |

| | | | | | | | | | | | |
|---|---|---|---|---|---|---|---|---|---|---|---|
| 440 | Nfatc1 | 7465.686 | -0.39056 | 0.11166 | 3.49762 | 0.00047 | 0.01346 | 6089.523 | 10683.59 | 4949.241 | 8140.392 |
| 441 | Fuca1 | 1429.071 | 0.35373 | 0.10124 | -3.4941 | 0.00048 | 0.01359 | 1899.711 | 642.3625 | 2350.771 | 823.4396 |
| 442 | Zfp260 | 431.3488 | -0.32065 | 0.09177 | 3.49392 | 0.00048 | 0.01359 | 429.3364 | 511.2329 | 376.749 | 408.0771 |
| 443 | Ppp3ca | 3581.2 | 0.45829 | 0.13131 | -3.4903 | 0.00048 | 0.01375 | 3960.298 | 2626.302 | 4124.212 | 3613.99 |
| 444 | Mfng | 1106.655 | 0.40853 | 0.11707 | -3.48961 | 0.00048 | 0.01375 | 976.3267 | 1011.126 | 1094.536 | 1344.633 |
| 445 | Zap70 | 28.55081 | 1.78493 | 0.51178 | -3.48769 | 0.00049 | 0.01381 | 23.63688 | 2.083244 | 78.07331 | 10.4098 |
| 446 | Tuba1a | 1442.909 | 0.6008 | 0.17228 | -3.48728 | 0.00049 | 0.01381 | 2115.583 | 411.1912 | 2618.089 | 626.7715 |
| 447 | Rasd1 | 302.0288 | 0.6709 | 0.19249 | -3.48533 | 0.00049 | 0.01386 | 362.44 | 168.662 | 407.29 | 269.7231 |
| 448 | Gm9864 | 98.90372 | -1.09915 | 0.31538 | 3.48516 | 0.00049 | 0.01386 | 108.5314 | 146.951 | 73.39444 | 66.73797 |
| 449 | Sp100 | 6688.018 | 0.3147 | 0.09043 | -3.47995 | 0.0005 | 0.0141 | 9206.115 | 3259.015 | 10231.62 | 4055.318 |
| 450 | Kifap3 | 38.3617 | -1.04547 | 0.30058 | 3.47818 | 0.0005 | 0.01417 | 51.84869 | 41.53208 | 40.58967 | 19.47636 |
| 451 | Cdc25a | 446.6485 | 0.40165 | 0.1155 | -3.47757 | 0.00051 | 0.01417 | 233.4148 | 540.1773 | 298.8969 | 714.1048 |
| 452 | Hyi | 179.0205 | 0.70146 | 0.20188 | -3.47467 | 0.00051 | 0.01429 | 110.9982 | 163.5299 | 174.1873 | 267.3667 |
| 454 | Pml | 5196.568 | 0.5734 | 0.16518 | -3.47128 | 0.00052 | 0.01444 | 7716.579 | 2069.069 | 7909.814 | 3090.812 |
| 455 | Fam101b | 333.6741 | 0.87263 | 0.25146 | -3.47024 | 0.00052 | 0.01447 | 468.3459 | 22.16445 | 802.2004 | 41.9855 |
| 456 | Crb3 | 96.6743 | 0.74647 | 0.21552 | -3.46353 | 0.00053 | 0.01477 | 87.22812 | 73.75296 | 102.1452 | 123.5709 |
| 457 | Rps4l | 100.0852 | 0.89223 | 0.25762 | -3.46335 | 0.00053 | 0.01477 | 69.60842 | 76.9742 | 108.0548 | 145.7033 |
| 458 | Mid1ip1 | 298.2185 | 0.47473 | 0.13709 | -3.46301 | 0.00053 | 0.01477 | 298.3511 | 243.9845 | 311.598 | 338.9404 |
| 459 | Ell2 | 843.2463 | 0.48778 | 0.14098 | -3.45981 | 0.00054 | 0.01491 | 678.3626 | 765.3271 | 855.2666 | 1074.029 |
| 460 | Tusc3 | 358.7181 | 0.46389 | 0.13424 | -3.45571 | 0.00055 | 0.01511 | 343.3565 | 274.2894 | 437.9684 | 379.2581 |
| 461 | Gcsam | 14.47406 | -2.48121 | 0.71911 | 3.45037 | 0.00056 | 0.01537 | 27.50777 | 21.52575 | 6.760026 | 2.102701 |
| 462 | Arhgef3 | 2300.158 | -0.40309 | 0.11684 | 3.45002 | 0.00056 | 0.01537 | 3750.983 | 1323.158 | 3127.202 | 999.2885 |
| 463 | Id2 | 693.4425 | 0.73014 | 0.21172 | -3.44856 | 0.00056 | 0.01542 | 759.2266 | 400.2948 | 945.0509 | 669.1976 |
| 464 | Abhd6 | 141.8256 | -0.56434 | 0.1637 | 3.44747 | 0.00057 | 0.01545 | 133.4211 | 190.6021 | 115.0819 | 128.1974 |
| 465 | Cystm1 | 166.9591 | 0.95544 | 0.27745 | -3.44364 | 0.00057 | 0.01563 | 53.98052 | 181.3599 | 75.42058 | 357.0754 |
| 466 | Hs3st2 | 31.29354 | 1.50953 | 0.43841 | -3.44317 | 0.00057 | 0.01563 | 37.25223 | 9.543597 | 49.01784 | 29.3605 |
| 467 | Neurl3 | 1956.358 | -0.57396 | 0.16676 | 3.44187 | 0.00058 | 0.01567 | 3578.195 | 449.9898 | 3496.273 | 300.9744 |
| 468 | Hecw2 | 20.72575 | -1.19841 | 0.34831 | 3.44063 | 0.00058 | 0.01571 | 20.45985 | 28.94072 | 21.41712 | 12.08531 |
| 469 | Ddx55 | 835.3897 | 0.28242 | 0.08222 | -3.43515 | 0.00059 | 0.016 | 661.035 | 906.7718 | 670.2359 | 1103.516 |
| 470 | Ddx25 | 63.56512 | 1.53904 | 0.44826 | -3.43338 | 0.0006 | 0.01607 | 88.22333 | 3.242936 | 151.1295 | 11.6647 |
| 471 | Rnf114 | 2317.653 | 0.26928 | 0.07847 | -3.43167 | 0.0006 | 0.01611 | 2268.304 | 1994.47 | 2601.104 | 2406.734 |
| 472 | Parp10 | 724.0022 | 0.28354 | 0.08263 | -3.43154 | 0.0006 | 0.01611 | 818.2132 | 537.6221 | 885.1697 | 655.0039 |
| 473 | Cdkn1a | 113.5937 | 0.58943 | 0.17185 | -3.42998 | 0.0006 | 0.01617 | 103.687 | 101.1078 | 97.40341 | 152.1766 |
| 474 | Ptprn | 15.85096 | 1.64614 | 0.47999 | -3.4295 | 0.0006 | 0.01617 | 12.13122 | 6.513554 | 22.40666 | 22.35239 |
| 475 | Zfp263 | 1745.967 | -0.19828 | 0.05793 | 3.42278 | 0.00062 | 0.01654 | 1326.878 | 2331.692 | 1294.752 | 2030.548 |
| 476 | Prodh | 106.9424 | 0.59535 | 0.17397 | -3.4221 | 0.00062 | 0.01655 | 50.11415 | 119.6766 | 76.23206 | 181.7469 |
| 477 | Il1rn | 44.66756 | 1.0394 | 0.30398 | -3.41924 | 0.00063 | 0.01669 | 2.697182 | 54.32195 | 7.541093 | 114.11 |
| 478 | Rhob | 3419.472 | 1.01588 | 0.29734 | -3.41658 | 0.00063 | 0.01682 | 5921.853 | 204.2313 | 7128.626 | 423.1761 |
| 479 | Igkv8-24 | 469.8718 | 0.67174 | 0.19669 | -3.41516 | 0.00064 | 0.01687 | 640.6959 | 145.1116 | 860.7012 | 232.9787 |
| 480 | Gm15638 | 10.32821 | -2.09467 | 0.61363 | 3.41355 | 0.00064 | 0.01694 | 13.06058 | 22.12055 | 2.133708 | 3.997989 |
| 481 | Phlpp1 | 2283.438 | -0.53602 | 0.15716 | 3.41067 | 0.00065 | 0.01708 | 673.5701 | 4567.589 | 752.695 | 3139.898 |
| 482 | Traf5 | 904.017 | 0.45812 | 0.1344 | -3.40874 | 0.00065 | 0.01717 | 1412.88 | 326.6876 | 1424.882 | 451.6182 |
| 483 | Rab20 | 106.1459 | -0.94152 | 0.27635 | 3.40698 | 0.00066 | 0.01724 | 63.06206 | 191.8289 | 71.28767 | 98.40517 |
| 484 | Crabp2 | 13.26234 | 1.399 | 0.41094 | -3.4044 | 0.00066 | 0.01737 | 3.04038 | 12.75332 | 1.454456 | 35.8012 |
| 485 | Ier5 | 4576.084 | 0.55913 | 0.16426 | -3.40396 | 0.00066 | 0.01737 | 6004.419 | 2169.341 | 6923.462 | 3207.116 |
| 486 | Camkk1 | 114.1713 | -0.68853 | 0.20238 | 3.4021 | 0.00067 | 0.01745 | 97.64647 | 162.7979 | 95.71096 | 100.5297 |
| 487 | Shisa5 | 15739.26 | 0.39834 | 0.11714 | -3.40072 | 0.00067 | 0.0175 | 27192.32 | 3736.108 | 27097.2 | 4931.394 |
| 488 | Rapgef5 | 349.2919 | -0.85022 | 0.2501 | 3.39953 | 0.00068 | 0.01754 | 394.8397 | 442.8475 | 317.2824 | 242.1979 |
| 489 | 6330403A0: | 38.88587 | 1.53812 | 0.4526 | -3.3984 | 0.00068 | 0.01758 | 24.47554 | 4.836162 | 111.1581 | 15.07367 |
| 491 | Fam105a | 240.8887 | -0.48511 | 0.14316 | 3.38865 | 0.0007 | 0.01818 | 302.353 | 220.1048 | 283.4688 | 157.6281 |
| 492 | Hcst | 112.7466 | 0.82585 | 0.24374 | -3.3882 | 0.0007 | 0.01818 | 154.9971 | 38.87246 | 186.7687 | 70.34828 |
| 493 | Reln | 71.66414 | -1.20389 | 0.35602 | 3.38154 | 0.00072 | 0.01859 | 136.8936 | 33.21865 | 102.695 | 13.84934 |
| 494 | Clec16a | 864.2326 | 0.30165 | 0.08927 | -3.37919 | 0.00073 | 0.01867 | 619.0094 | 898.2662 | 832.8653 | 1106.789 |
| 495 | Hhex | 873.1711 | 0.64965 | 0.19231 | -3.37816 | 0.00073 | 0.01868 | 1508.317 | 101.3812 | 1722.712 | 160.2743 |
| 496 | Ncapg2 | 196.2073 | -0.52672 | 0.15593 | 3.37795 | 0.00073 | 0.01868 | 223.8951 | 230.2926 | 171.6345 | 159.0069 |
| 497 | Sidt2 | 4343.871 | -0.25852 | 0.07655 | 3.37724 | 0.00073 | 0.01869 | 5254.393 | 4016.301 | 4748.754 | 3356.036 |
| 498 | Bmpr2 | 26.27525 | -1.19313 | 0.35346 | 3.37559 | 0.00074 | 0.01877 | 25.26488 | 44.88756 | 16.20037 | 18.74818 |
| 499 | Clint1 | 4136.157 | -0.18517 | 0.05492 | 3.37138 | 0.00075 | 0.01902 | 3060.548 | 5678.042 | 2812.904 | 4993.135 |
| 500 | Btg3 | 367.4972 | 0.59378 | 0.17637 | -3.36656 | 0.00076 | 0.01932 | 338.3163 | 311.0367 | 348.937 | 471.6986 |
| 501 | Mbnl1 | 6822.052 | -0.32663 | 0.0972 | 3.36026 | 0.00078 | 0.01973 | 9036.725 | 5624.02 | 8147.223 | 4480.238 |
| 503 | Chka | 1758.8 | 0.3942 | 0.11737 | -3.35872 | 0.00078 | 0.0198 | 1342.309 | 1815.752 | 1488.459 | 2388.682 |
| 504 | Ighg2c | 66.12465 | -2.58115 | 0.76896 | 3.35666 | 0.00079 | 0.01991 | 167.0284 | 79.78988 | 11.42657 | 6.253717 |
| 505 | Klhl9 | 289.6166 | -0.3639 | 0.10844 | 3.35581 | 0.00079 | 0.01993 | 209.0975 | 413.5425 | 215.0833 | 320.743 |
| 507 | Oosp1 | 15.79429 | 1.92037 | 0.57392 | -3.34605 | 0.00082 | 0.0206 | 18.6416 | 1.850419 | 32.17802 | 10.50711 |
| 508 | Mir132 | 13.53409 | 1.81394 | 0.54241 | -3.34424 | 0.00083 | 0.0207 | 7.176203 | 7.785784 | 8.139245 | 31.03512 |
| 509 | N4bp1 | 1415.459 | -0.3519 | 0.10532 | 3.34118 | 0.00083 | 0.02089 | 1041.985 | 2014.128 | 1029.79 | 1575.933 |
| 510 | Fam206a | 842.5819 | -0.40615 | 0.12162 | 3.33937 | 0.00084 | 0.02099 | 595.9824 | 1241.395 | 598.5387 | 934.4112 |
| 511 | Gcc1 | 564.8371 | -0.31295 | 0.09378 | 3.33725 | 0.00085 | 0.02108 | 513.0532 | 688.8997 | 504.0978 | 553.2979 |
| 512 | Tmed8 | 3441.774 | -0.45221 | 0.13551 | 3.33705 | 0.00085 | 0.02108 | 2291.162 | 5393.912 | 2147.759 | 3934.262 |
| 513 | Bin1 | 2236.458 | 0.38435 | 0.1152 | -3.33644 | 0.00085 | 0.02109 | 3997.139 | 688.0006 | 3360.75 | 899.9421 |
| 514 | Dclre1c | 1234.446 | -0.68404 | 0.20507 | 3.33562 | 0.00085 | 0.0211 | 1010.68 | 1942.898 | 783.5356 | 1200.672 |
| 515 | Ighg2b | 102.6915 | -1.76445 | 0.52905 | 3.33511 | 0.00085 | 0.0211 | 230.046 | 106.1186 | 47.69955 | 26.9017 |
| 516 | Spic | 297.249 | 1.27802 | 0.38326 | -3.33464 | 0.00085 | 0.0211 | 473.2988 | 29.06392 | 611.6721 | 74.96124 |

| | | | | | | | | | | | |
|---|---|---|---|---|---|---|---|---|---|---|---|
| 517 | Itgb3 | 1176.508 | -0.64966 | 0.19497 | 3.33214 | 0.00086 | 0.02125 | 1005.61 | 1585.549 | 1110.746 | 1004.126 |
| 518 | Slamf1 | 545.0378 | 0.61155 | 0.18371 | -3.32884 | 0.00087 | 0.02147 | 531.3472 | 439.1079 | 537.0091 | 672.6871 |
| 519 | Slc30a4 | 1020.279 | -0.38997 | 0.11725 | 3.32591 | 0.00088 | 0.02165 | 917.6943 | 1315.012 | 847.2642 | 1001.143 |
| 520 | Strbp | 2421.475 | 0.23188 | 0.06976 | -3.32402 | 0.00089 | 0.02176 | 3224.882 | 1390.265 | 3437.319 | 1633.435 |
| 521 | Acox3 | 1517.799 | -0.3957 | 0.11908 | 3.32296 | 0.00089 | 0.0218 | 1954.505 | 1286.515 | 1853.062 | 977.1134 |
| 522 | Xxylt1 | 557.363 | -0.34156 | 0.10282 | 3.3218 | 0.00089 | 0.02185 | 502.7103 | 774.4297 | 341.9827 | 610.3292 |
| 523 | Hap1 | 52.3676 | 1.4562 | 0.4387 | -3.31936 | 0.0009 | 0.022 | 79.52391 | 5.566755 | 106.8152 | 17.56454 |
| 524 | Vim | 3202.972 | -0.4138 | 0.12473 | 3.31739 | 0.00091 | 0.02211 | 4917.915 | 1982.242 | 4425.845 | 1485.886 |
| 525 | Dtx4 | 198.5207 | -0.51622 | 0.15572 | 3.31509 | 0.00092 | 0.02226 | 166.5909 | 261.9039 | 183.0949 | 182.4931 |
| 526 | Tnf | 160.1805 | 0.77289 | 0.23324 | -3.31368 | 0.00092 | 0.02233 | 64.79684 | 175.2471 | 98.65752 | 302.0204 |
| 527 | Pon3 | 42.97425 | -1.16764 | 0.35278 | 3.30987 | 0.00093 | 0.02259 | 53.06494 | 42.67558 | 57.89565 | 18.26082 |
| 528 | Slc12a7 | 327.6665 | -0.39742 | 0.12013 | 3.30809 | 0.00094 | 0.02269 | 348.9551 | 356.5559 | 334.2646 | 270.8903 |
| 529 | Hic1 | 130.2767 | 0.75138 | 0.22728 | -3.30596 | 0.00095 | 0.02282 | 60.85085 | 142.1388 | 77.6752 | 240.4418 |
| 530 | Uhrf1 | 441.6401 | -0.39673 | 0.12029 | 3.29812 | 0.00097 | 0.02343 | 482.289 | 531.1431 | 349.3072 | 403.8213 |
| 531 | Rinl | 270.251 | 0.54944 | 0.16675 | -3.29499 | 0.00098 | 0.02365 | 182.2644 | 259.4435 | 259.0971 | 380.1989 |
| 532 | Ifitm2 | 90.64268 | 0.99539 | 0.30233 | -3.29243 | 0.00099 | 0.02382 | 56.00214 | 72.49737 | 85.78721 | 148.284 |
| 533 | Ggh | 117.9281 | 0.69635 | 0.21166 | -3.29002 | 0.001 | 0.02398 | 107.8853 | 73.91604 | 168.6509 | 121.2603 |
| 534 | Hopxos | 9.099027 | 2.25813 | 0.68704 | -3.28674 | 0.00101 | 0.02422 | 9.886728 | 0.686292 | 19.27628 | 6.546804 |
| 535 | Gm14085 | 52.58333 | 2.14479 | 0.65271 | -3.28599 | 0.00102 | 0.02424 | 38.89944 | 2.205749 | 151.6317 | 17.59644 |
| 536 | Zxdb | 225.7707 | -0.37435 | 0.11397 | 3.28465 | 0.00102 | 0.02431 | 218.6086 | 276.3992 | 195.6118 | 212.4633 |
| 537 | Acot7 | 181.15 | 0.60262 | 0.18352 | -3.28375 | 0.00102 | 0.02434 | 97.56075 | 189.0498 | 148.9927 | 288.9966 |
| 538 | Ncoa7 | 289.3927 | -0.58261 | 0.17755 | 3.28142 | 0.00103 | 0.02446 | 316.5984 | 317.8888 | 312.0785 | 211.0049 |
| 539 | Spag9 | 4881.798 | -0.49298 | 0.15024 | 3.28131 | 0.00103 | 0.02446 | 4029.641 | 7415.745 | 2826.822 | 5254.985 |
| 540 | Ccrl2 | 46.79938 | 1.05073 | 0.32079 | -3.27542 | 0.00106 | 0.02489 | 65.43551 | 10.68004 | 87.4799 | 23.60205 |
| 541 | Smpdl3a | 352.6457 | 0.59883 | 0.18294 | -3.27334 | 0.00106 | 0.02501 | 594.3513 | 53.01164 | 682.1171 | 81.10288 |
| 542 | Cd22 | 11926.93 | -0.51567 | 0.15757 | 3.27261 | 0.00107 | 0.02501 | 22842.26 | 3689.668 | 18602.47 | 2573.322 |
| 543 | Cyb561a3 | 2395.602 | -0.33874 | 0.10351 | 3.27252 | 0.00107 | 0.02501 | 4173.849 | 883.895 | 3825.016 | 699.6463 |
| 544 | Cenpk | 34.78753 | -1.15235 | 0.35224 | 3.27146 | 0.00107 | 0.02506 | 56.47386 | 26.46119 | 44.76658 | 11.44849 |
| 545 | Tmem135 | 644.2227 | 0.31917 | 0.09762 | -3.26956 | 0.00108 | 0.02518 | 662.1734 | 518.6662 | 748.425 | 647.6261 |
| 546 | 5330426P16 | 59.80492 | -0.77832 | 0.23833 | 3.26575 | 0.00109 | 0.02548 | 103.758 | 50.46275 | 56.18183 | 28.81708 |
| 547 | Efcab7 | 47.3664 | 0.8373 | 0.25658 | -3.26333 | 0.0011 | 0.02561 | 51.26933 | 26.19956 | 65.07557 | 46.92114 |
| 548 | Ly86 | 2199.685 | 0.46005 | 0.14098 | -3.26316 | 0.0011 | 0.02561 | 2978.227 | 998.0467 | 3446.033 | 1376.434 |
| 549 | Kif15 | 108.8457 | -0.62171 | 0.19055 | 3.26273 | 0.0011 | 0.02561 | 151.0429 | 108.7944 | 105.4918 | 70.05397 |
| 550 | Map4 | 2082.962 | -0.31776 | 0.09743 | 3.2615 | 0.00111 | 0.02567 | 1967.007 | 2564.109 | 1745.424 | 2055.306 |
| 551 | Churc1 | 169.4689 | 0.5185 | 0.15902 | -3.26067 | 0.00111 | 0.02567 | 163.3257 | 119.6751 | 222.2058 | 172.6689 |
| 552 | Aicda | 21.354 | -1.88796 | 0.57902 | 3.26059 | 0.00111 | 0.02567 | 44.55057 | 14.35539 | 23.67694 | 2.833105 |
| 553 | Gm13456 | 72.53198 | 0.96576 | 0.29637 | -3.25863 | 0.00112 | 0.0258 | 63.91045 | 44.41482 | 94.65078 | 87.15185 |
| 554 | Ifi44 | 18.56629 | 1.67632 | 0.51517 | -3.25392 | 0.00114 | 0.02619 | 21.32183 | 3.596049 | 35.80772 | 13.53955 |
| 555 | Slc7a7 | 156.7341 | 0.69824 | 0.21467 | -3.25265 | 0.00114 | 0.02626 | 179.0853 | 72.03725 | 257.4944 | 118.3193 |
| 556 | Nr2c1 | 143.8631 | -0.50913 | 0.15668 | 3.24957 | 0.00116 | 0.0265 | 155.0443 | 169.4121 | 133.0519 | 117.944 |
| 557 | Ttc28 | 601.9041 | 0.84681 | 0.26068 | -3.24847 | 0.00116 | 0.02655 | 499.6586 | 332.478 | 969.3513 | 606.1284 |
| 558 | Ndc80 | 168.7029 | -0.83066 | 0.25576 | 3.24784 | 0.00116 | 0.02656 | 368.8772 | 81.73256 | 178.9828 | 45.21896 |
| 559 | Tgfbr2 | 6830.874 | -0.22879 | 0.07056 | 3.24262 | 0.00118 | 0.02701 | 6768.378 | 7475.502 | 6701.567 | 6378.047 |
| 560 | Hal | 12.15324 | 1.79614 | 0.55489 | -3.23692 | 0.00121 | 0.0275 | 2.68745 | 4.189735 | 25.17581 | 16.55998 |
| 561 | Ulk1 | 635.6929 | -0.27322 | 0.08443 | 3.2362 | 0.00121 | 0.02753 | 651.6154 | 708.8687 | 596.5677 | 585.7197 |
| 563 | Als2 | 724.8065 | -0.37123 | 0.11482 | 3.23299 | 0.00123 | 0.02779 | 513.7976 | 1107.266 | 423.0048 | 855.1572 |
| 564 | Cep170b | 512.1281 | -0.32577 | 0.10085 | 3.23011 | 0.00124 | 0.02802 | 244.5027 | 866.1716 | 248.5175 | 689.3205 |
| 565 | Trp53i11 | 3232.252 | 0.79651 | 0.24666 | -3.22914 | 0.00124 | 0.02807 | 5626.27 | 263.2213 | 6576.132 | 463.3857 |
| 566 | Zdhhc8 | 824.3833 | -0.27961 | 0.08667 | 3.22619 | 0.00125 | 0.02827 | 677.7239 | 1068.688 | 672.0384 | 879.0833 |
| 567 | Ggps1 | 413.9576 | -0.33831 | 0.10486 | 3.22611 | 0.00125 | 0.02827 | 437.8688 | 453.3658 | 406.7588 | 357.837 |
| 568 | Arhgap25 | 2108.637 | 0.41651 | 0.12919 | -3.22392 | 0.00126 | 0.02844 | 3183.75 | 936.523 | 3060.237 | 1254.037 |
| 569 | Tmc8 | 985.183 | 0.24907 | 0.07736 | -3.21947 | 0.00128 | 0.02883 | 864.6968 | 1003.104 | 880.8035 | 1192.128 |
| 570 | Jund | 6627.585 | 0.75901 | 0.23595 | -3.21684 | 0.0013 | 0.02905 | 10529.29 | 775.6271 | 13879.04 | 1326.384 |
| 571 | Myof | 2535.247 | -0.65838 | 0.20478 | 3.21503 | 0.0013 | 0.02918 | 1470.873 | 4410.167 | 1483.545 | 2776.403 |
| 572 | Golga1 | 429.1876 | -0.38597 | 0.12015 | 3.21232 | 0.00132 | 0.02941 | 362.3995 | 573.1752 | 344.2536 | 436.9222 |
| 573 | Pgm2l1 | 401.7297 | -0.51483 | 0.16039 | 3.20987 | 0.00133 | 0.02956 | 446.9504 | 465.9778 | 369.2306 | 324.76 |
| 574 | Il5ra | 480.4475 | 0.56781 | 0.17703 | -3.20734 | 0.00134 | 0.02977 | 698.1876 | 169.5841 | 800.8798 | 253.1386 |
| 575 | A530054K1 | 449.0213 | -0.40551 | 0.12653 | 3.20478 | 0.00135 | 0.02998 | 280.0286 | 689.0422 | 308.4481 | 518.5662 |
| 576 | Atrnl1 | 130.0224 | -0.50687 | 0.15824 | 3.20325 | 0.00136 | 0.03009 | 158.584 | 136.8975 | 129.1629 | 95.44537 |
| 577 | Selplg | 755.8432 | -0.35952 | 0.11227 | 3.20217 | 0.00136 | 0.03015 | 838.8635 | 696.6217 | 945.48 | 542.4076 |
| 578 | E430025E21 | 1260.985 | -0.35172 | 0.10998 | 3.1981 | 0.00138 | 0.03053 | 1208.969 | 1558.726 | 1055.718 | 1220.529 |
| 579 | Gpr176 | 22.86018 | 1.47569 | 0.46172 | -3.19609 | 0.00139 | 0.0306 | 8.081815 | 13.77057 | 27.7667 | 41.82162 |
| 580 | Abca1 | 1809.133 | -0.83787 | 0.26216 | 3.19606 | 0.00139 | 0.0306 | 3787.525 | 341.216 | 2919.725 | 188.0649 |
| 581 | Nuak2 | 707.8003 | 0.31168 | 0.09754 | -3.19526 | 0.0014 | 0.03062 | 1039.733 | 322.3767 | 1068.643 | 400.4488 |
| 582 | Wipf1 | 3763.179 | -0.31661 | 0.09922 | 3.19088 | 0.00142 | 0.03104 | 5214.521 | 3036.873 | 4364.038 | 2437.286 |
| 583 | Zfp874a | 142.1754 | -0.49213 | 0.15429 | 3.18957 | 0.00142 | 0.03113 | 119.5758 | 206.0751 | 97.07305 | 145.9777 |
| 584 | Fam102b | 293.7171 | -0.53264 | 0.16704 | 3.18873 | 0.00143 | 0.03113 | 407.1363 | 228.6739 | 381.4923 | 157.5658 |
| 585 | Tmsb4x | 25784.55 | 0.65178 | 0.20441 | -3.18856 | 0.00143 | 0.03113 | 29355.76 | 14655.81 | 35953.27 | 23173.36 |
| 586 | Ncapd2 | 718.7486 | -0.37559 | 0.11783 | 3.18765 | 0.00143 | 0.03113 | 1075.76 | 534.7309 | 852.3488 | 412.1547 |
| 587 | Add1 | 3445.289 | -0.29311 | 0.09195 | 3.18759 | 0.00143 | 0.03113 | 5694.56 | 1651.624 | 5087.562 | 1347.41 |
| 588 | Cog1 | 697.6128 | -0.25901 | 0.0814 | 3.18198 | 0.00146 | 0.03169 | 667.5015 | 774.2154 | 701.5904 | 647.1439 |
| 589 | Kcnj10 | 167.8527 | 1.61838 | 0.50919 | -3.17835 | 0.00148 | 0.03204 | 305.7707 | 5.464183 | 339.5532 | 20.62266 |
| 590 | Slc2a3 | 633.9378 | -0.32211 | 0.10148 | 3.17422 | 0.0015 | 0.03244 | 1098.55 | 474.3096 | 584.8259 | 378.0656 |

| | | | | | | | | | | | |
|---|---|---|---|---|---|---|---|---|---|---|---|
| 591 | Sik1 | 9747.495 | 0.65869 | 0.20783 | -3.16935 | 0.00153 | 0.03291 | 18263.17 | 1063.122 | 17973.82 | 1689.869 |
| 592 | Racgap1 | 465.3517 | 0.49007 | 0.15464 | -3.16908 | 0.00153 | 0.03291 | 289.1897 | 545.2872 | 258.5242 | 768.4056 |
| 593 | Mlkl | 35.28706 | 0.96038 | 0.30318 | -3.16767 | 0.00154 | 0.03302 | 25.1918 | 27.35816 | 34.0553 | 54.54298 |
| 594 | Fnip1 | 1447.345 | -0.46011 | 0.14528 | 3.16692 | 0.00154 | 0.03305 | 1353.239 | 1827.057 | 1284.273 | 1324.81 |
| 595 | AI504432 | 506.9052 | -0.85659 | 0.27072 | 3.16406 | 0.00156 | 0.03324 | 898.1899 | 463.5367 | 414.235 | 251.6592 |
| 596 | Pdzd2 | 260.8471 | 0.9282 | 0.29338 | -3.16384 | 0.00156 | 0.03324 | 388.8828 | 38.23931 | 541.9456 | 74.32068 |
| 597 | Inadl | 772.6755 | -0.36223 | 0.11469 | 3.15836 | 0.00159 | 0.03378 | 470.585 | 1197.154 | 493.5513 | 929.4113 |
| 598 | Pex1 | 590.231 | -0.33984 | 0.10761 | 3.15817 | 0.00159 | 0.03378 | 480.0425 | 796.8184 | 455.6018 | 628.4613 |
| 599 | Gm6254 | 31.80191 | 1.94715 | 0.61689 | -3.15642 | 0.0016 | 0.03393 | 28.91387 | 16.04275 | 6.353779 | 75.89724 |
| 600 | Unc93b1 | 7775.014 | 0.31232 | 0.09896 | -3.15588 | 0.0016 | 0.03393 | 12158.02 | 2988.737 | 12236.75 | 3716.551 |
| 601 | Srgn | 9817.364 | 0.58066 | 0.18407 | -3.15463 | 0.00161 | 0.03397 | 8613.593 | 8183.753 | 10177.04 | 12295.07 |
| 602 | Ccnb2 | 61.85617 | -0.8112 | 0.25715 | 3.1546 | 0.00161 | 0.03397 | 104.9114 | 47.25527 | 68.33626 | 26.92179 |
| 603 | Slc35d2 | 143.771 | 0.60896 | 0.1934 | -3.14875 | 0.00164 | 0.0346 | 181.7085 | 45.9159 | 276.1108 | 71.34867 |
| 604 | Nfkbia | 3778.379 | 0.42989 | 0.13655 | -3.14815 | 0.00164 | 0.03462 | 2947.442 | 3642.218 | 3608.45 | 4915.406 |
| 605 | Prkce | 1448.346 | -0.42681 | 0.13598 | 3.13882 | 0.0017 | 0.03559 | 2378.158 | 709.8262 | 2177.988 | 527.4124 |
| 606 | Cxcl10 | 19.97239 | 1.4969 | 0.47697 | -3.13836 | 0.0017 | 0.03559 | 25.485 | 7.83409 | 23.02786 | 23.5426 |
| 607 | Hmmr | 50.56217 | -0.92114 | 0.29351 | 3.13836 | 0.0017 | 0.03559 | 75.11385 | 53.09402 | 46.52421 | 27.51659 |
| 608 | Cks2 | 282.2876 | 0.47547 | 0.15151 | -3.13818 | 0.0017 | 0.03559 | 259.9349 | 249.6625 | 271.3054 | 348.2476 |
| 609 | Vcl | 383.4114 | -0.89961 | 0.28678 | 3.13696 | 0.00171 | 0.03568 | 405.7372 | 478.2549 | 397.9747 | 251.6789 |
| 610 | Rpl30 | 10677.9 | 0.60939 | 0.19431 | -3.13609 | 0.00171 | 0.03573 | 9383.582 | 7989.104 | 13085.04 | 12253.88 |
| 611 | Zfp456 | 186.5872 | -0.57917 | 0.18475 | 3.13491 | 0.00172 | 0.0358 | 149.4378 | 286.5362 | 119.3954 | 190.9796 |
| 612 | Msn | 21046.19 | -0.31045 | 0.09905 | 3.13429 | 0.00172 | 0.0358 | 24793.85 | 20401.68 | 22549.52 | 16439.7 |
| 613 | Gem | 2065.504 | 0.35317 | 0.11269 | -3.13411 | 0.00172 | 0.0358 | 2871.348 | 1172.625 | 2719.244 | 1498.801 |
| 614 | Sub1 | 5886.262 | 0.57411 | 0.18322 | -3.13351 | 0.00173 | 0.03581 | 5528.168 | 4479.864 | 6837.706 | 6699.31 |
| 615 | Got1 | 1662.016 | 0.32096 | 0.10257 | -3.12921 | 0.00175 | 0.03625 | 1502.054 | 1539.564 | 1681.909 | 1924.536 |
| 616 | Fam63b | 1338.343 | -0.44733 | 0.14298 | 3.12862 | 0.00176 | 0.03625 | 1525.143 | 1383.497 | 1432.193 | 1012.537 |
| 617 | Scml4 | 473.7565 | -0.63109 | 0.20172 | 3.12854 | 0.00176 | 0.03625 | 1203.435 | 80.52926 | 559.7032 | 51.35884 |
| 618 | Cpm | 942.2373 | 0.91721 | 0.29343 | -3.1258 | 0.00177 | 0.03653 | 1685.644 | 54.46839 | 1923.088 | 105.7492 |
| 619 | Sorl1 | 9613.393 | -0.56724 | 0.18152 | 3.12488 | 0.00178 | 0.03659 | 12471.24 | 10321.8 | 8725.044 | 6935.481 |
| 620 | Nr1d2 | 1379.798 | -0.2417 | 0.07742 | 3.12216 | 0.0018 | 0.03687 | 1303.321 | 1562.706 | 1333.793 | 1319.371 |
| 621 | 11-Sep | 3382.958 | -0.31657 | 0.10155 | 3.11756 | 0.00182 | 0.03739 | 1864.848 | 5511.079 | 1733.744 | 4422.16 |
| 622 | Rffl | 404.5147 | 0.30081 | 0.0965 | -3.11705 | 0.00183 | 0.0374 | 384.955 | 362.5163 | 424.2787 | 446.3089 |
| 623 | Itgb5 | 149.0396 | 1.45296 | 0.4662 | -3.11661 | 0.00183 | 0.0374 | 240.6205 | 8.458796 | 319.9017 | 27.17736 |
| 624 | Bcl6b | 32.3847 | 1.00497 | 0.32268 | -3.11442 | 0.00184 | 0.03759 | 13.68976 | 28.54484 | 28.58632 | 58.71788 |
| 625 | Zfp937 | 50.81691 | -0.85145 | 0.27341 | 3.11417 | 0.00184 | 0.03759 | 80.86818 | 50.72438 | 44.02574 | 27.64936 |
| 626 | Atp10a | 53.32596 | 0.70432 | 0.22622 | -3.11345 | 0.00185 | 0.03762 | 27.32339 | 57.97296 | 32.13361 | 95.87389 |
| 627 | Il12rb1 | 35.39892 | 1.10365 | 0.35454 | -3.11295 | 0.00185 | 0.03763 | 56.53677 | 9.606304 | 53.8445 | 21.60812 |
| 628 | 2900005J15 | 21.31077 | 1.02694 | 0.33042 | -3.108 | 0.00188 | 0.0382 | 3.313445 | 24.26319 | 7.319627 | 50.34682 |
| 629 | Ildr1 | 354.6368 | 0.54695 | 0.17604 | -3.10699 | 0.00189 | 0.03827 | 427.9351 | 257.5412 | 356.6897 | 376.3813 |
| 630 | Rbpjl | 20.1859 | 1.59696 | 0.51407 | -3.10649 | 0.00189 | 0.03828 | 27.11618 | 3.23297 | 38.41639 | 11.97805 |
| 631 | Kif18b | 43.33676 | -1.03397 | 0.3329 | 3.10594 | 0.0019 | 0.03829 | 67.67498 | 46.27344 | 37.25081 | 22.14781 |
| 632 | Elmsan1 | 5527.338 | -0.58527 | 0.18851 | 3.10464 | 0.00191 | 0.0384 | 5560.596 | 7141.733 | 4670.202 | 4736.822 |
| 633 | Igkv16-104 | 391.3775 | 0.59071 | 0.19068 | -3.09783 | 0.00195 | 0.03923 | 515.8967 | 129.2446 | 725.2897 | 195.0791 |
| 634 | Zfp365 | 38.54705 | 0.90548 | 0.29265 | -3.09406 | 0.00197 | 0.03962 | 32.47455 | 18.34997 | 67.55519 | 35.80849 |
| 635 | Vma21 | 980.1361 | 0.33419 | 0.10801 | -3.09397 | 0.00197 | 0.03962 | 691.3661 | 1081.243 | 785.2865 | 1362.649 |
| 636 | Bspry | 289.1905 | 0.64122 | 0.20735 | -3.09248 | 0.00198 | 0.03973 | 70.74028 | 367.2504 | 142.7751 | 575.9961 |
| 637 | Mrpl52 | 554.3062 | 0.7737 | 0.25021 | -3.09225 | 0.00199 | 0.03973 | 329.9316 | 508.33 | 500.2689 | 878.6945 |
| 638 | Dusp1 | 2899.659 | 1.6151 | 0.52255 | -3.0908 | 0.002 | 0.03987 | 5308.08 | 61.71213 | 6006.617 | 222.2278 |
| 639 | Tmem88 | 22.23075 | 1.36244 | 0.44123 | -3.0878 | 0.00202 | 0.04021 | 30.94656 | 7.424922 | 29.52433 | 21.02718 |
| 640 | Il10rb | 1151.206 | 0.25119 | 0.08138 | -3.08663 | 0.00202 | 0.0403 | 756.3454 | 1311.169 | 977.3763 | 1559.932 |
| 641 | Utrn | 931.2353 | -0.89944 | 0.29149 | 3.08572 | 0.00203 | 0.04037 | 1485.646 | 537.6058 | 1419.101 | 282.589 |
| 642 | Ddx24 | 3322.49 | 0.30933 | 0.10027 | -3.08484 | 0.00204 | 0.04041 | 2887.242 | 3145.308 | 3356.711 | 3900.698 |
| 643 | Vps33a | 1072.725 | -0.20913 | 0.0678 | 3.08448 | 0.00204 | 0.04041 | 832.9736 | 1441.391 | 770.5383 | 1245.998 |
| 644 | Tpx2 | 108.5158 | -0.60866 | 0.19751 | 3.08167 | 0.00206 | 0.04073 | 142.1988 | 114.3387 | 102.7138 | 74.81203 |
| 645 | Eif3j2 | 155.2338 | 1.41649 | 0.45986 | -3.08026 | 0.00207 | 0.04087 | 27.48221 | 129.2264 | 92.05556 | 372.1712 |
| 646 | Zfp36 | 8909.775 | 0.37853 | 0.12318 | -3.07297 | 0.00212 | 0.04181 | 13394.86 | 3568.009 | 14030.48 | 4645.752 |
| 647 | 4833420G1 | 1044.804 | -0.25251 | 0.08227 | 3.06924 | 0.00215 | 0.04228 | 1444.271 | 799.9158 | 1262.679 | 672.3495 |
| 648 | Slc35a5 | 272.2151 | -0.40505 | 0.1321 | 3.06634 | 0.00217 | 0.04262 | 253.1678 | 351.2071 | 220.527 | 263.9585 |
| 649 | Tmem184b | 921.7393 | 0.41569 | 0.13564 | -3.06475 | 0.00218 | 0.04279 | 779.481 | 881.4171 | 848.3845 | 1177.675 |
| 650 | B4galnt1 | 2398.634 | 0.25908 | 0.08459 | -3.06272 | 0.00219 | 0.04301 | 3613.036 | 1538.328 | 2601.984 | 1841.19 |
| 651 | Clec12a | 226.9592 | 0.74467 | 0.24342 | -3.05926 | 0.00222 | 0.04345 | 354.625 | 33.44039 | 462.1141 | 57.65742 |
| 653 | Ndnf | 8.943662 | 1.57473 | 0.51535 | -3.05563 | 0.00225 | 0.04391 | 5.491421 | 4.757958 | 9.67187 | 15.8534 |
| 654 | Rbpj | 2579.585 | 0.33015 | 0.10822 | -3.05085 | 0.00228 | 0.04455 | 1067.99 | 3568.922 | 1191.196 | 4490.23 |
| 655 | Map3k12 | 53.99116 | 0.86002 | 0.28215 | -3.04808 | 0.0023 | 0.04489 | 69.3908 | 28.43743 | 65.28112 | 52.85528 |
| 656 | Cebpb | 112.9868 | 1.13933 | 0.37387 | -3.04743 | 0.00231 | 0.04492 | 118.5861 | 49.00419 | 172.3781 | 111.9787 |
| 657 | Unc119 | 424.7587 | 0.4323 | 0.14207 | -3.04288 | 0.00234 | 0.04554 | 623.9258 | 154.8655 | 710.7279 | 209.5155 |
| 658 | Slfn2 | 3952.384 | 0.70048 | 0.23027 | -3.04194 | 0.00235 | 0.04561 | 1032.116 | 5024.354 | 1517.577 | 8235.488 |
| 660 | Skap2 | 498.6052 | -0.33013 | 0.10876 | 3.03527 | 0.0024 | 0.04648 | 691.7631 | 367.1015 | 642.6227 | 292.9335 |
| 661 | Scand1 | 177.0886 | 1.26167 | 0.41572 | -3.03492 | 0.00241 | 0.04648 | 139.3199 | 82.67102 | 276.9476 | 209.416 |
| 663 | Cast | 1611.655 | -0.29984 | 0.09883 | 3.0339 | 0.00241 | 0.04656 | 1972.585 | 1464.364 | 1821.855 | 1187.817 |
| 664 | Hes1 | 364.1813 | 0.84326 | 0.27798 | -3.03351 | 0.00242 | 0.04656 | 712.2782 | 50.54761 | 601.0691 | 92.83048 |
| 665 | Kansl1l | 345.3657 | -0.64189 | 0.21164 | 3.03296 | 0.00242 | 0.04656 | 457.2539 | 315.7153 | 408.155 | 200.3387 |
| 666 | Fos | 6851.699 | 1.9375 | 0.63888 | -3.03265 | 0.00242 | 0.04656 | 14467.07 | 52.06747 | 12585.13 | 302.5247 |

| | | | | | | | | | | | |
|---|---|---|---|---|---|---|---|---|---|---|---|
| 667 | Havcr1 | 182.7161 | 0.83872 | 0.27665 | -3.03166 | 0.00243 | 0.04664 | 37.76371 | 228.8459 | 48.41615 | 415.8386 |
| 668 | Kctd9 | 517.0797 | -0.28064 | 0.09264 | 3.02926 | 0.00245 | 0.04694 | 374.8776 | 741.7575 | 341.771 | 609.9128 |
| 669 | Gm11707 | 14.89981 | 1.72727 | 0.57029 | -3.02876 | 0.00246 | 0.04695 | 18.48144 | 2.081027 | 29.82224 | 9.214522 |
| 670 | Tap2 | 1614.693 | 0.35323 | 0.11669 | -3.02702 | 0.00247 | 0.04715 | 1764.216 | 1208.393 | 1939.769 | 1546.396 |
| 671 | Nfkbid | 5010.979 | 0.45788 | 0.15133 | -3.02572 | 0.00248 | 0.04729 | 5348.132 | 3703.101 | 5893.878 | 5098.804 |
| 672 | Wars2 | 354.6344 | -0.36751 | 0.12155 | 3.02362 | 0.0025 | 0.04755 | 301.4921 | 484.2326 | 257.1941 | 375.6187 |
| 673 | Tmem17 | 44.40365 | 0.83633 | 0.27667 | -3.02285 | 0.0025 | 0.0476 | 29.28418 | 35.1067 | 50.09752 | 63.12618 |
| 674 | Rgl1 | 620.1177 | 0.43073 | 0.14267 | -3.01911 | 0.00254 | 0.04812 | 748.6018 | 187.4552 | 1290.335 | 254.0784 |
| 675 | Tcf12 | 2698.742 | -0.35189 | 0.11657 | 3.01863 | 0.00254 | 0.04813 | 2246.885 | 3533.584 | 2249.866 | 2764.632 |
| 676 | Ssbp3 | 251.9325 | -0.45058 | 0.14931 | 3.01772 | 0.00255 | 0.0482 | 402.6683 | 135.1795 | 371.0655 | 98.81672 |
| 677 | Cd93 | 1074.794 | -0.6305 | 0.20897 | 3.01714 | 0.00255 | 0.04822 | 1328.526 | 1144.885 | 1090.997 | 734.7698 |
| 678 | Tmem104 | 327.0485 | 0.39396 | 0.13065 | -3.01527 | 0.00257 | 0.04838 | 285.8292 | 313.7861 | 296.37 | 412.2085 |
| 679 | Aspm | 98.74662 | -0.59713 | 0.19816 | 3.01343 | 0.00258 | 0.04861 | 159.2269 | 83.38697 | 97.94714 | 54.42541 |
| 680 | Gen1 | 62.74261 | -0.6809 | 0.22613 | 3.01114 | 0.0026 | 0.0489 | 94.90995 | 55.2322 | 66.99852 | 33.82976 |
| 681 | Rasgef1b | 11997.18 | -0.22263 | 0.07401 | 3.0081 | 0.00263 | 0.04932 | 12896.55 | 13378.36 | 10253.11 | 11460.7 |
| 682 | Nnt | 495.1172 | 0.34869 | 0.11595 | -3.0072 | 0.00264 | 0.04938 | 486.7926 | 350.8205 | 694.8474 | 448.0081 |
| 683 | Smg6 | 1776.742 | -0.30839 | 0.10256 | 3.00685 | 0.00264 | 0.04938 | 1689.823 | 2150.501 | 1531.564 | 1735.079 |
| 684 | Tspan2 | 195.3572 | -0.62752 | 0.2088 | 3.00544 | 0.00265 | 0.04953 | 363.947 | 64.90896 | 311.1257 | 41.44715 |
| 685 | Cmtm6 | 2952.547 | 0.20828 | 0.06931 | -3.00508 | 0.00266 | 0.04953 | 3426.687 | 2393.435 | 3225.293 | 2764.775 |
